# Longitudinal Urine Metabolic Profiling and Gestational Age Prediction in Pregnancy

**DOI:** 10.1101/2022.07.10.499478

**Authors:** Songjie Chen, Xiaotao Shen, Liang Liang, Monika Avina, Hanyah Zackriah, Laura Jelliffe-Pawlowski, Larry Rand, Michael Snyder

## Abstract

Pregnancy is a critical time that has long-term impacts on both maternal and fetal health. During pregnancy, the maternal metabolome undergoes dramatic systemic changes, although correlating longitudinal changes in maternal urine remain largely unexplored. We applied an LCMS-based untargeted metabolomics profiling approach to analyze 346 longitudinal maternal urine samples collected throughout pregnancy for 36 women from diverse ethnic backgrounds with differing clinical characteristics. We detected 20,314 metabolic peaks and annotated 875 metabolites. Altered metabolites include a broad panel of glucocorticoids, lipids, and amino acid derivatives, which revealed systematic pathway alterations during pregnancy. We also developed a machine-learning model to precisely predict gestational age (GA) at time of sampling using urine metabolites that provides a non-invasive method for pregnancy dating. This longitudinal maternal urine study demonstrates the clinical utility of using untargeted metabolomics in obstetric settings.

**One Sentence Summary:** Machine-learning based gestational age and due date using longitudinal urine samples of pregnancy.

## INTRODUCTION

The accurate dating of GA provides essential guidelines for the prenatal medical care. Current GA dating approaches based on the last menstrual period (LMP) are problematic given imprecise recollection of dates and symptoms like breakthrough bleeding in early pregnancy, which may be mistaken for a period (*1–3*). Fetal ultrasound is the most precise current measure of GA, but is limited by both timing and access to resources (*4–8*). GA dating is more accurate the earlier an ultrasound is performed, and optimal pregnancy dating can be achieved prior to 20-weeks (*9–11*). However, this also requires both sophisticated equipment and well-trained sonographers (*6,8,12–14*). Thus, more affordable, accessible, and accurate GA dating methodology represents an unmet clinical need, particularly for pregnant women across diverse socio-economic backgrounds.

Recent developments in omics profiling technology provide new possibilities for characterization of both normal and high-risk pregnancies. Pregnancy is a highly dynamic programmed process that induces a broad spectrum of changes in the maternal transcriptome, proteome, and metabolome (*15,19*). Notably, the metabolome, as the direct outcome of diverse biochemical reactions, is a highly sensitive readout of metabolic regulation during pregnancy (*20–21*). Investigation of longitudinal maternal metabolomic alternations over the course of pregnancy has the potential to be a highly informative approach for mechanistic investigation and a breakthrough tool for GA dating. This approach has recently attracted more attention (*21,33,34*) but has relied mostly on maternal blood samples (*21,35–37*). Use of maternal urine for GA dating and metabolic profiling has yet to be explored and may provide a cost effective and non-invasive method that could be easily translated into clinical settings. If found to be useful, it would transform the prenatal care, especially in under-resourced regions.

Thus far, pregnancy-related metabolic research has focused primarily on identifying biomarkers as indicators of risk for adverse pregnancy outcomes like preeclampsia, preterm birth, and gestational diabetes (*22–29*). Emerging data suggests, however, that increased surveillance of metabolomic changes during pregnancy may also provide improved opportunities for understanding of maternal metabolic alterations at differential gestational ages (GA), stratify risk-based biomarkers by clinical benefit, and to better elucidate the pregnancy process (*21,30–32*).

In this study, we profiled longitudinal urine samples collected from 36 pregnant women receiving prenatal care in public and private clinics in San Francisco and annotated a large number of urine metabolites correlated with pregnancy progression. Using this data, we investigated potential alterations in maternal metabolic pathways throughout pregnancy and used this data to predict GA. Although the pregnancy induced metabolome alternations are highly personalized, we were able to identify such individualized alternations.

## RESULTS

### Study Design of the SMART-D Pregnancy Cohort

This observational study aimed to assess whether the urine metabolome in pregnancy could be used to identify dynamic metabolic changes during pregnancy and predict GA by week. To do this, we used urine samples collected from 36 pregnant women receiving prenatal care in San Francisco who were recruited into the SMART Diaphragm (SMART-D) study between November 2014 and October 2018 (**Fig. 1a**). The SMART-D study developed and iterated a patient-controlled, vaginally inserted device that detects microscopic changes in cervical collagen structure to provide earlier predictions of preterm birth risk and open a new potential treatment window. Diverse samples were collected during longitudinal visits over the course of pregnancy and postpartum periods, including urine samples, cervical-vaginal swabs, etc., together with detailed clinical and device measurement data. Urine samples used for analyses in the present study were collected as part of the SMART-D study protocol wherein at least one urine sample from each participant was collected for each trimester. Each participant contributed 3 - 13 samples throughout pregnancy; overall, each week of pregnancy after 15 weeks was represented by at least one sample across participants (**Fig. 1a**, **Methods**). High-resolution liquid chromatography-mass spectrometry (LC-MS) was used to characterize the metabolome of all collected urine samples (**Methods**).

**Figure 1.**
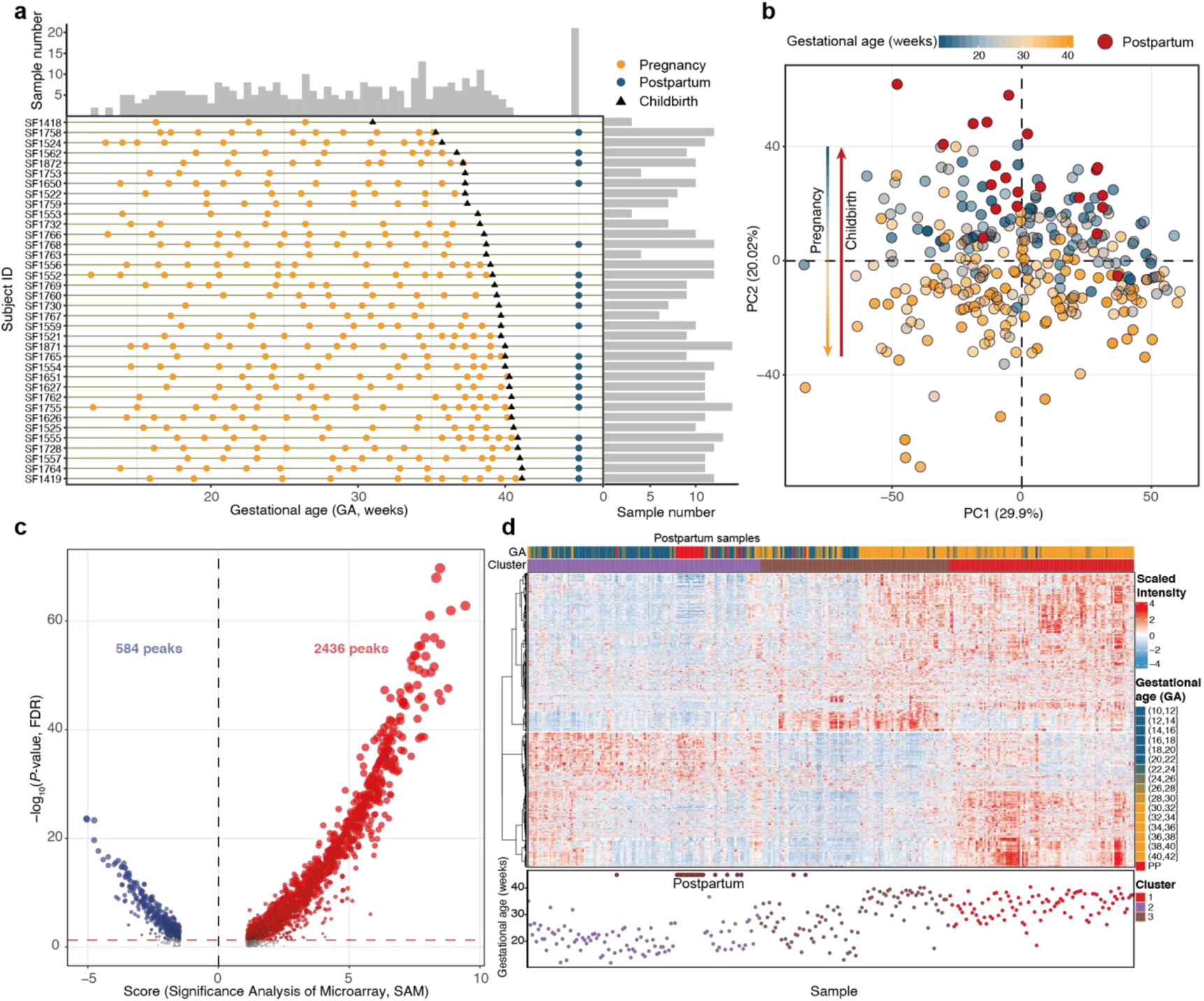
Overview of urine collected as part of the Smart-D study. (**a**) The sampling time points for individual participants. Each row represents an individual participant. The histogram and bar on the top and the right show the number of samples collected at each gestational age range (bin width = 0.5 weeks) and from each individual participant, respectively. Orange dots represent samples taken during pregnancy, blue dots represent samples taken after childbirth, and black triangles represent childbirth. (**b**) Principal Component Analysis (PCA) distributed individual urine samples according to gestational age (based on metabolic peaks with QC RSD < 30%). The two PCs explaining the largest part of the variation are shown. (**c**) The volcano plot shows the altered metabolic peaks during pregnancy, using the linear regression model (FDR adjusted *P*-value < 0.05) and SAM test (FDR adjusted *P*-value < 0.05). Red dots represent metabolic features that increased during pregnancy and blue dots represent features that decreased during pregnancy. Red dots represent increased metabolic peaks according to pregnancy and blue dots represent decreased metabolic peaks according to pregnancy. (**d**) Unsupervised k-means consensus-clustering analysis of 3,020 altered metabolic features clustered into three groups. Postpartum samples (represented by red in the GA legend) were clustered with early pregnancy samples.

Participants of diverse backgrounds were included in the SMART-D study from which these samples are derived (**Fig. S1a**). The 36 participants in the cohort were comprised of five ethnicities (Asian, Black, Latina, Pacific Islander, and White), with ages ranging from 21 to 39 years old. The pre-pregnancy body mass index (BMI) of participants varied from 19.5 to 57.2. Parity ranged from 1 to 9. Detailed characteristics for all participants are provided in **Table S1**. The high-density sampling design of the SMART-D study enabled the close monitoring of dynamic metabolome alterations of women throughout pregnancy.

**Table 1.**
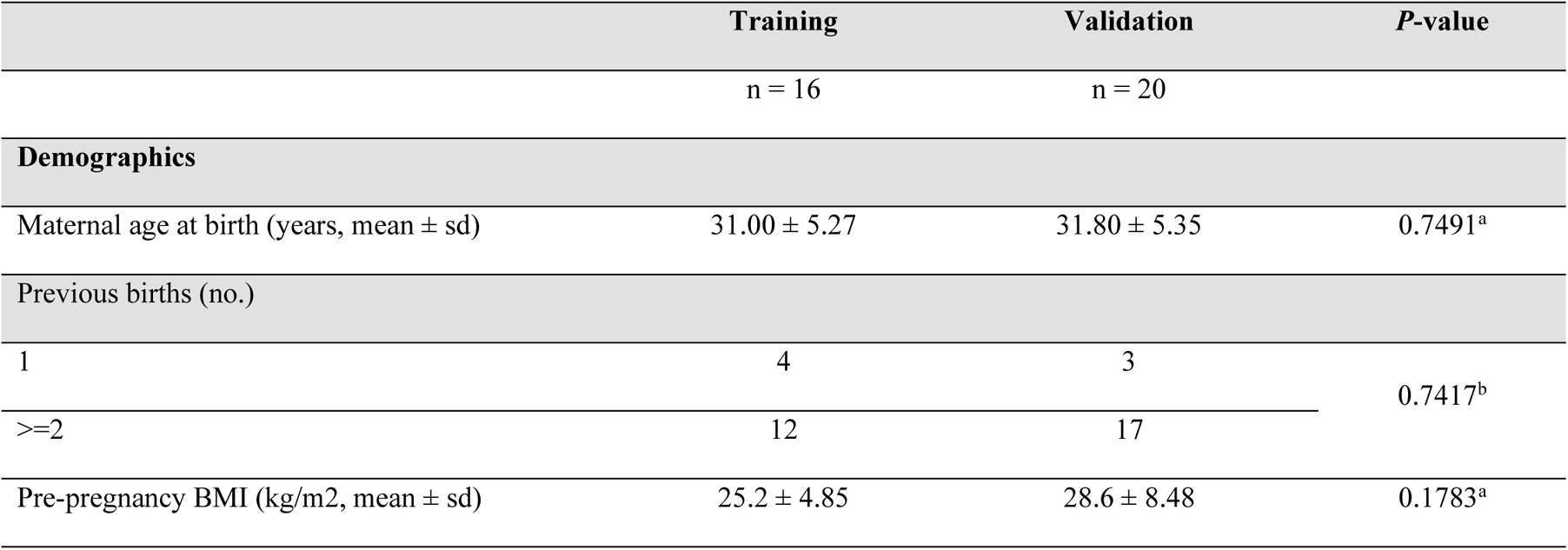
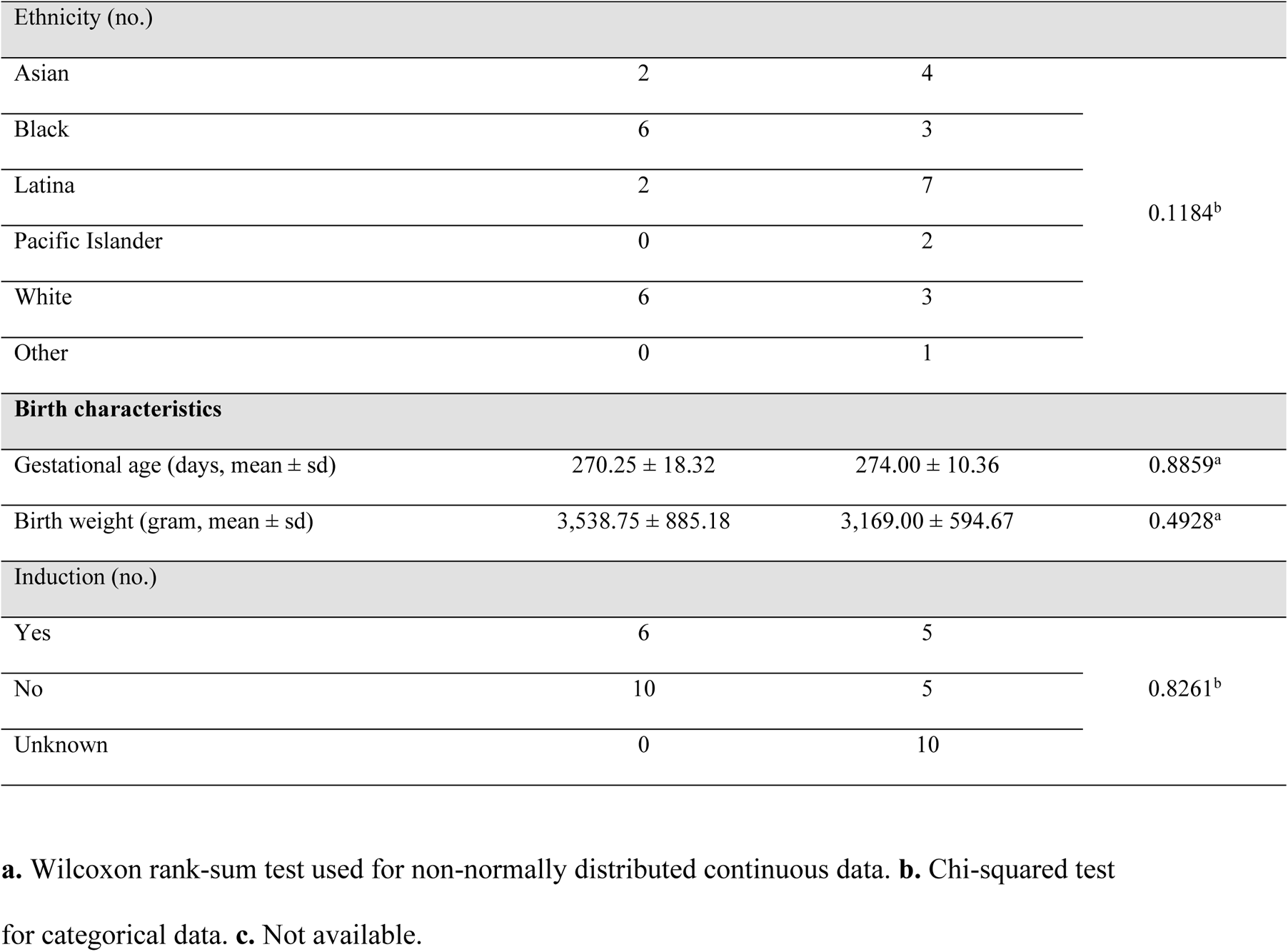
Demographics and birth characteristics of the training and validation dataset.

|  | Training | Validation | *P*-value |
| --- | --- | --- | --- |
|  | n = 16 | n = 20 |  |
| **Demographics** |  |  |  |
| Maternal age at birth (years, mean ± sd) | 31.00 ± 5.27 | 31.80 ± 5.35 | 0.7491<sup>a</sup> |
| Previous births (no.) |  |  |  |
| 1 | 4 | 3 | 0.7417<sup>b</sup> |
| $\geq 2$ | 12 | 17 | |
| Pre-pregnancy BMI (kg/m<sup>2</sup>, mean ± sd) | 25.2 ± 4.85 | 28.6 ± 8.48 | 0.1783<sup>a</sup> |

| Ethnicity (no.) |  |  |  |
| --- | --- | --- | --- |
| Asian | 2 | 4 | 0.1184<sup>b</sup> |
| Black | 6 | 3 |  |
| Latina | 2 | 7 |  |
| Pacific Islander | 0 | 2 |  |
| White | 6 | 3 |  |
| Other | 0 | 1 |  |
| Birth characteristics |  |  |  |
| Gestational age (days, mean ± sd) | 270.25 ± 18.32 | 274.00 ± 10.36 | 0.8859<sup>a</sup> |
| Birth weight (gram, mean ± sd) | 3,538.75 ± 885.18 | 3,169.00 ± 594.67 | 0.4928<sup>a</sup> |
| Induction (no.) |  |  |  |
| Yes | 6 | 5 | 0.8261<sup>b</sup> |
| No | 10 | 5 |  |
| Unknown | 0 | 10 |  |

**a.** Wilcoxon rank-sum test used for non-normally distributed continuous data. **b.** Chi-squared test for categorical data. **c.** Not available.

### The Urine Metabolome Accurately Reflects Metabolic Alterations During Pregnancy

Untargeted high-resolution metabolomics was performed on all collected urine samples. After data processing (peak detection and alignment) and cleaning (missing value processing, normalization and batch integration, outlier removal), we detected 20,314 metabolic peaks (or metabolic features, characterized by unique accurate mass and retention time) including 15,398 and 4,916 metabolic peaks in positive and negative modes, respectively. Forty-four samples were removed as outliers; 302 samples remained for all subsequent analyses (**Fig. S1b**, **Methods**). Quality of urine metabolomics data was assessed using Principal Component Analysis (PCA), which showed no batch effect. Additionally, most QC samples clustered tightly in the center among samples in positive, negative, and combined datasets (**Fig. S2**), indicating the high quality of our acquired metabolomics dataset. In addition, PCA including all metabolic peaks with QC RSD < 30% revealed a continuous separation between samples from early and later GA (**Fig. 1b**). Interestingly, the postpartum urine samples most closely resemble early GA urine samples (**Fig. 1b**). Additionally, most individual participants followed the same patterns of metabolic change as the overall dataset. (**Fig. S3**).

Overall metabolome alterations during pregnancy were then examined. Significance analysis for microarrays (SAM) and linear regression model (acquisition batch, BMI, mother age, parity, and ethnicity were adjusted as confounders) were utilized to discover altered metabolic peaks during pregnancy (SAM FDR < 0.05 and linear regression model FDR < 0.05, **Methods**). 14.87% of all detected metabolic peaks (3,020 out of 20,314 peaks, with 2,436 and 584 metabolic peaks in positive and negative modes, respectively) were significantly altered during pregnancy (**Fig. 1c**). Altered metabolic peaks were then used for unsupervised k-means consensus-clustering (**Fig. S4**, **Method**). Three robust clusters clearly correlated with gestational age were detected, namely cluster 1: 10-26 weeks, cluster 2: 26-32 weeks, and cluster 3: 32-42 weeks (**Fig. 1d**). Additionally, consistent with results from PCA, almost all samples after childbirth were included in cluster 1, which contains most of the early GA samples, suggesting that the urine metabolome rapidly returned to baseline and reflected early GA patterning. Taken together, these results demonstrated that a high-quality urine metabolome accurately reflects systemic metabolic alterations throughout pregnancy.

### Functional Metabolic Network and Alteration of Pathways During Pregnancy

An important strength of this study is the high-density urine sampling, which offered a more detailed view of the altered metabolic regulation at each stage of pregnancy. Because sampling times varied between participants, samples were assigned to 14 GA ranges ((10, 16] (baseline), (16, 18], (18, 20], (20, 22], (22, 24], (24, 26], (26, 28], (28, 30], (30, 32], (32, 34], (34, 36], (36, 38], (38, 42] and [PP) (postpartum)). To ensure the robustness and reproducibility of the statistical analysis, each GA range included at least 10 subjects and 10 samples. Altered metabolic peaks were identified using the Wilcoxon signed-rank test (FDR adjusted P-value < 0.05) for each GA range compared to the baseline GA range (10-16] (**Fig. 2b**). In order to assess the robustness of the identified altered metabolic peaks, we assessed whether peaks remained significantly altered in each of the subsequent GA ranges (except the PP range) and found that 84.83% of the identified peaks remained significantly altered in each of the GA ranges, suggesting a predictable overarching pattern of changes in metabolic features across pregnancy (**Fig. S5**).

**Figure 2.**
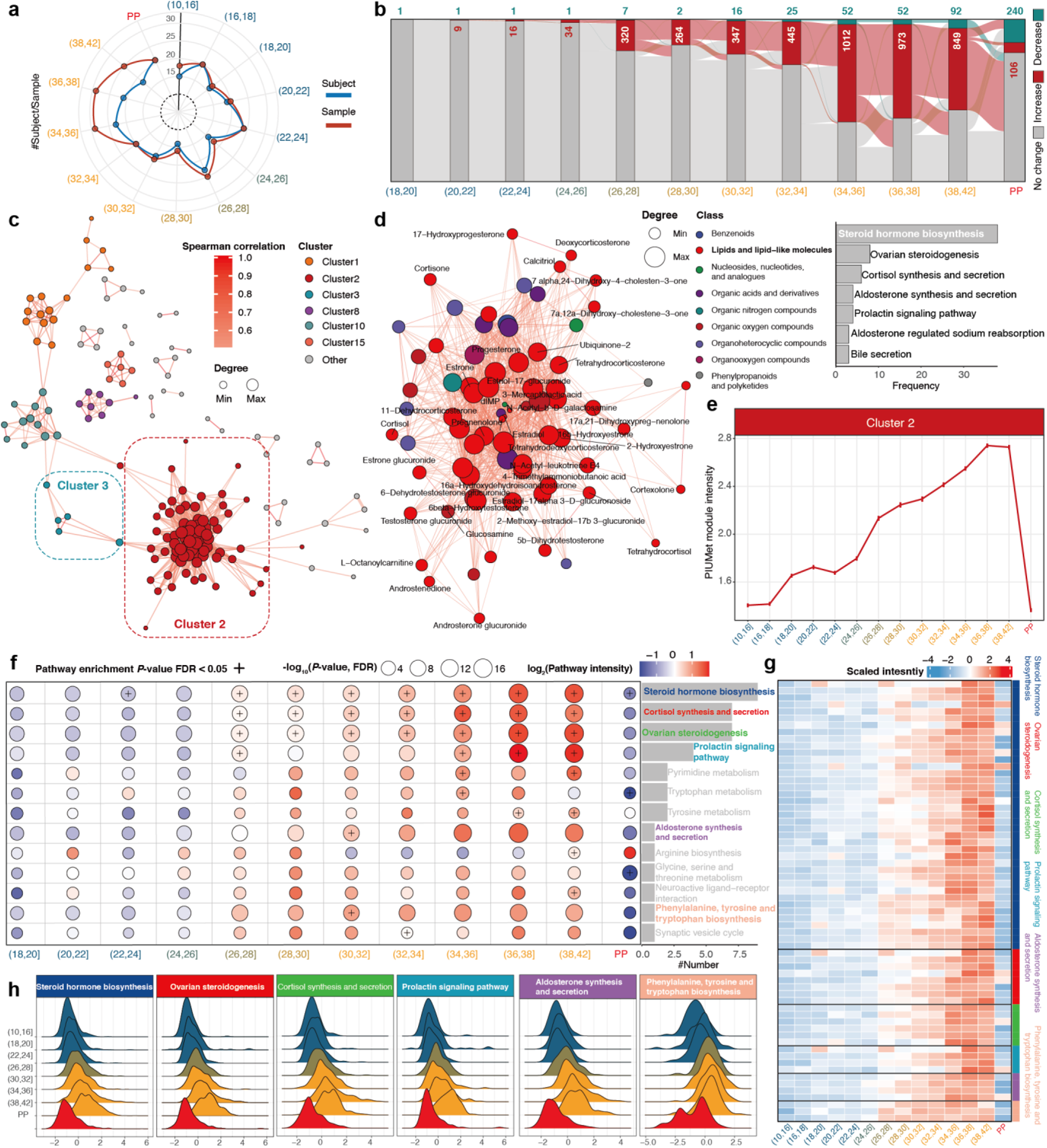
The dynamic changes of the urine metabolome during pregnancy. (**a**) Subject and sample numbers in each GA range. (**b**) The altered metabolic peaks in each GA range compared to baseline (10-16 weeks). The number of decreased and increased metabolic peaks are shown (decreases in green, increases in red). (**c**) The correlation network utilized the annotated metabolic peaks from PIUMet. Each color represents clusters identified using community analysis. (**d**) Cluster 2 which is identified utilized community analysis and changes are consistent in the cluster and at metabolite level (**Fig. S6**). The frequency of pathways of each metabolite is shown on the right panel. (**e**) The change of cluster 2 during pregnancy. The dots represent the mean values of cluster 2 in different GA ranges and the bars represent the standard error of the mean (SEM). (**f**) Altered pathways during pregnancy in each GA range. (**g**) This heatmap shows the changes of 6 metabolic pathways in metabolite (metabolic peak) level. (**h**) Ridgeline plot shows the changes of 6 metabolic pathways in pathway level.

A number of altered metabolic peaks were found to be clearly increased from early to late GA range during pregnancy (**Fig. 2b**), which is consistent with the PCA (**Fig. 1b**) and k-means consensus-clustering results (**Fig. 1d**). After childbirth, the number of altered metabolic peaks dramatically decreased compared to baseline. Based on the number of altered metabolic signatures of maternal urine, it was determined that pregnancy metabolic signals could be divided into four distinct periods, namely 10-18 weeks (no altered metabolic peaks), 18-26 weeks (altered metabolic peaks: 1-34), 26-34 weeks (altered metabolic peaks: 254-445) and 34-42 weeks (altered metabolic peaks: 849-1,012). These results matched the clusters from the unsupervised consensus-clustering described above (**Fig. 1d**).

In order to explore whether specific metabolic networks (or modules) were altered at different GA ranges over the pregnancy process, altered metabolic peaks were processed using PIUMet (*54*) and a network of altered metabolites was identified for each GA range. All annotated metabolites were then used to build a cross-sectional correlation network based on their Spearman correlations across our cohort (**Fig. 2c** and **Method**). In this correlation network, nodes (N) correspond to annotated metabolic peaks and edges (E). Two nodes being connected means that the correlation between those nodes is significant (FDR adjusted *P*-value < 0.05) and strong (absolute Spearman correlation > 0.5). The resulting cross-sectional correlation network contained 160 nodes and 1,148 edges. Interestingly, 80.4% of annotated metabolites from PIUMet (160 of 199) appeared in the correlation network. The topology of the network suggests that dense interactions occur between altered metabolites. Although metabolite levels change dynamically throughout pregnancy (**Fig. 1d**), it appeared clear that these changes are highly coordinated by a regulatory network.

We also examined the main clusters (or subnetworks) of related measurements from the correlation network using the community analysis method based on edge betweenness centrality, an unsupervised approach that iteratively prunes the network (removing the edges with the highest betweenness centrality) to reveal densely interconnected subgroups (clusters). From this, we detected 20 clusters with a modularity of 0.30. Seven clusters with at least 5 nodes were selected for subsequent analysis (**Fig. 2c** and **Fig. S6a**). Compared to the original correlation network, the 7 clusters retained 76.25% of nodes and 96.95% of edges remaining after edge punning, suggesting that the 7 clusters represented most of the correlations in the original network. These clusters may represent physiologically-related and -correlated metabolites during pregnancy.

We assessed the alterations of the 7 clusters during pregnancy by cluster and by the intensity of metabolic features, respectively (**Fig. S6b, c**). Only clusters 2 and 3 showed consistent changes during pregnancy at both cluster and metabolite level. Cluster 2 is the largest cluster (75 nodes and 963 edges) and contains lipids and lipid-like molecules (51 of 75, **Fig. 2d**), suggesting this cluster, including many steroids, represents a lipid-related metabolic regulation network during pregnancy. To investigate the physiological functions of this cluster, we then calculated the frequency of pathways to which all metabolites in cluster 2 belong. We found the top seven pathways were steroid hormone biosynthesis, ovarian steroidogenesis, cortisol synthesis and secretion, aldosterone synthesis and secretion, the prolactin signaling pathway, and aldosterone-regulated sodium reabsorption and bile secretion (right panel in **Fig. 2d**). Intriguingly, levels of cluster 2 metabolites increase throughout pregnancy (**Fig. 2e**) and exhibited a rapid increase in week 18 and week 26, which is consistent with the periods defined in **Fig. 1d** and **Fig. 2b**. Cluster 3 metabolites follow similar trends as cluster 2 metabolites during pregnancy (**Fig. S6b, c**). Cluster 3 contains 5 metabolites (3-methylguanine, 7-methylguanine, L-phenylalanine, asymmetric dimethylarginine, and (S)-3-hydroxy-N-methylcoclaurine) and no single pathway mapped to more than 1 metabolite (**Fig. S6c**). However, 4 out of 5 are amino acid modification metabolites, suggesting cluster 3 was related to the amino acid metabolism.

To further explore the altered metabolic pathways of pregnancy, pathway enrichment analysis was done using PIUMet for each GA range (based on the KEGG database). Thirteen pathways were selectively enriched for at least one GA range - most of which increased during pregnancy (FDR adjusted *P*-value < 0.05 and overlap ≥ 3, **Fig. 2f**). All 13 pathways were assessed and it was found that 6 were consistent at the metabolite and pathway level (**Fig. 2g, h**), namely steroid hormone biosynthesis, cortisol synthesis and secretion, ovarian steroidogenesis, the prolactin signaling pathway, aldosterone synthesis and secretion, and phenylalanine, tyrosine, and tryptophan biosynthesis. Five out of 6 pathways overlapped with the pathways in the regulated network (cluster 2) as described above.

### Gestational Age Prediction Using the Urine Metabolome Model

An accurate and noninvasive method of estimating gestational age has the potential to inform prenatal and neonatal care in instances where dating is uncertain. To this end, we examined whether the urine metabolome could be used to estimate gestational age. Urine samples during pregnancy were assigned to training (16 subjects, 125 samples) validation (20 subjects, 156 samples) datasets by acquisition batch (**Fig. S1b**). The demographics and birth characteristics of training and validation datasets were not significantly different (*P*-value > 0.05, **Table S1**).

The prediction model was constructed by starting with all metabolic peaks for feature selection based on the Boruta algorithm (see **Methods**). We then removed the metabolic peaks without acceptable peak shapes, with the remaining 28 metabolic peaks as potential biomarkers, and used these biomarkers to build a Random Forest prediction model (**Methods**, **Table S2, Fig. S7a**). The training dataset was utilized as the internal dataset and to validate prediction accuracy using the bootstrap method (1,000 times; see **Methods**). The root mean squared error (RMSE) between actual and predicted gestational age (weeks) was found to be 2.35 weeks and adjusted R^2^ was 0.86 (Pearson correlation r = 0.93; *P*-value < 2.2×10^-6^) (**Fig. S7b**). For the external validation dataset, the RMSE was 2.66 weeks and the adjusted R^2^ was 0.79 (Pearson correlation r = 0.89; *P*-value < 2.2×10^-6^) (**Fig. S7c**). This result demonstrated that the prediction model is not overfitting. Overall, our results demonstrated that the urine metabolome may be useful for accurately predicting gestational age.

The impact of patient demographics on prediction accuracy was also assessed. Maternal BMI, age, parity, and ethnicity were included with 28 metabolic peaks to construct a prediction model. The RMSE of this model was 2.70 and adjusted R^2^ was 0.76, which demonstrated no significant differences compared to the prediction model utilizing 28 metabolic peaks. Inclusion of subject demographics minimally improved prediction accuracy.

### Prediction of Gestational Age at the Individual Level

We demonstrated that the pregnancy urine metabolome could accurately predict gestational age with 28 metabolic peaks using a Random Forest model. Metabolic peaks were annotated using the in-house MS^2^ pipeline based on in-house and public MS^2^ databases (**Methods**). While 875 of 20,314 total level 1 or level 2 metabolites were annotated in the full dataset, only 5 out of the 28 metabolic peaks in our final model were annotated (**Table S2**). Therefore, we further used the 875 annotated metabolites to predict gestational age in individual patients. After feature selection based on the Boruta algorithm (see **Methods**), 32 metabolites were selected as potential biomarkers. To ensure the robustness and reproducibility of our prediction model, we excluded those metabolites without acceptable peak shapes, then manually screened and excluded metabolites without good MS^2^ spectral matches (**Fig. S8, Fig. S9, and Methods**). Finally, 21 metabolites were included as the final biomarkers to build the prediction model in the training dataset (**Fig. 3a, Table S3**). To create an overview of the 21 metabolite biomarkers, we utilized the Classyfire algorithm (*55*) to access their chemical class information. Interestingly, most of the metabolite biomarkers were lipids and lipid-like molecules (e.g., hormones) (**Fig. 3a, Table S3**), such as 5α-preganane-3, 20-dione, pregnenolone, and progesterone, which is consistent with our previous findings from maternal plasma (*21,38–40*). Most of these metabolites had high ranks in the prediction model (ranks: 2, 4, 6, 7, 8, 9, 11, and 16; importance ratio: 44.12%; **Fig. 3a** and **Fig. S10**). Importantly, the 21 metabolite biomarkers achieved a prediction accuracy for gestational age comparable to the model that used metabolic features. Specifically, the adjusted R^2^ are 0.81 (Pearson correlation r = 0.90, *P*-value < 2.2×10^-6^) and 0.77 (Pearson correlation r = 0.87, *P*-value < 2.2×10^-6^) for internal and external validation datasets, respectively (**Fig. 3b and c**). The RMSE were 2.89 and 2.97 weeks for internal and external validation datasets, respectively (**Fig. 3b and c**). To avoid overfitting, a 1,000-time permutation test was performed, and the results suggest that our model is not overfitting (**Fig. S11**). Intriguingly, we also found that model performance improved significantly as the pregnancy progressed. As **Fig. 4c and d** show, the RMSE for both training (RMSE = 4.71 for the first trimester, 2.81 for the second trimester, 2.82 for the third trimester) and validation datasets (RMSE = 7.30 weeks for T1, 3.14 for T2, 2.81 for T3) increased from the first trimester to the third. In addition, we found that there was no significant difference in the prediction accuracy compared to the prediction model using metabolic peaks, especially in the validation dataset (metabolite model *vs.* metabolic peak model: RMSE = 2.97 *vs.* 2.66 weeks and adjusted R^2^ = 0.77 *vs.* 0.79). These results suggest that we can use urine metabolite biomarkers to predict gestational age, which has important potential clinical applications. These findings were also applied to gestational age estimation for individual participants. For the external validation dataset, 16 of the 20 participants had adjusted R^2^s larger than 0.75 (**Fig. 3d and Fig. S12**, and **Table S4**). These results indicate that our prediction model is also robust for individual prediction. Since our cohort includes women with diverse demographic and clinical characteristics (**Fig. S1a**, and **Table S1**), this also suggests that our prediction model has utility for pregnant women from diverse backgrounds. Our cohort includes a nearly two-decade age range, infant birth weight from 1,940.0 to 6,185.0 grams (IQR: 511.25), pre-pregnancy BMI from 19.49 to 57.23 (IQR: 8.39), and parity from 1 to 9 (IQR: 2) (**Table S1**). We also evaluated the impact of these personal characteristics on prediction accuracy at an individual level. The correlations between RMSE/adjusted R^2^ and continuous characteristics were calculated (**Methods**). Surprisingly, **Fig. 3e** shows that the continuous characteristics, namely, age (maternal age at birth), birth weight, pre-pregnancy BMI, and parity are not significantly correlated with prediction accuracy (Pearson correlation, all absolute correlations < 0.5 and all *P*-values > 0.05, **Methods**). Importantly, **Fig. 3e** shows that there are three outlier participants for birth weight, BMI, and parity, respectively. For participant S1760, the BMI is 57.23 (mean of all: 27.09), which is significantly different from that of most of the participants (*P*-value < 0.001). Notably, our prediction model still achieved high prediction accuracy for this participant (RMSE = 1.05, adjusted R^2^ = 0.93; **Fig. 3d**). For participant S1762, who had a parity of nine (mean of all: 2.92, *P*-value < 0.001), we also achieved good prediction accuracy (RMSE = 2.94, adjusted R^2^ = 0.90; **Fig. 3d**). Participant S1562 with birth weight 6,185.0 grams (mean of all: 3,396.5 grams, *P*-value < 0.001) was not in the external validation dataset, so only her internal validation results are shown (**Fig. 3f**). As expected, the prediction accuracy for this participant was also good (RMSE = 3.70, adjusted R^2^ = 0.95). We also tested if categorical characteristics, such as ethnicity, affected prediction accuracy. The results show that the prediction accuracy appeared to be unaffected by those characteristics. (**Fig. S13**, ANOVA test, all *P*-values > 0.05). Taken together, these findings demonstrate that the prediction model for gestational age based on metabolite biomarkers is very robust and can accommodate diversity at an individual level.

**Figure 3.**
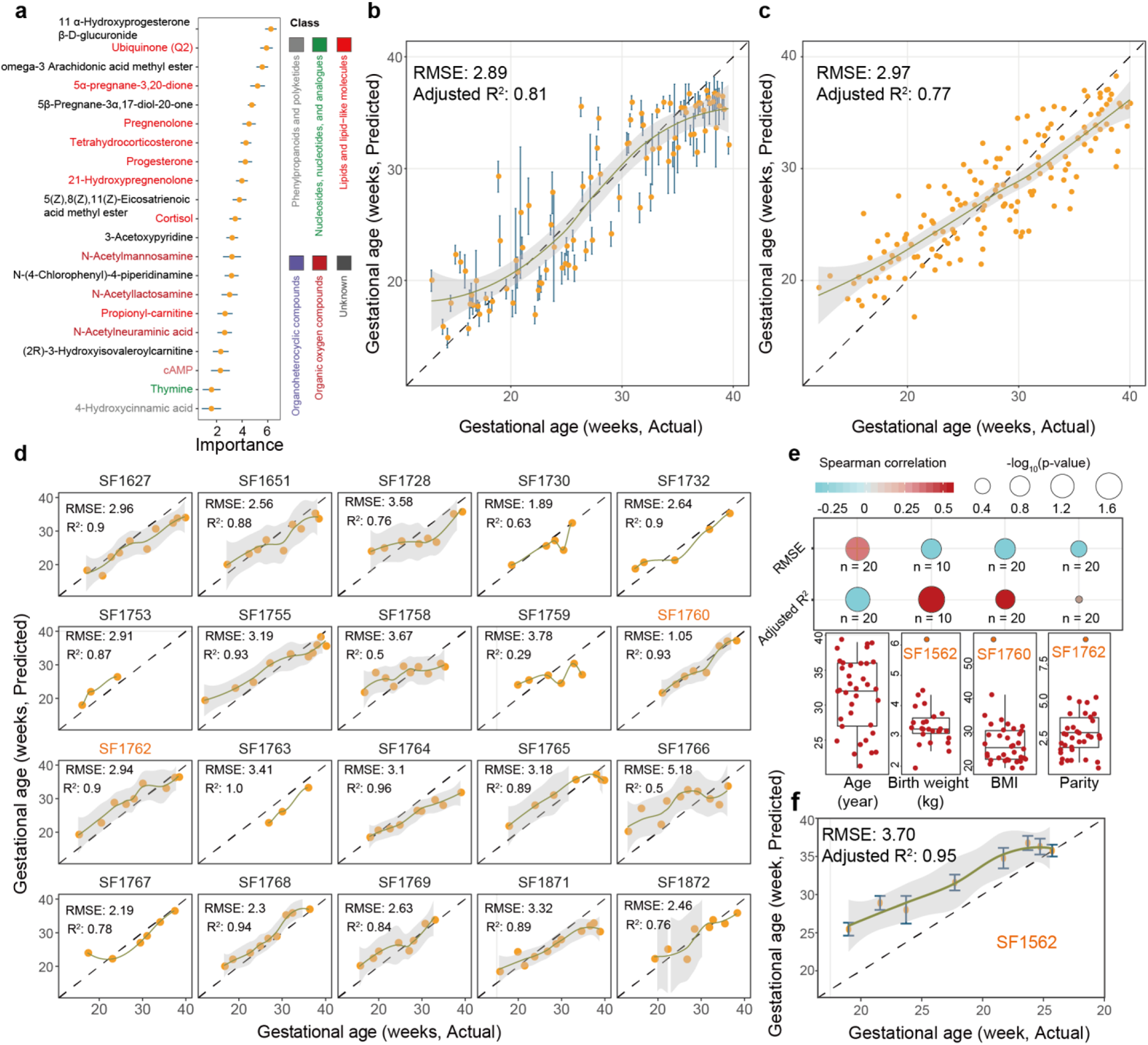
Deep urine metabolomics can predict gestational age at the individual level. (**a**) 21 metabolites were selected as biomarkers. The colors represent the chemical class of metabolites. (**b-c**) Gestational age predicted by 21 metabolites (Y-axis) is highly concordant with clinical values determined by the standard of care (first-trimester ultrasound, x-axis) in internal validation (**b**) and external validation dataset (**c**). (**d**) The prediction accuracy for each individual participant. (**e**) The continuous characteristics have no effect on gestational age prediction accuracy. (**f**) The outlier participant SF1562 in birth weight also achieves good prediction accuracy in the internal validation dataset using the bootstrap method.

**Figure 4.**
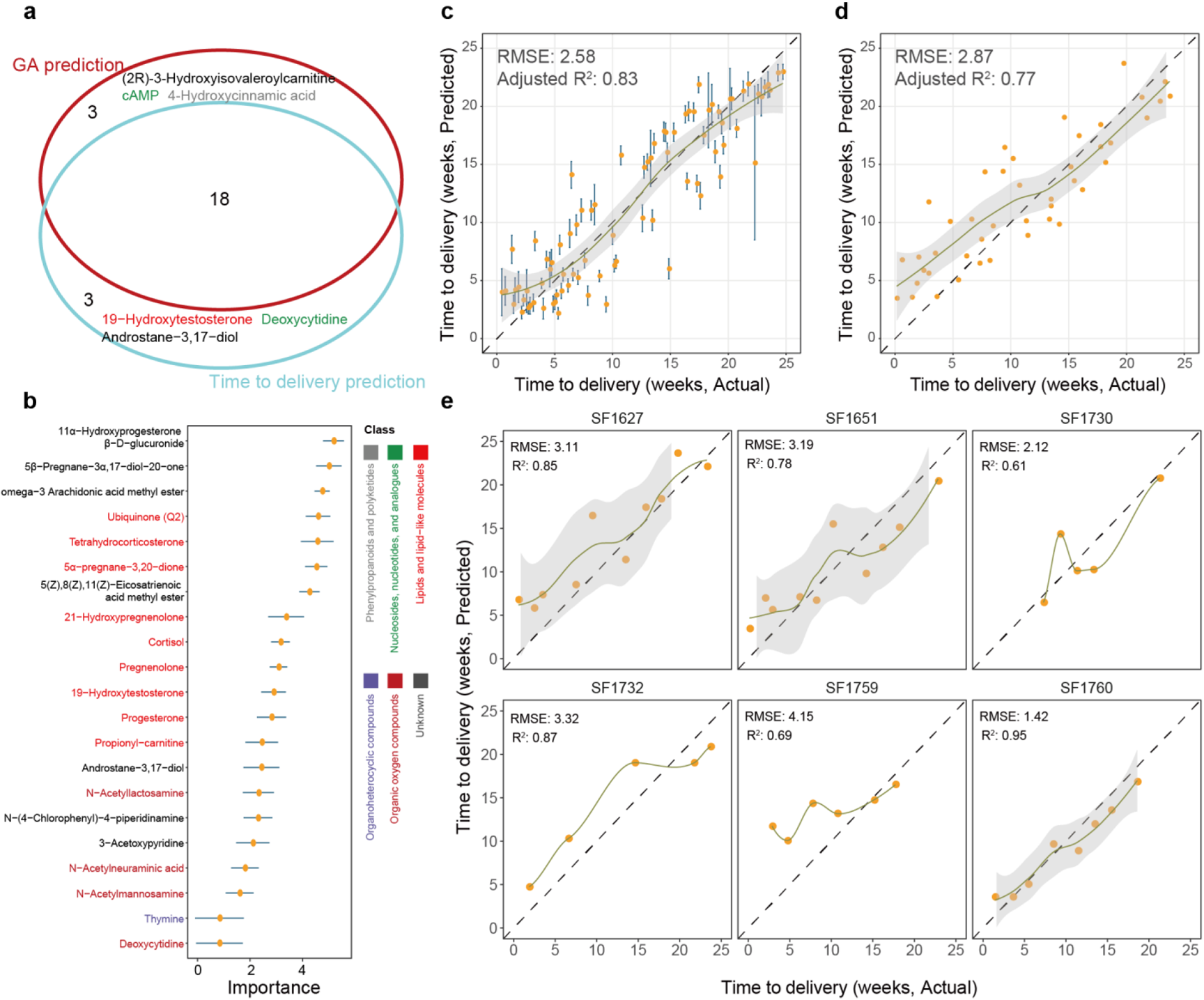
Deep urine metabolomics can predict time-to-delivery at the individual level. **(a)** The overlap between the metabolites in the prediction model for gestational age and the time-to-delivery model, respectively. **(b)** The 21 metabolite biomarkers for RF model for time to delivery model. Colors represent the chemical class of metabolites. **(c-d)** Using 21 metabolite biomarkers to build the prediction model, predicted time-to-delivery (Y-axis) is highly concordant to actual values in internal validation **(c)** and external validation dataset **(d)**. **(e)** The prediction accuracy for each individual participant.

### Prediction of Time to Delivery

We next tested whether the urine metabolome could predict time-to-delivery using annotated metabolites. “Time-to-delivery” was defined as the difference between the gestational age at sample collection and gestational age at delivery, which is a criterion independent of ultrasound-estimated gestational age. In this test, we first removed the participants who had scheduled Cesarean sections from the dataset and then used the remaining 20 participants (14 subjects for training and 6 for validation datasets, respectively) for prediction model construction and validation (see **Methods**, **Table S1**). Finally, 21 metabolites were included, 18 of which overlapped with the metabolite markers in the prediction model for gestational age (**Fig. 4a and b**, **Table S3**). The values predicted by our model agreed with actual values for both training (RMSE = 2.58 weeks; adjusted R^2^ = 0.83; Pearson correlation r = 0.94, *P*-value < 2.2×10^-6^) (**Fig. 4c**) and validation dataset (RMSE = 2.87; adjusted R^2^ = 0.77; Pearson correlation r = 0.88, *P*-value = 4.91×10^-15^) (**Fig. 4d**), showing accurate time-to-delivery prediction. The permutation test also shows that this model does not overfit the data (**Fig. S14**). Interestingly, we also found that the prediction accuracy was independent of study patient demographics, much like the case of the prediction model for gestational age (**Fig. 4e**, **Fig. S15**). These results demonstrate that the prediction model for sampling time-to-delivery based on metabolite biomarkers is also very robust and accounts for diverse characteristics on an individual level.

### Altered Metabolomic Signatures During Pregnancy

We also explored the biological function of the 24 metabolite markers that were found to differ significantly as pregnancy progressed. First, the Classyfire algorithm was utilized to determine the chemical class information of all 24 metabolite biomarkers. Most of the metabolite biomarkers (9 out of 24, 37.5%, 8 are unknown) are lipids and lipid-like molecules (hormone, **Table S3**, and **Fig. S16**), which is consistent with our finding above at the metabolic feature level. To capture the altered metabolic signatures during pregnancy, the hierarchical clustering and fuzzy c-mean clustering algorithms were utilized to group the 24 metabolite markers, which clustered into two groups with contrasting regulation patterns during pregnancy progression (**Fig. 5 (a-c)**). The first group was downregulated during pregnancy but increased to normal levels postpartum, including a panel of carnitines and signaling compounds such as cAMP (**Fig. S17a**) whereas the second group demonstrated increased abundance as the pregnancy progressed and then fell to normal levels postpartum (**Fig. S17b**). This group comprises diverse hormones and intermediates, such as 19-hydroxytestosterone, cortisol, pregnenolone, 5α-pregnane-3,20-dione, etc. These hormones were highly enriched in the glucocorticoid and mineralocorticoid biosynthesis, growth hormones, and lipid metabolism and signaling pathways. Some metabolic markers, progesterone for instance, have been applied in clinical tests for therapeutic treatment of preterm birth and pregnancy loss (*41–44*). These data suggest that other members of the steroid group with similar regulation behavior as progesterone could also serve as potential candidates for diagnostic monitoring or therapeutic targeting during pregnancy. Correlation analysis of the metabolome at different GA periods showed overarching significant alterations as the pregnancy progressed (**Fig. 5d**). The early stages of pregnancy showed a positive correlation between the metabolite intensity and GA, while the late stage showed a negative correlation. Their relative distances to postpartum levels also showed a similar pattern. This suggested significant alteration of urine metabolome is common in the later stages of pregnancy with the potential to precisely predict delivery time. The correlation between different metabolic markers demonstrated positive correlations between most markers and some potential delivery-related factors including maternal BMI and birth weight, indicating co-regulation of most metabolic pathways, except for the negative correlation between BMI and pregnenolone (**Fig. 5e**). Several studies have suggested a higher risk of preterm birth among obese women (*49–51*). Thus, the examination of pregnenolone levels could aid in GA prediction and preterm birth risk. BMI is negatively correlated with most lipid metabolite biomarkers (**Fig. S18**), although only BMI-Pregneolone demonstrated significant correlation here (FDR adjusted *P*-value < 0.05, FDR). In fact, BMI was shown to exhibit negative correlations with other lipids, except 19-Hydroxytestosterone, but none of these trends were statistically significant (FDR adjusted *P*-value > 0.05, **Fig. S18**).

**Figure 5.**
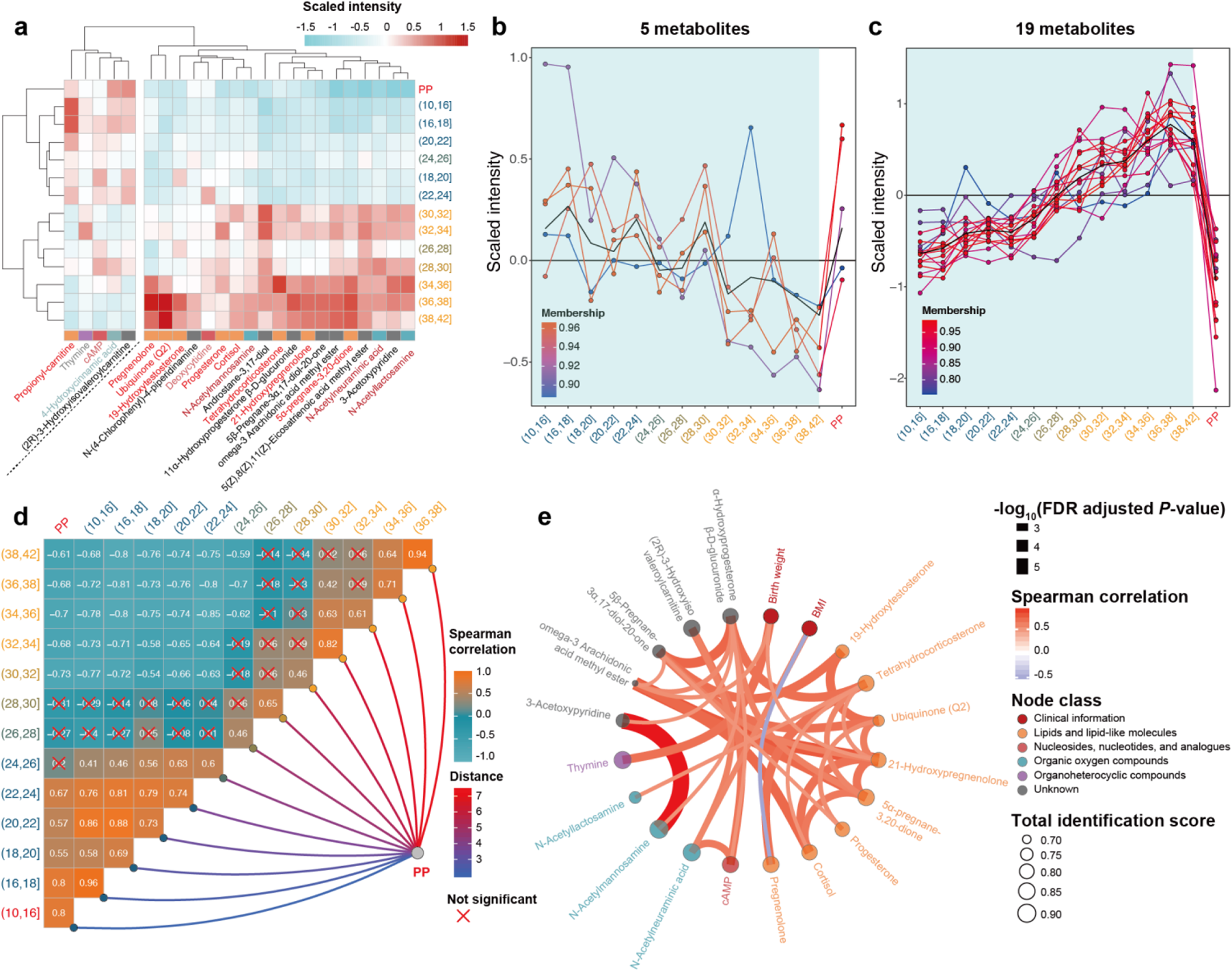
Integrative analysis of metabolic markers of gestational ages. The clustering of identified metabolites markers for gestational age prediction models. Based on different stages of gestational age (Y-axis, showing gestational weeks), markers were clustered into two main groups, one was upregulated in early stages and downregulated in late stages, while the other group showed a contrast pattern, with an upregulation in late stages. **(b-c)** The fuzzy-c mean clustering of metabolite biomarkers based on gestational weeks. The identified metabolite markers could be clustered into two groups, one with a consistent downregulation as pregnancy progresses followed by a return to normal levels postpartum. (**b**) the other group showed an upregulation during pregnancy then was downregulated postpartum (**c**). **(d)** The correlation of metabolome alternations at different gestational weeks and their relative distance to postpartum. The early stage of pregnancy showed a positive correlation with postpartum, while the late stage showed a negative correlation with postpartum, confirming significant alternations of the urine metabolome in the later stages of pregnancy **(e)** The correlation between different metabolic markers. Sizes of nodes showed the total identification score. Colors of nodes demonstrated different classes of metabolites. Colors and widths of the cords showed correlation and adjusted p-values. We also included BMI and birth weight into the correlation analysis and found a negative correlation between BMI and pregnenolone.

## DISCUSSION

A reliable estimate of gestational age is critical for the provision of preventive prenatal health care and appropriate interventions as the medical needs of the mother and fetus change throughout pregnancy ^21-52^. Although substantial work has been done to elucidate the dynamic metabolic pathways of pregnancy progression using collected blood samples (*18,19,21,53*), the dynamic pregnancy urine metabolome has been only sparsely characterized. For this study, we applied an unbiased comprehensive metabolic profiling approach to analyze urine samples from pregnant women who were participants in the SMART-D cohort to better understand prenatal and post-natal metabolic dynamics in maternal urine. We developed models to estimate gestational age at the time of sampling and predictions for time-to-delivery from sampling. Metabolic models for gestational age at time-to-delivery were validated at the cohort and individual levels and found to be highly predictive (training database: adjusted R^2^ 0.81, RMSE 2.89; test database: R^2^ 0.77, RMSE 2.97) (Fig. 3c, 3d). Importantly, prediction was found to improve with increased sampling frequency. Our gestational age model tended to overestimate gestational age during early pregnancy and underestimate gestational age in later pregnancy. These discrepancies may be due to underlying heterogeneous biological processes happening throughout pregnancy and will need further investigation in a larger population. Overall, our findings suggest that the pregnancy urine metabolome can be successfully leveraged to estimate gestational age at sampling and to predict time-to-delivery.

Pregnenolone, progesterone and corticoid were all upregulated in the glucocorticoid pathways during pregnancy and related metabolites, used in the time-to-delivery prediction model, were enriched for glucocorticoid and CMP-N-acetylneuraminate biosynthesis pathways. These hormones have been reported to play key roles in pregnancy regulation (*42–44*). For instance, progesterone has been approved for the treatment of amenorrhea, metrorrhagia, and infertility (*45–48*). In our previous study of maternal plasma collected in an independent cohort, we also identified tetrahydrodeoxycorticosterone (THDOC), estriol glucuronide, and progesterone as potential markers for GA estimation (*21*). These markers were further validated by our findings in maternal urine. Other identified derivatives in the same steroid hormone group of estrogens and progesterone derivatives, as well as uncharacterized steroid-like compounds discovered in this study, may also play roles in pregnancy although their functions remain unclear. Furthermore, N-acetylmannosamine and N-acetylneuroaminate were both significantly upregulated in the CMP-N-acetylneuraminate biosynthesis pathway, although the impact of these signaling molecules on pregnancy-related processes remains to be explored.

As a proof-of-principle, our results show that urine metabolomic profiles can be used to track gestation throughout pregnancy. By applying a random forest model, we successfully predicted gestational age based on a panel of urine metabolites (Fig 3c, 3d), including diverse glucocorticoids, lipids, gluconoids, and amino acid derivatives, indicating comprehensive regulation of glucocorticoid biosynthesis and CMP-N-acetylneuraminate biosynthesis by pregnancy.

Collectively, the characterized alterations in the maternal urine metabolome demonstrated a strong correlation between GA and pregnancy progression. The ability to determine GA accurately and conveniently and to predict time-to-delivery will aid in monitoring fetal development and targeting interventions to improve maternal and infant health outcomes. This study also revealed substantial differences between maternal urine metabolites in healthy pregnancies and those ending in preterm birth, which suggests variation of metabolic regulation mechanisms among different pregnancy outcomes. Close monitoring of maternal metabolism alterations and their correlation with GA has the potential to provide more insight into the programmed regulation of fetal development and the development of pregnancy disorders. The non-invasive nature and accessibility of the urine metabolome enables improved determination of GA without limitation across various clinical settings, including in middle-to-low-income countries where women may have limited access to early prenatal care.

## MATERIALS AND METHODS

### Participant enrolment and urine sample collection

Sample collection: 346 urine samples were collected at multiple time points during the pregnancy process (11.8–40.7 weeks) and postpartum period for 36 healthy women. The SMART-D cohort represents an ethnically diverse group of participants with a wide range age and BMI distribution (**Table S1**, **Fig. S1b**). The samples were collected longitudinally and delivered to analysis into two batches (**Table S5**).

### Gestational age (GA) estimation

The American Congress of Obstetricians and Gynecologists (ACOG) guidelines were used to define accurate gestational age dating for all the participants in the SMART-D cohort.

### Chemical material and internal standard preparation

MS-grade water, methanol and acetonitrile were purchased from Fisher Scientific (Morris Plains, NJ, USA). MS-grade acetic acid was purchased from Sigma Aldrich (St. Louis, MO, USA). Analytical grade internal standards were purchased from Sigma Aldrich (St. Louis, MO, USA). The internal standard mixture of acetyl-d3-carnitine, phenylalanine-3,3-d2, tiapride, trazodone, reserpine, phytosphingosine, and chlorpromazine was 1: 50 diluted with 3:1 acetonitrile and water for HILIC, and water for RPLC.

### Urine sample preparation

Urine samples were thawed and centrifuged at 17,000 rcf for 10 min. 250 μL supernatant was diluted with 750 μL internal standard mixture, vortex for 10 seconds and centrifuged at 17,000 rpm for 10 min at 4 °C. The supernatant was taken for subsequent LC-MS analysis. A quality control (QC) sample was generated by pooling up all the samples and injected between every 10 sample injections to monitor the consistency of the retention time and the signal intensity.

### LC-MS data acquisition

Hypersil GOLD HPLC column and guard column was purchased from Thermo Scientific (San Jose, CA, USA). Mobile phases for RPLC consisted of 0.06% acetic acid in water (phase A) and MeOH containing 0.06% acetic acid (phase B). Metabolites were at a flow rate of 0.25 mL/min, leading to a backpressure of 120-160 bar at 99% phase A. A linear 1 - 80% phase B gradient was applied over 9 - 10 min. The heating temperature of the column was set to 60 °C and the sample injection volume was 5 μL.

MS acquisition was performed on an Q Exactive HF Hybrid Quadrupole-Orbitrap mass spectrometer (Thermo Scientific, San Jose, CA, USA) cooperating in both the positive and negative ESI mode (acquisition from *m/z* 500 to 2,000) with a resolution set at 30,000 (at *m/z* 400). The MS^2^ spectrum of the QC sample was acquired under different fragmentation energy (25 eV and 50 eV) of the top 10 parent ions.

### Data processing and cleaning

#### Raw data processing

First, all the MS raw data (.raw format) were converted to .mzXML (MS^1^ raw data) and .mgf format data (QC MS^2^ data) using ProteoWizard software (http://proteowizard.sourceforge.net/). The detailed parameter settings for the data conversion are listed in **Table S6**. Second, all the .mzXML format data was grouped into 3 folders (named as “Blank”, “QC” and ”Subject”) and then subjected for the peak detection and alignment. Third, the peak detection and alignment were performed using our in-house pipeline. Briefly, the peak detection and alignment were performed using the centWave algorithm (R package **xcms**, version 3.8.1). The key parameters were set as follows: method = “centWave”; ppm = 15; snthr = 10; peakwidth = c(5, 30); snthresh = 10, prefilter = c(3, 500); minifrac = 0.5; mzdiff = 0.01; binSize = 0.025 and bw = 5. Finally, the generated MS^1^ peak table includes the mass-to-charge ratio (*m/z*), retention time (RT), and peak abundances for all the samples, and other information. This MS^1^ peak table is used for the next data cleaning.

#### Data cleaning

The data cleanings of the peak table were also performed using our in-house pipeline. First, the peaks detected in less than 20 % QC samples were removed from the peak table as noisy. Second, the samples with more than 50% missing values were removed as outlier samples. Third, the remaining missing values (NA) were imputed using the k-nearest neighbors (KNN) algorithm (R package **impute**). Then, the peak intensity was divided by the mean peak intensity for data normalization to remove the unwanted analytical variations occurring intra-batches. Finally, the ratio of mean values of each peak in two batches was utilized as the correct factor to do data integration. The in-house pipeline for data processing and data cleaning, script, and parameter setting can be found in GitHub (https://github.com/jaspershen/metflow2).

### General statistical analysis and data visualization

The majority of the statistical analysis and data visualization is performed using Rstudio (version 1.2.5019) and R language (version 3.6.0) in a Windows 10 X 64 OS. Most of the R packages and their dependencies used in this study are maintained in CRAN (https://cran.r-project.org/) or Bioconductor (https://bioconductor.org/). The directory used R packages are **plyr** (version 1.8.5), **stringr** (version 1.4.0), **dplyr** (version 0.8.3), **purrr** (version 0.3.3), **readr** (version 1.3.1), **readxl** (version 1.3.1), **tidyr** (1.0.0), **tibble** (version 2.1.3), **ggplot2** (version 3.2.1), **ggsci** (version 2.9), **patchwork** (version 1.0.0), and **igraph** (1.2.4.2). All information about the used R packages in data analysis are provided in **Table S7**. The main script for analysis and data visualization in this study is provided in GitHub (https://github.com/jaspershen/smartD_project).

In general, before all the statistical analysis, the data are first log10 transformed and then auto scaled as flow:

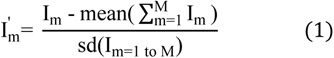

The categorical data are described as the frequency counts and percentages, and the values of all continuous variables are presented as the mean plus or minus the standard deviation (SD) or standard error of the mean (SEM). Most metabolic peaks showed right-skewed distribution (**Fig. S8**); thus, the nonparametric methods (Wilcoxon rank-sum test, spearman correlation) are utilized for non-parametric statistical tests. All the *P*-values are adjusted utilizing False Discovery Rate (FDR, R base function ***p.adjust***). PCA analysis is performed utilizing the R base function **prcomp**. The R package **ggplot2** (version 3.2.21) was used to perform most of the data visualization in this study.

### SAM test and linear regression model to detect overall altered metabolic peaks during pregnancy

To find the metabolic peaks which significantly changed according to GA during pregnancy, significance analysis of microarray (SAM) and linear regression model between GA and metabolic peaks were utilized. SAM assigns a score to each metabolic peak on the basis of change in peak expression relative to the standard deviation of repeated measurements. For metabolic peaks with scores greater than an adjustable threshold, SAM uses permutations of the repeated measurements to estimate the percentage of metabolic peaks identified by chance, the false discovery rate (FDR). We performed SAM utilizing the SAM function in R package **samr**, and resp.type was set as “Quantitative” and FDR was set as 0.05 with 1,000 permutation tests. For the linear regression model, the R base function lm was utilized. To adjust the potential confounders, the participants’ baseline, namely acquisition batch, BMI, mother age, parity, ethnicity were also imputed into the linear regression model, and the metabolic peaks with FDR adjusted *P*-value < 0.05 were selected. Finally, only the 3,020 metabolic peaks (2,436 increased metabolic peaks and 584 decreased metabolic peaks) that are significant in both two methods were used for subsequent K-means consensus-clustering analysis.

### K means consensus-clustering

Unsupervised K means consensus-clustering of the 302 urine metabolome samples was performed with the R package **CancerSubtypes and ConsensusClusterPlus** using the 3,020 metabolic peaks that were discovered by SAM and linear regression model. The data was log10-transformed. Samples clusters were detected based on K-means clustering, Euclidean distance and 1,000 resampling repetitions in ExecuteCC function in the range of 2 to 6 clusters. The generated empirical cumulative distribution function (CDF) plot initially showed optional separation 2 and 3 clusters for all urine samples. And from the consensus matrix heatmaps we can also 2, 3 and 4 clusters seem to have good clustering. To further decide how many clusters (k) should be generated, the silhouette information from clustering was extracted using silhouette_SimilarityMatrix function. We compared k = 2, 3, 4, and found that when k = 3 we get the high stability for clustering (**Fig. S4**). So finally, all the urine samples were assigned to 3 clusters (**Fig. 1d**).

### PIUMet analysis

PIUMet is a network-based tool (http://fraenkel.mit.edu/PIUMet/) which infers putative metabolites corresponding to features and molecular mechanisms underlying their dysregulation, which means that they can transfer metabolic peak information to network information. For each GA range, the altered metabolic peaks were outputted as txt format files with three columns: m/z, polarity, and −log_10_ (FDR adjusted *P*-value) and then uploaded into the PIUMet website. The parameters are set as below: number of trees: 10, edge reliability: 2, negative prize degree: 0.0005, and number of repeats: 1. Then all the results from PIUMet are processed by an in-house pipeline which is provided in our GitHub (https://github.com/jaspershen/smartD_project). Briefly, all the annotation results from PIUMet for each GA range were combined, and if one metabolite matches more than one metabolic peak, only the metabolic peaks that appeared more than two GA ranges were kept, and other metabolic peaks were removed. Then for each metabolite, the matched metabolic peaks are used to extract quantitative information and the mean values were used as quantitative values for this metabolite.

### Correlation network and community analysis

Due to most metabolic peaks (metabolites) are not normally distributed across all urine samples, so the Spearman correlation was used to build the correlation in our analysis. For the annotation result from PIUMet and dataset combined by metabolite and clinical variables, the correlations between each pair variable were calculated, and then only the absolute correlation > 0.5 and FDR adjusted *P*-value < 0.05 were kept to construct correlation networks.

We performed community analysis using the method based on edge betweenness developed by Girvan and Newman which is embedded in R package **igraph**. In a network, the edge betweenness score of an edge measures the number of shortest paths through it. So, the idea of edge betweenness based community structure detection is that it is likely that edges connecting separate modules have high edge betweenness as all the shortest paths from one module to another must traverse through them. Briefly, this is an iterative process, in each iteration, the edges with the highest edge betweenness score were removed, and the process was repeated until only individual nodes remain. Finally, we will get a hierarchical map, a rooted tree, called a dendrogram of the graph. The leaves of the tree are the individual nodes, and the root of the tree represents the whole graph (network). Then an unbiased method, modularity of the detected community structure was used to analyze the correlation network at a cut level. The modularity of community structure corresponds to an arrangement of edges that is statistically improbable when compared to an equivalent network with edges placed at random. At every iteration of the community analysis, we computed the modularity and analyzed the communities at the iteration which maximized this quantity. A visualization of community modularity vs. iteration is shown in Figure **S7a**. To make sure that our findings are robust and reliable, only the communities (or clusters) with at least 3 modes were kept for subsequent analysis. All the networks were visualized using R package **ggraph** (version 2.0.0).

### Kyoto Encyclopedia of Genes and Genomes (KEGG) pathway enrichment analysis

The human KEGG pathway database is downloaded from KEGG (https://www.genome.jp/kegg/) utilized R package KEGGREST. The original KEGG database has 275 metabolic pathways, and then we separated it into metabolic pathways and disease pathways based on the “Class” information for each pathway, the pathways with “Human Disease’’ class were assigned into the disease pathway database, which contains 74 pathways and remained 201 pathways were assigned into metabolic pathway database. The pathway enrichment analysis is used in the Hypergeometric distribution test. *P*-values are adjusted by the FDR method and the cutoff was set as 0.05. The code is an in-house pipeline and can be found on GitHub (https://jaspershen.github.io/smartD_project/).

### Metabolite annotation

To achieve accurate metabolite annotation for this study, three criteria, (1) accurate mass (*m/z*), (2) retention time (RT) and (3) MS^2^ spectral similarities are used for metabolite annotation. The public MS^2^ spectral databases have no retention times for standards, so only the accurate mass and MS^2^ spectral similarity are used. For each matching, we calculate the match score to represent the match similarity. Each score gives the standardized range from 0 to 1, meaning from no match (0) to a perfect match (1), respectively. Metabolite annotation was first performed using our in-house pipeline based on our in-house and downloadable public MS^2^ spectra databases. Then the remaining unannotated metabolic peaks were matched with the online databases NIST (https://chemdata.nist.gov/) and METLIN.

For the metabolite annotated using the in-house database, we have accurate mass (*m/z*), retention time (RT) and MS^2^ spectra, so the annotations are level 1 according to MSI. For metabolites annotated using the public databases, only accurate mass and MS^2^ spectra are used for matching, so the annotation is level 2 according to MSI.

#### Data organization

All the MS^2^ spectra (.mgf format) from QC samples were matched with MS^1^ peaks in peak table according to accurate mass (*m/z*, tolerance is set as ±25 ppm) and RT (tolerances is set as ±10 seconds) using the code provided by MetDNA [Metabolic reaction network-based recursive metabolite annotation for untargeted metabolomics]^56^. If one MS^1^ peak matches multiple MS^2^ spectra, only the most abundant MS*^2^* spectrum is kept. Finally, the generated MS^1^/MS^2^ pairs were used to match with our in-house and public MS^2^ spectral databases (HMDB [http://www.hmdb.ca/], MoNA [https://mona.fiehnlab.ucdavis.edu/], and MassBank [https://massbank.eu/MassBank/]).

#### Accurate mass and RT match score

The match tolerance for the MS^1^ *m/z* value is set as ± 25 ppm and RT match tolerance is set as ±10 seconds. Only the metabolites that meet those tolerances are kept. The match scores refer to MS-DIAL and are calculated as follows:

Accurate mass

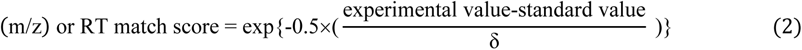

Where the experiment value is the experimental *m/z* or RT from MS^1^ peak table, and the standard value is the standard *m/z* or RT from MS^2^ spectral databases. These equations are based on the assumption that for accurate mass and retention time match scores, the differences between experimental and standard values follow the Gaussian distribution (normal distribution). The standard deviation δ is the accurate mass (*m/z*) or RT match tolerance.

#### MS^2^ spectral match score

The MS^2^ spectral match score is a combined value of three scores, namely forward dot-product (DP_f_), reverse dot-product (DP_r_), and the matched fragments ratio (MFR). Both the DP scores and MFR ranges are from 0 to 1, meaning from no match (0) to a perfect match (1). The intensities of the fragment ions in the MS^2^ spectra are rescaled so that the highest fragment ion is set to 1.

The forward and reverse dot-product are calculated as follow:

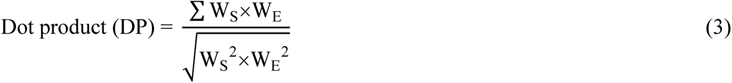

Where the weighted intensity vector, W = [relative intensity of fragment ion]^n^[*m/z* value]^m^, n = 1, m = 0; S = standard and E = experiment. DP from both forward and reverse matches are generated using this equation.

The matched fragment ratio (MRF) is utilized to assess how many fragments are matched in all fragments in both experiment and standard MS^2^ spectra and is calculated as follow:

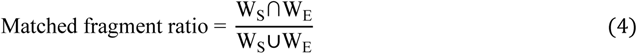

Where the weighted intensity vectors are the same as the equation (2). Ws∩W_E_ mean the number of matched fragments between standard and experiment MS^2^ spectra, and Ws∪W_E_ mean the number of all the fragments in standard and experiment MS^2^ spectra.

Finally, the MS^2^ spectral match score is combined the forward DP (DP*_f_*), reverse DP (DP*_r_*) and matched fragment ratio (MFR), and the weight for forward DP (W*_f_*), reverse DP (W*_r_*), and matched fragments ratio (W*_m_*) are set as 0.3, 0.6 and 0.1, respectively.

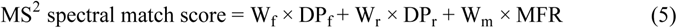

#### Total match score

Three match scores, namely accurate mass, retention time, and MS2 spectra match scores, are used to calculate the total match score as follow:

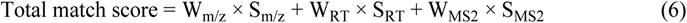

Where W_m/z_, W_RT_, and W_MS2_ are weighted for accurate mass (S_m/z_), RT (S_RT_), and MS2 (S_MS2_) spectral match scores, and set as 0.25, 0.25, and 0.5, respectively. For public MS^2^ spectral databases without RT information, the above three weights are set as 0.375, 0, and 0.625, respectively.

If one metabolic peak matches multiple metabolites, the annotated metabolites are sorted according to the total match score. And for all the potential metabolite markers in our study, we finally manually checked to confirm the accuracy of metabolite annotation.

### Metabolite biomarker confirmation

To make sure that the metabolite biomarkers we selected with the right annotations and have good reproducibility, we manually checked all the metabolite biomarkers. Two criteria, (1) peak shape, and (2) MS^2^ spectra match are included. Only the metabolite biomarkers with good peak shapes remain for prediction model construction. Metabolite biomarkers that have bad peak shapes may have bad reproducibility, so they are discarded. The metabolite biomarkers that have bad MS^2^ spectra match with standards are also removed to avoid the wrong annotation.

### The Random Forest prediction model

#### Feature selection

The Boruta algorithm (R package **Boruta**, version 6.0.0) is utilized to select potential biomarkers. Briefly, it duplicates the dataset and shuffles the values in each column. These values are called shadow features. Then, it trains a Random Forest classifier on the dataset, and checks for each of the real features if they have higher importance. If it does, the algorithm will record the feature as important. This process is repeated 100 iterations. In essence, the algorithm is trying to validate the importance of the feature by comparing it with randomly shuffled copies, which increases the robustness. This is performed by comparing the number of times a feature did better with the shadow features using a binomial distribution. Finally, the confirmed features are selected as potential biomarkers for Random Forest model construction.

#### Parameter optimization

All the parameters are used as default settings except ntree (*i.e.*, number of trees to grow) and mtry (*i.e.*, number of variables randomly sampled as candidates at each split) in the Random Forest model (R package **randomForest**, version 4.6-14). Those two parameters are optimized on the training dataset. The two parameters are combined together to form a set of parameter combinations. The performance of each parameter combination is evaluated using the mean squared error (MSE). The parameter combination with the smallest MSE is used to build the final prediction model.

#### Gestation age (GA) prediction model

All the samples acquired in batch 1 (16 subjects and 125 samples) are used as the training dataset. All the samples acquired in batch 2 (20 subjects and 156 samples) are used as the validation samples. First, the training dataset is utilized to get the potential biomarkers using the feature selection method described above. Then a Random Forest prediction model is built based on the training dataset. Then based on this prediction model, we also construct a linear regression model between predicted GA and actual GA (**Fig. S19**). Then the predicted GA from Random Forest is corrected by this linear regression model. So, the GA prediction model contains two models, namely Random Forest and linear regression model. Then the external validation model is utilized to demonstrate its prediction accuracy. The predicted GA and actual GA for the validation dataset are plotted to observe the prediction accuracy. Then the RMSE (root mean squared error) and adjusted R^2^ are used to quantify the prediction accuracy.

For internal validation, the bootstrap sampling method is utilized. Briefly, we randomly sampled the same number of samples from the training dataset with replacement (about 63% of the unique samples on average) and then used it as an internal training dataset to build the Random Forest prediction model using the same selected features and optimized parameters. The remaining about 37% of the samples on average were used as internal validation data. Those steps repeat 1,000 times. Finally, for each sample, we got more than one predicted GA value. The mean value of multiple predicted GA values is used as the final average predicted GA and used to calculate MSE and adjusted R^2^.

#### Sampling time to the delivery prediction model

Sampling time to delivery (week) is defined as the time difference between the delivery date and sample collection date. So, for each sample, time to delivery is calculated. Then we used those as responses to build a prediction model. All the steps are the same as the GA prediction model.

### Permutation test

The permutation test was utilized to calculate p-values to judge if the random forest prediction models that we constructed are not overfitting. First, all the responses (GA or time to delivery in this study) are randomly shuffled for both training and validation datasets, respectively. Then the potential biomarkers are selected and the parameters of random forest are optimized in the training dataset using the method described above. Thirdly, the random forest prediction model is built using the selected features and optimized parameters in the training dataset. Finally, we use this random forest prediction model to get the predicted responses for the validation dataset. Then we get the null RMSE and adjusted R^2^. We repeat this process 1,000 times, so we get 1,000 null RMSE and 1,000 null adjusted R^2^ vectors. Using maximum likelihood estimation, these null RMSE values and adjusted R^2^ values are modeled as Gamma distribution, and then the cumulative distribution function (CDF) is calculated. Finally, the *P*-values for the real RMSE and adjusted R^2^ are calculated from the null distributions, respectively.

### Fuzzy c-means clustering

The fuzzy c-means clustering algorithm (R packages **e0171** and **Mfuzz**) is utilized to cluster the metabolite biomarkers into different classes and explore the metabolite changes according to the gestation age (weeks). Because the participants’ samples were collected at different time points, so all the samples are grouped to different time ranges. The time ranges are from 11 weeks to 41 weeks and step is two, and the postpartum samples are grouped to the “PP” group. For the samples in the same time range group, each metabolite’s intensity is calculated by the mean value of all the samples in this group. So finally, we get a new data frame with 16 new observations. First, we optimized the parameter “m” (the degree of fuzzification) based on a method using the **Mfuzz** package. The optimal cluster number is determined based on the within-cluster sum of squared error. Then we used all default parameters to build the fuzzy c-means clustering. For each cluster, only the features with a membership score > 0.5 were considered, we chose this high stringency so that we can explore the dynamics of the core members of each cluster. In fuzzy c-means clustering, the membership score is the probability of a feature belonging to any cluster, each feature is assigned a cluster based on its top membership score (as opposed to k-means clustering, where the membership score is binary). The color of each feature is directly based on the membership score (from blue to red, membership score from low to high). The output results were not smoothed.

---

**Table S1.** The detailed information for all the participants in SMART-D study.

**Table S2.** 28 potential metabolic peak markers to predict gestational age.

**Table S3.** The detailed information of 24 metabolite biomarkers to predict gestational age and sampling time to delivery.

**Table S4.** Prediction result for each participant in prediction gestational age model.

**Table S5.** The detailed information for each urine sample in SMART-D study.

**Table S6.**
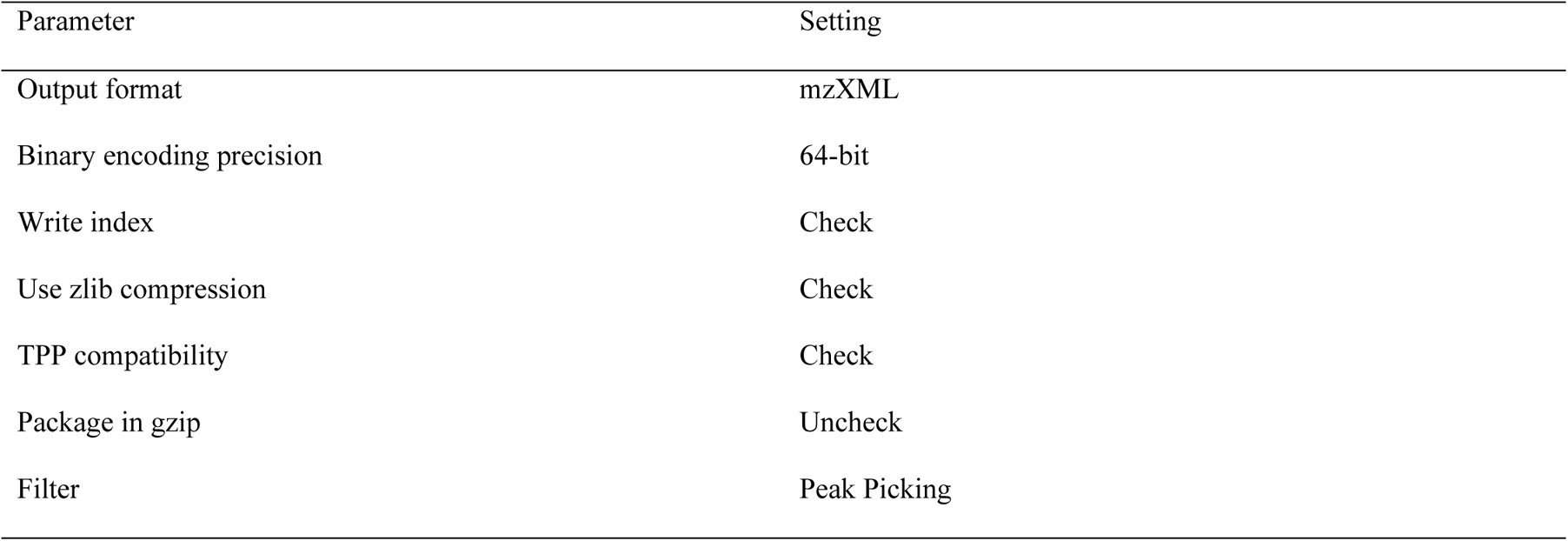
Parameters for MS raw data conversion using ProteoWizard.

**Table S7.**
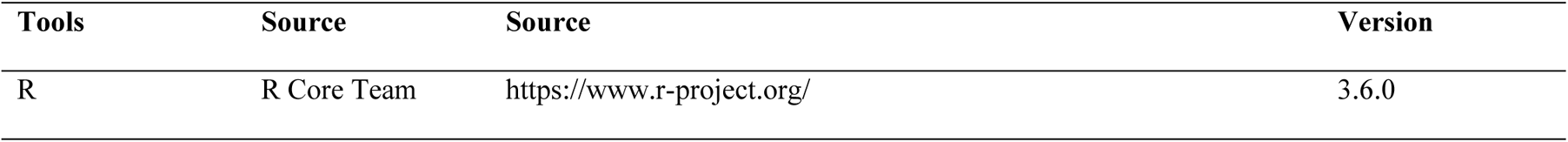
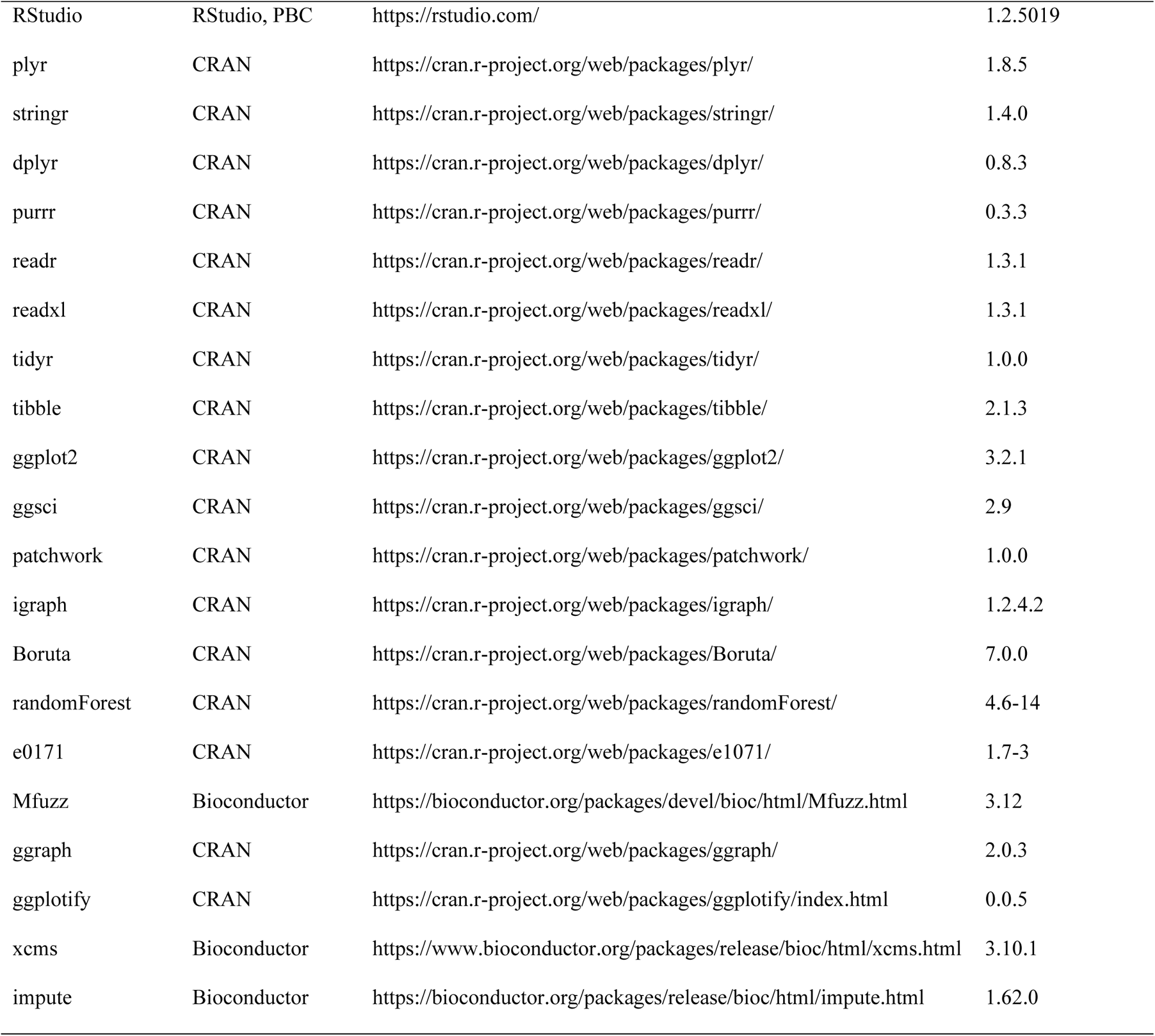
All software and R packages used in data analysis.

**Figure S1.**
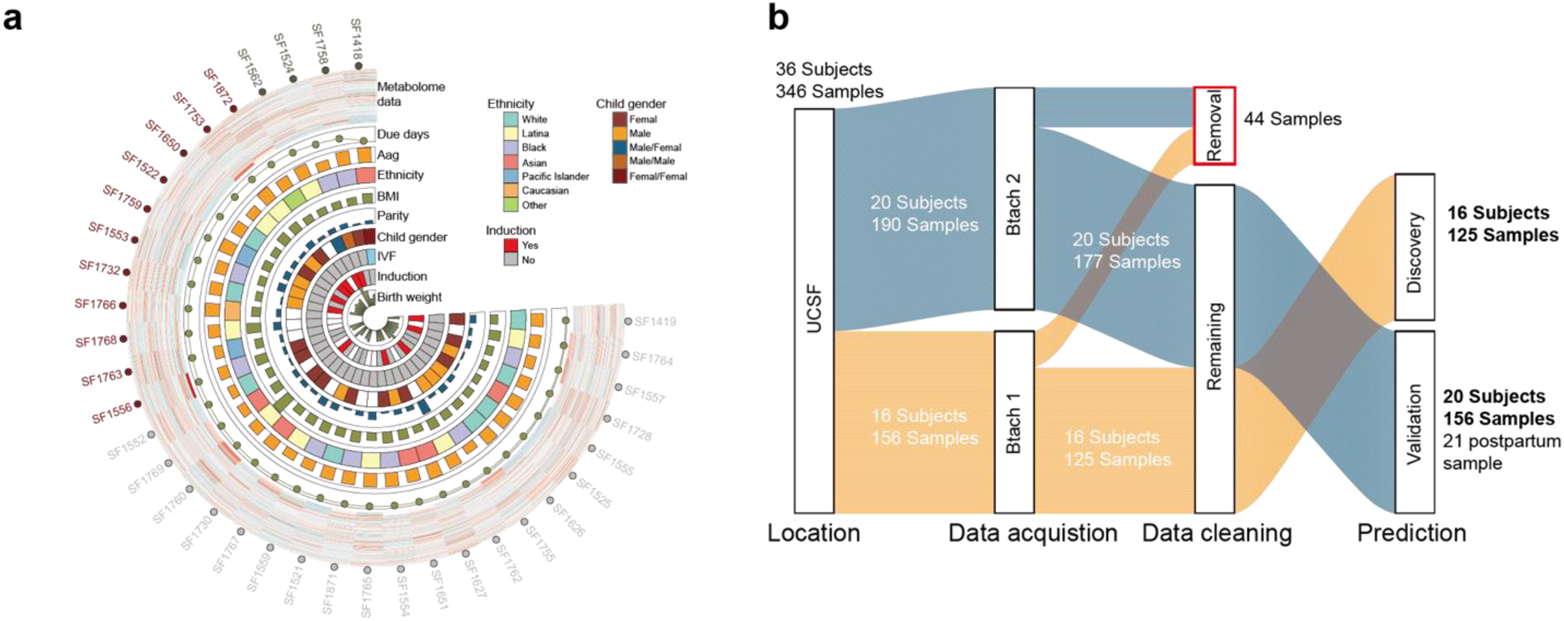
Study design of SMART-D study. (**a**) The metabolome and clinical information of 36 participants in SMART-D study. (**b**) Sample collection, data acquisition, data processing and analysis design for our study. UCSF (University of California, San Francisco), ZSFGH (Zuckerberg San Francisco General Hospital).

**Figure S2.**
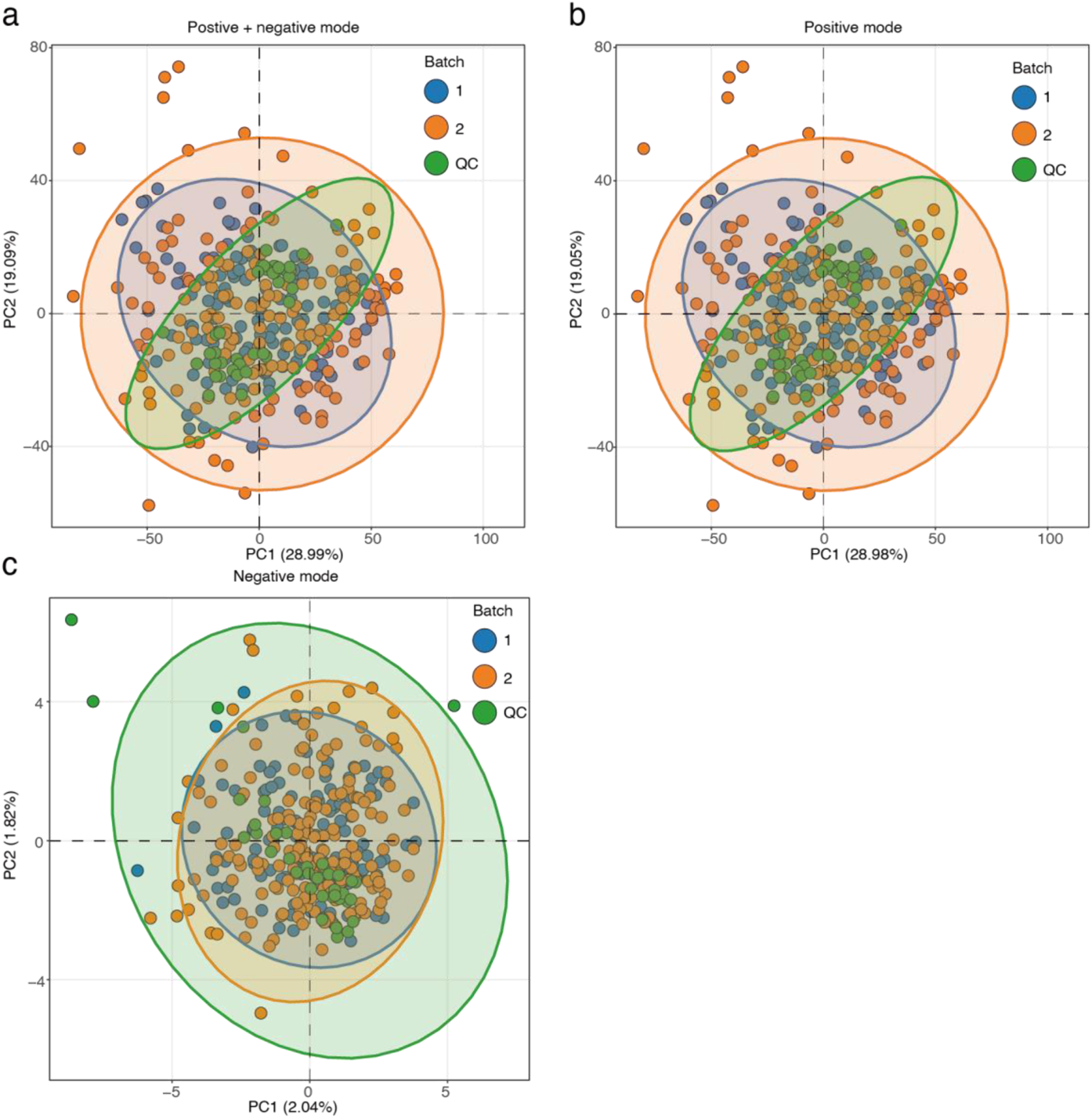
Data quality of urine metabolomics data. (**a**) Positive and negative mode data. (**b**) Only positive mode data. (**c**) Only negative data.

**Figure S3.**
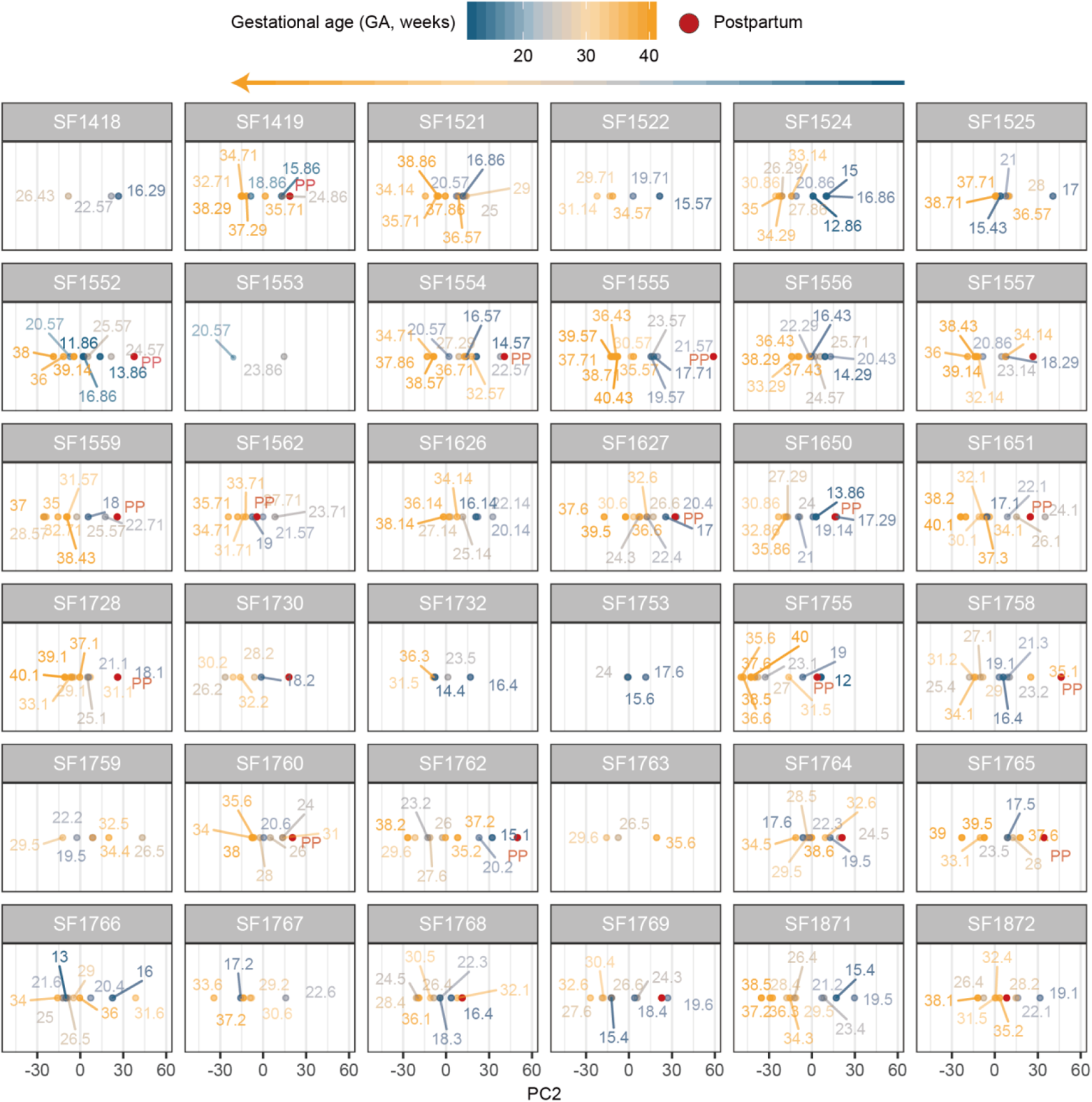
PCA score plot for each participant in SMART-D study.

**Figure S4.**
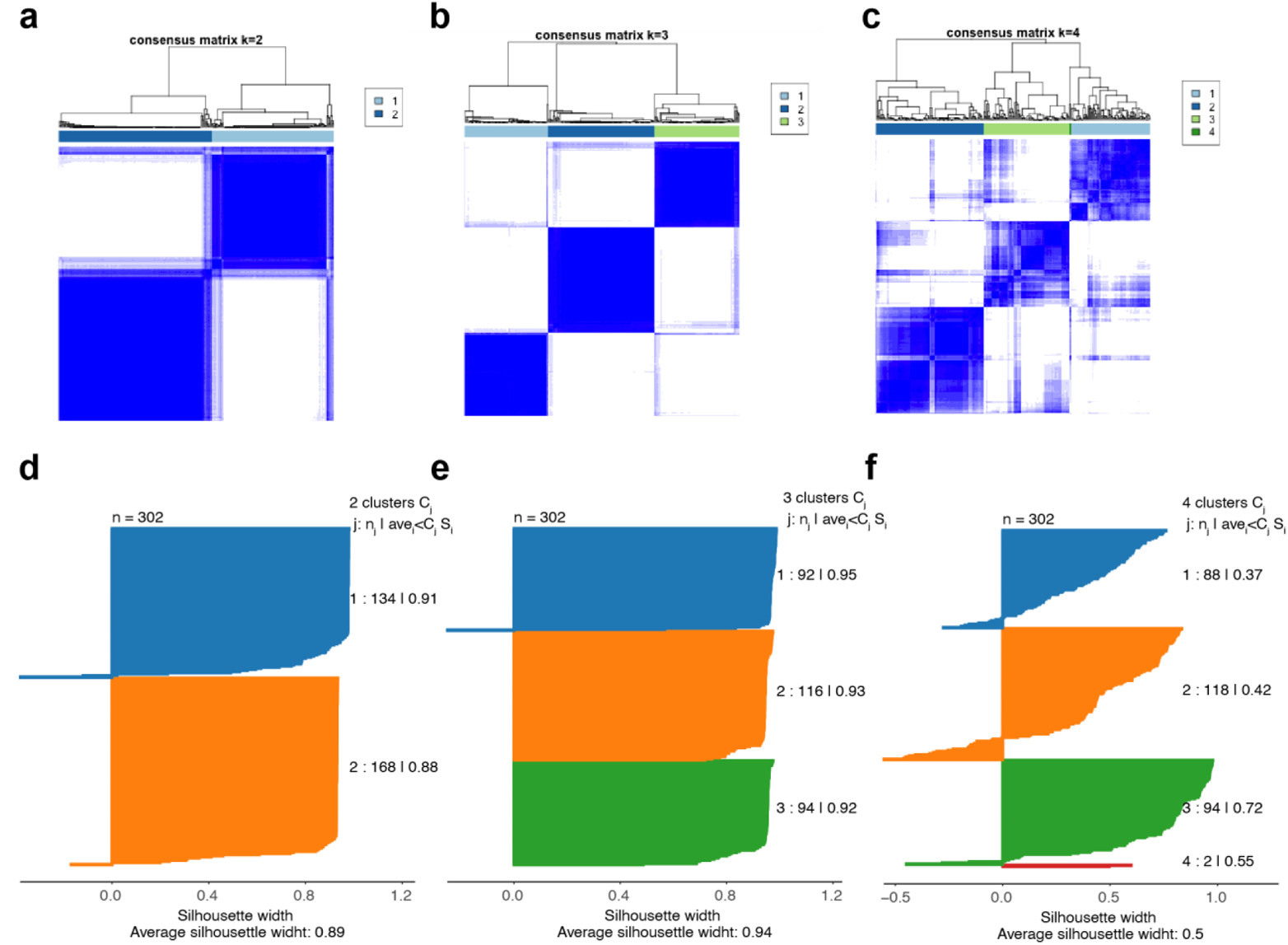
K means consensus-clustering for urine metabolome in 302 samples. The heatmaps of consensus matrix for k = 2 (**a**), k = 3 (**b**) and k = 4 (**c**) clusters based on 1,000 resampled datasets. The silhouette plots for k = 2 (**d**), k = 3 (**e**) and k = 4 (**f**) clusters.

**Figure S5.**
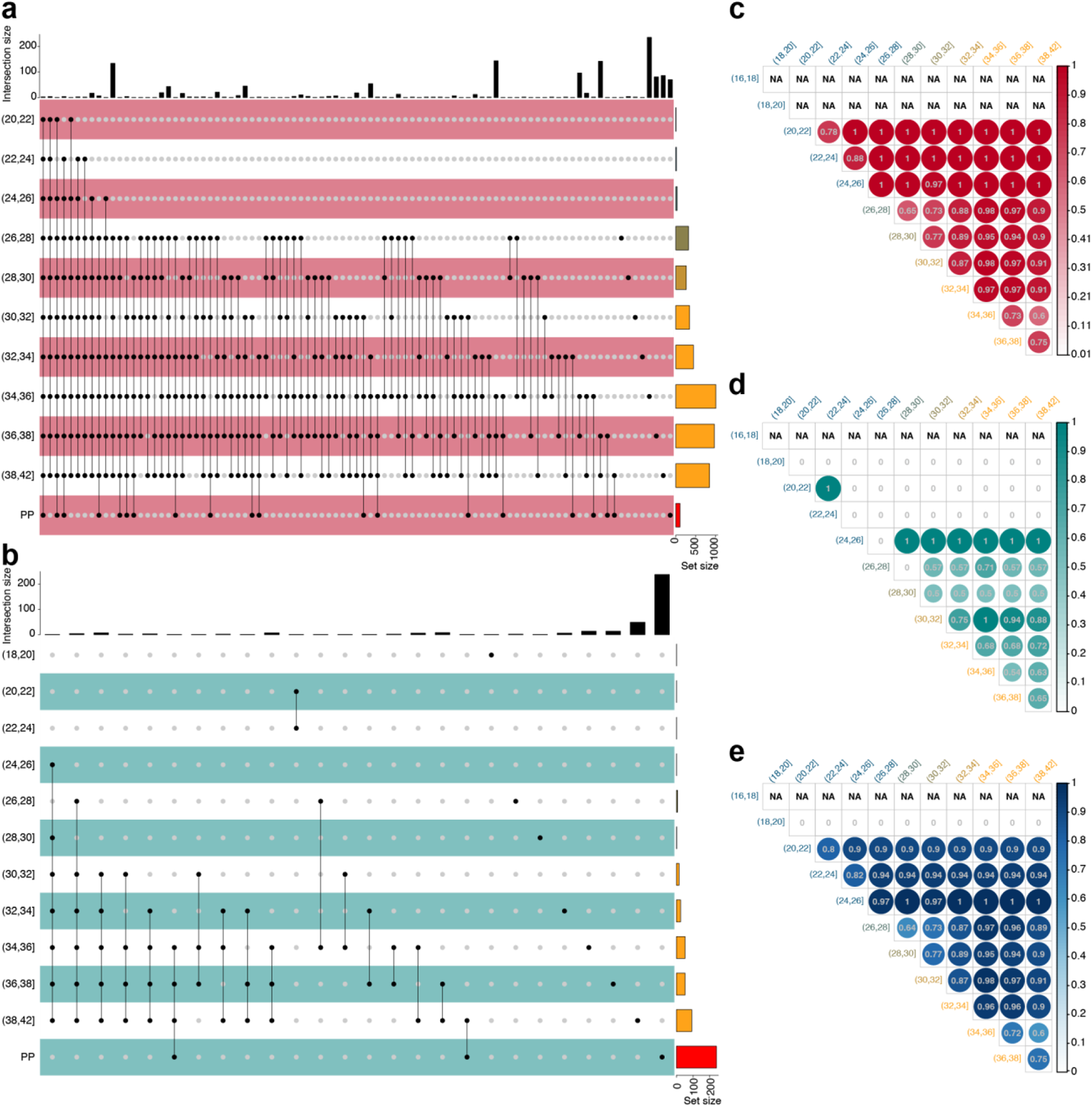
Altered metabolic peaks in different GA ranges and their consistency. (**a**) The upregulated metabolic peaks in different GA ranges and their overlap shown by upset plot. (**b**) The downregulated metabolic peaks in different GA ranges and their overlap shown by the upset plot. (**c**) The upregulated metabolic peaks in different GA ranges and their overlap with next GA ranges. (**d**) The downregulated metabolic peaks in different GA ranges and their overlap with next GA ranges. (**e**) All altered metabolic peaks in different GA ranges and their overlap with next GA ranges.

**Figure S6.**
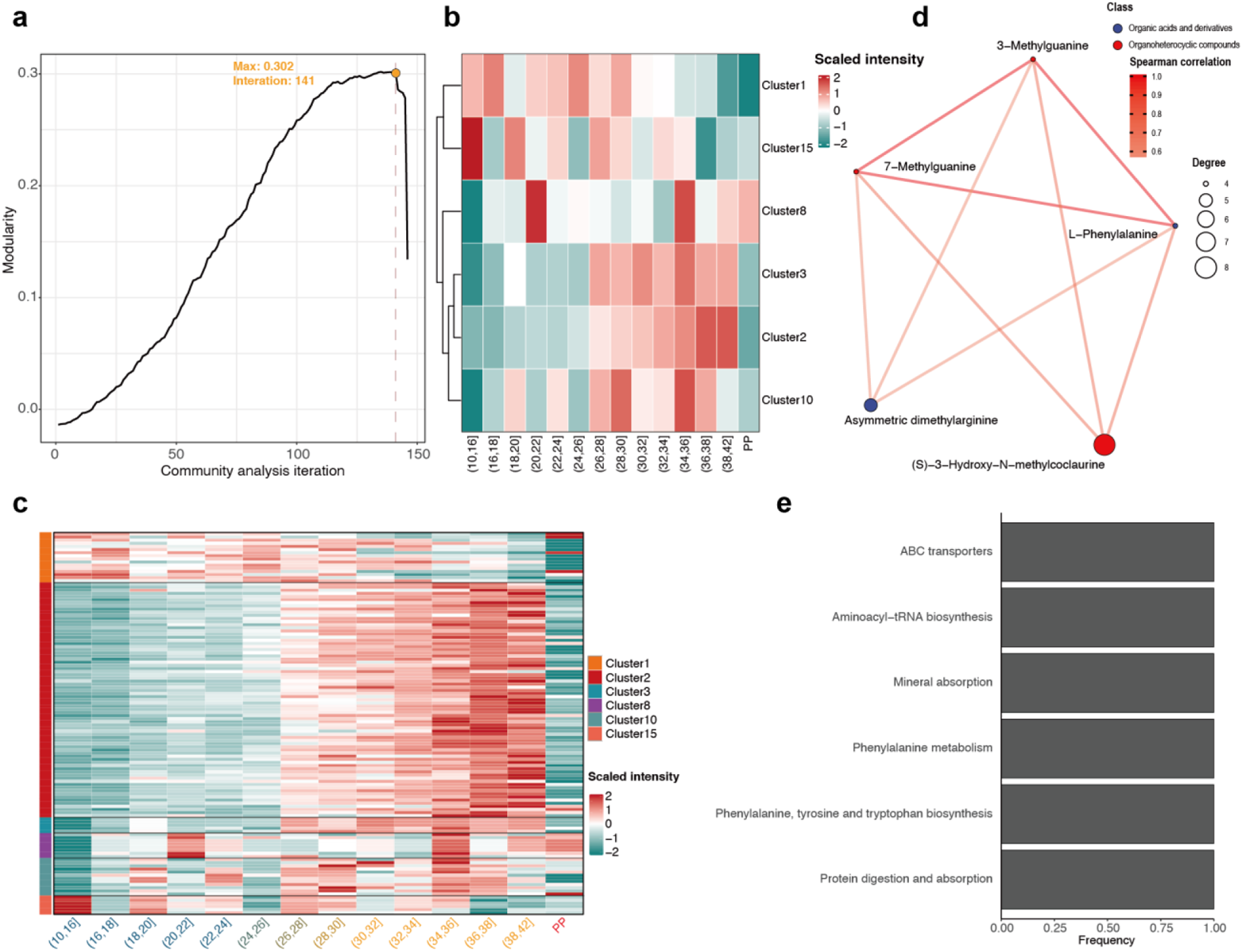
Community analysis for correlation network from PIUMet. (**a**) The maximum modularity observed in our correlation network community analysis was 0.302 at iteration 141 of community pruning. There were 150 total iterations of community analysis. (**b**) Heatmap to show the changes of 6 clusters during pregnancy at cluster level. (**c**) Heatmap to show the changes of 6 clusters during pregnancy at metabolite (metabolic peak) level. (**d**) Correlation network of cluster 3. (**e**) Bar plot to show the frequency of pathways of all metabolites in cluster 3 belong to.

**Figure S7.**
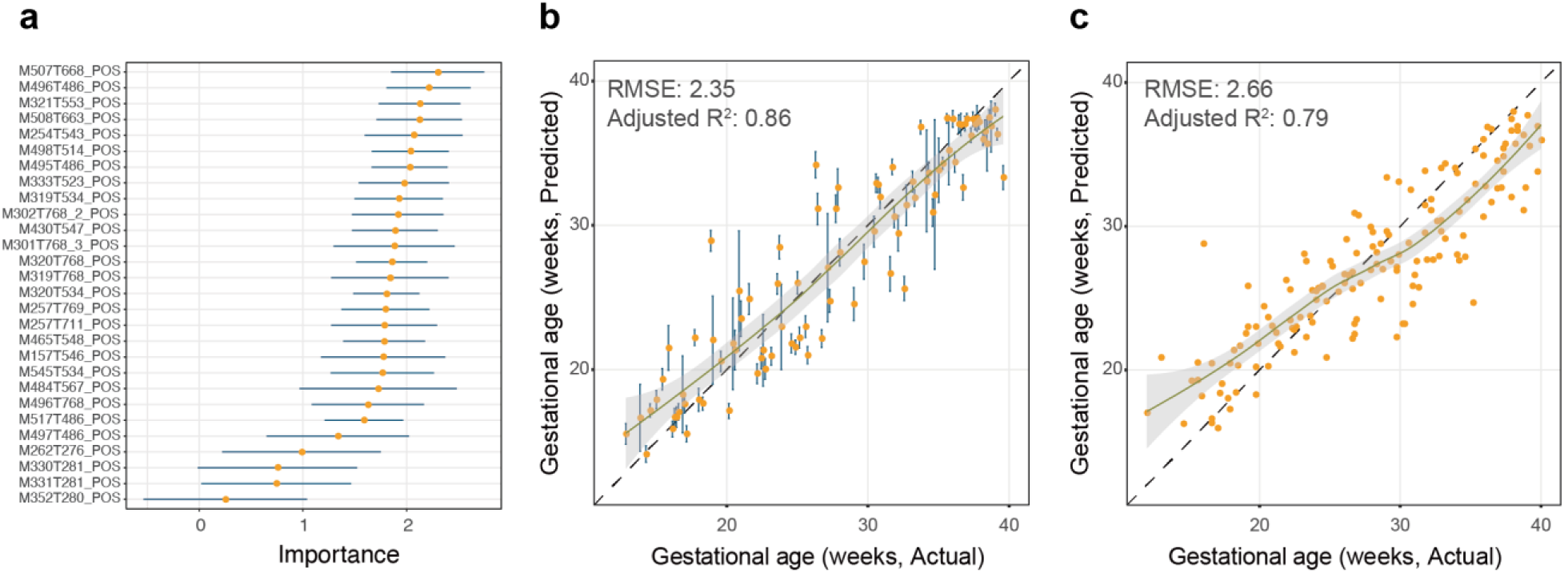
Urine metabolome can be used to predict gestational age. (**a**) 28 metabolic peaks were selected as potential biomarkers based on the Boruta algorithm for Random Forest prediction model. (**b-c**) Using 28 metabolic peak biomarkers to build prediction model, gestational age predicted by 28 metabolic peaks (Y-axis) is highly concordant to clinical values determined by the standard of care (first-trimester ultrasound, x-axis) in internal validation (**b**) and external validation dataset (**c**).

**Figure S8.**
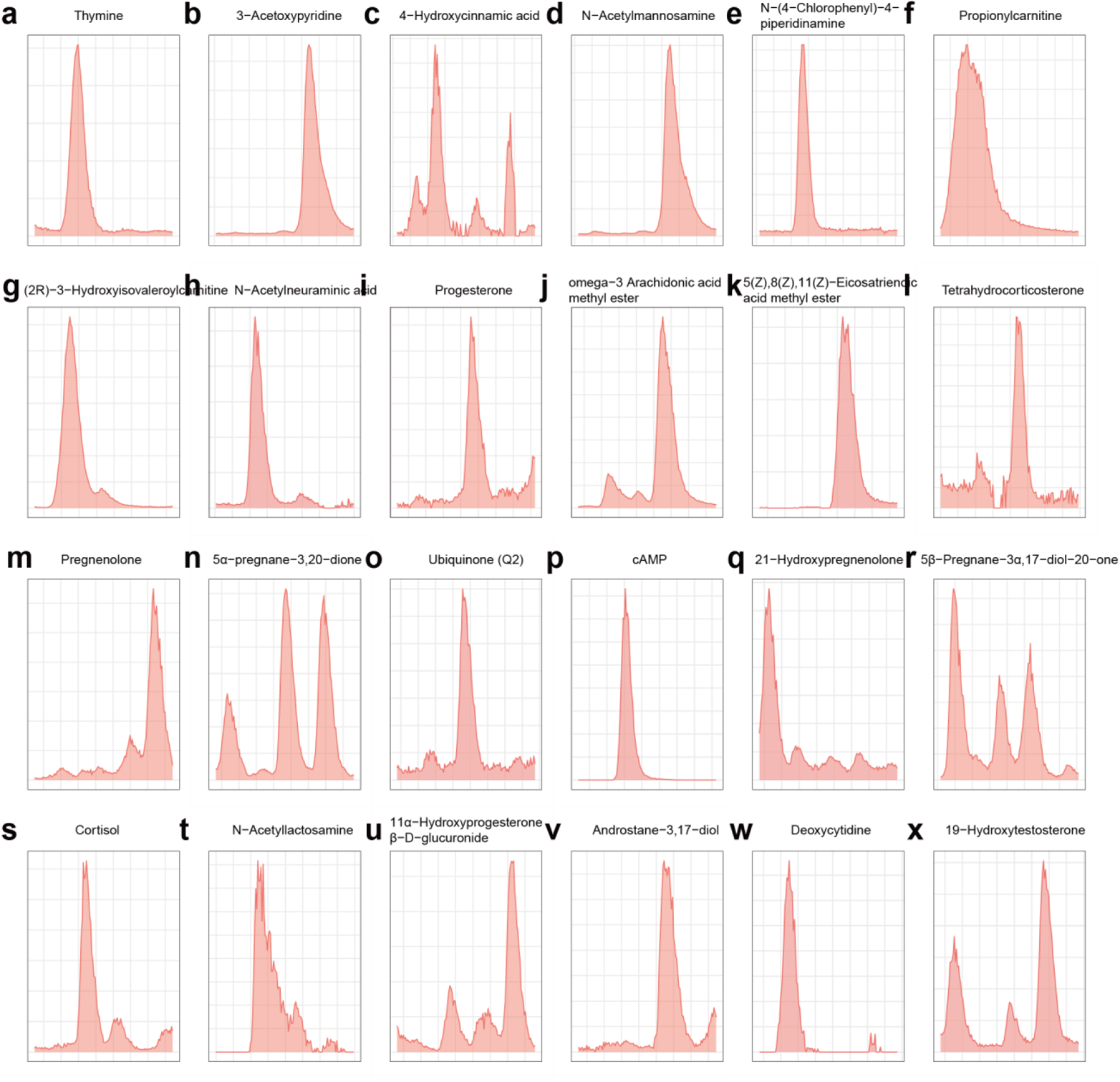
The peak shapes of 24 metabolite biomarkers in GA and sampling time to delivery models.

**Figure S9.**
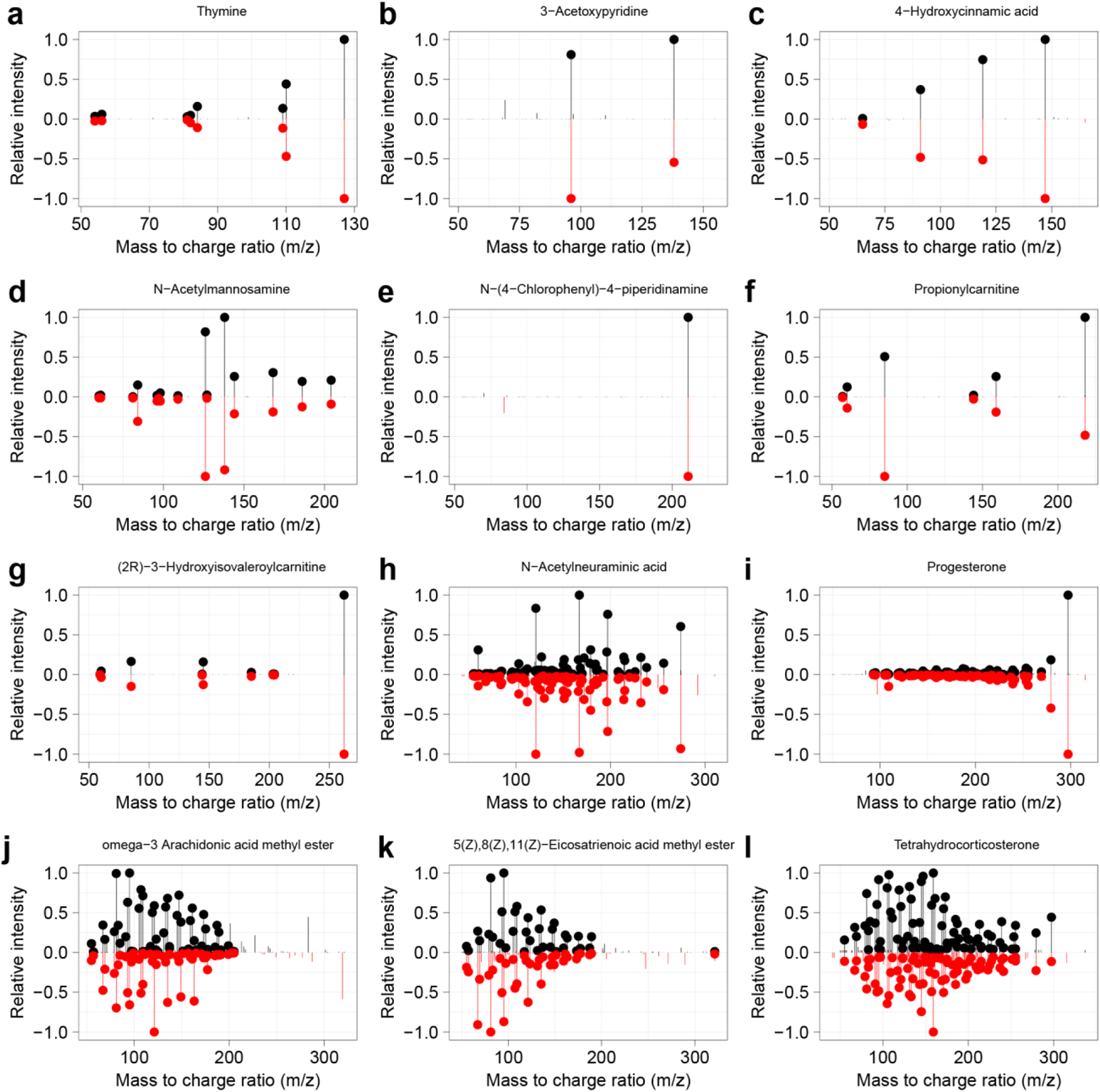
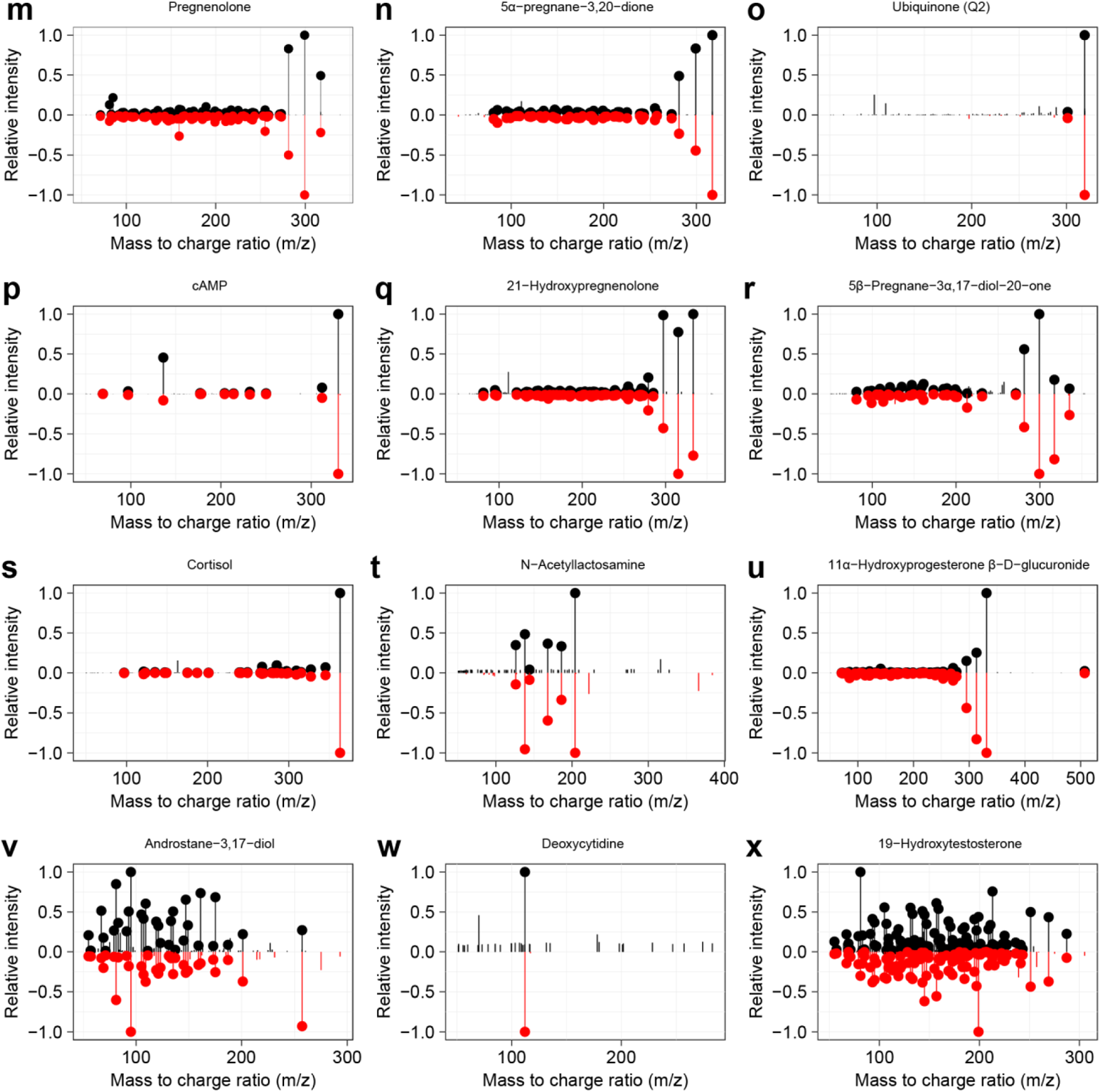
The MS^2^ spectra match of 24 metabolite biomarkers in gestational age and sampling time to delivery models.

**Figure S10.**
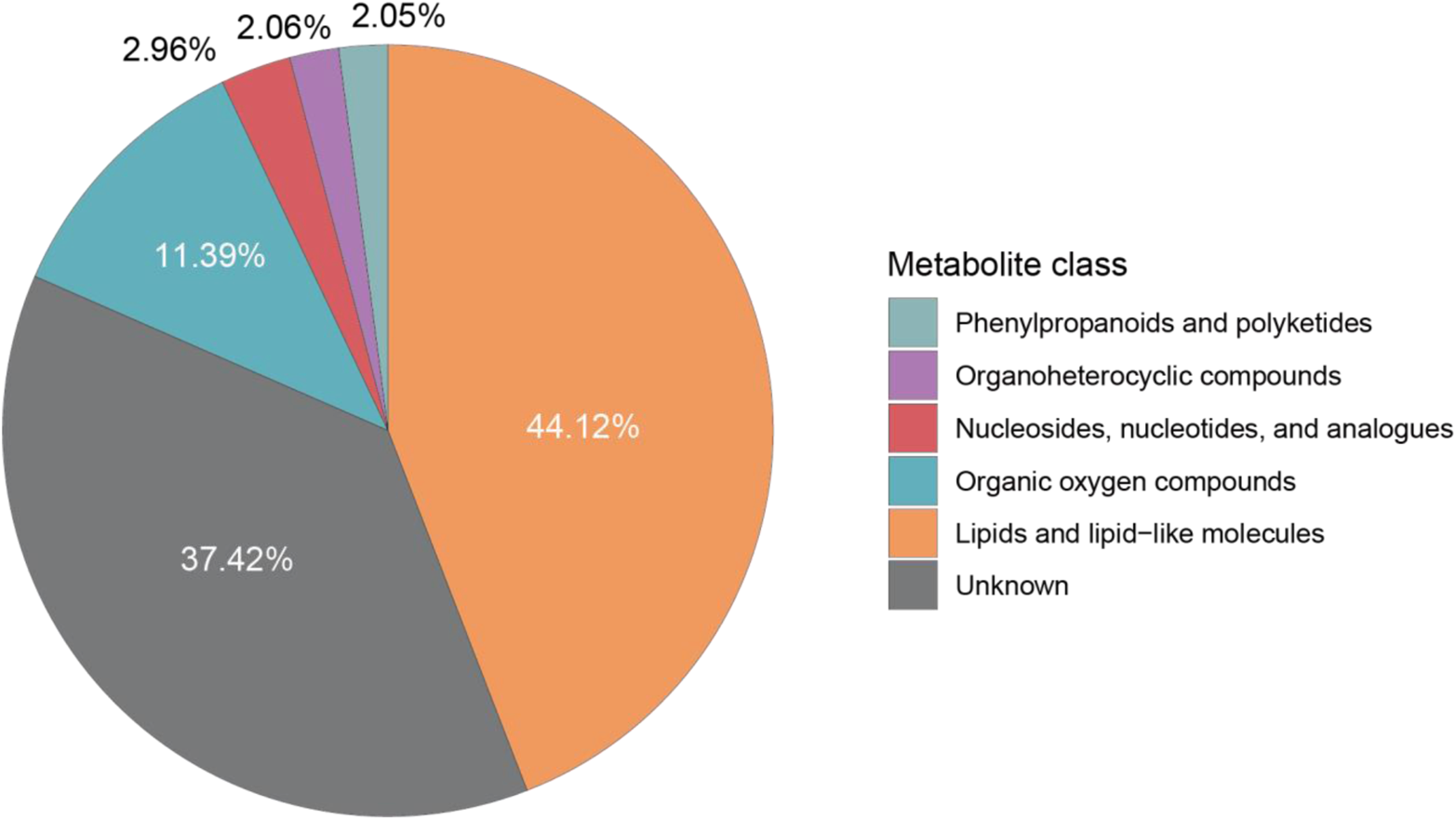
Importance ratio of different chemical class in prediction model for gestation age.

**Figure S11.**
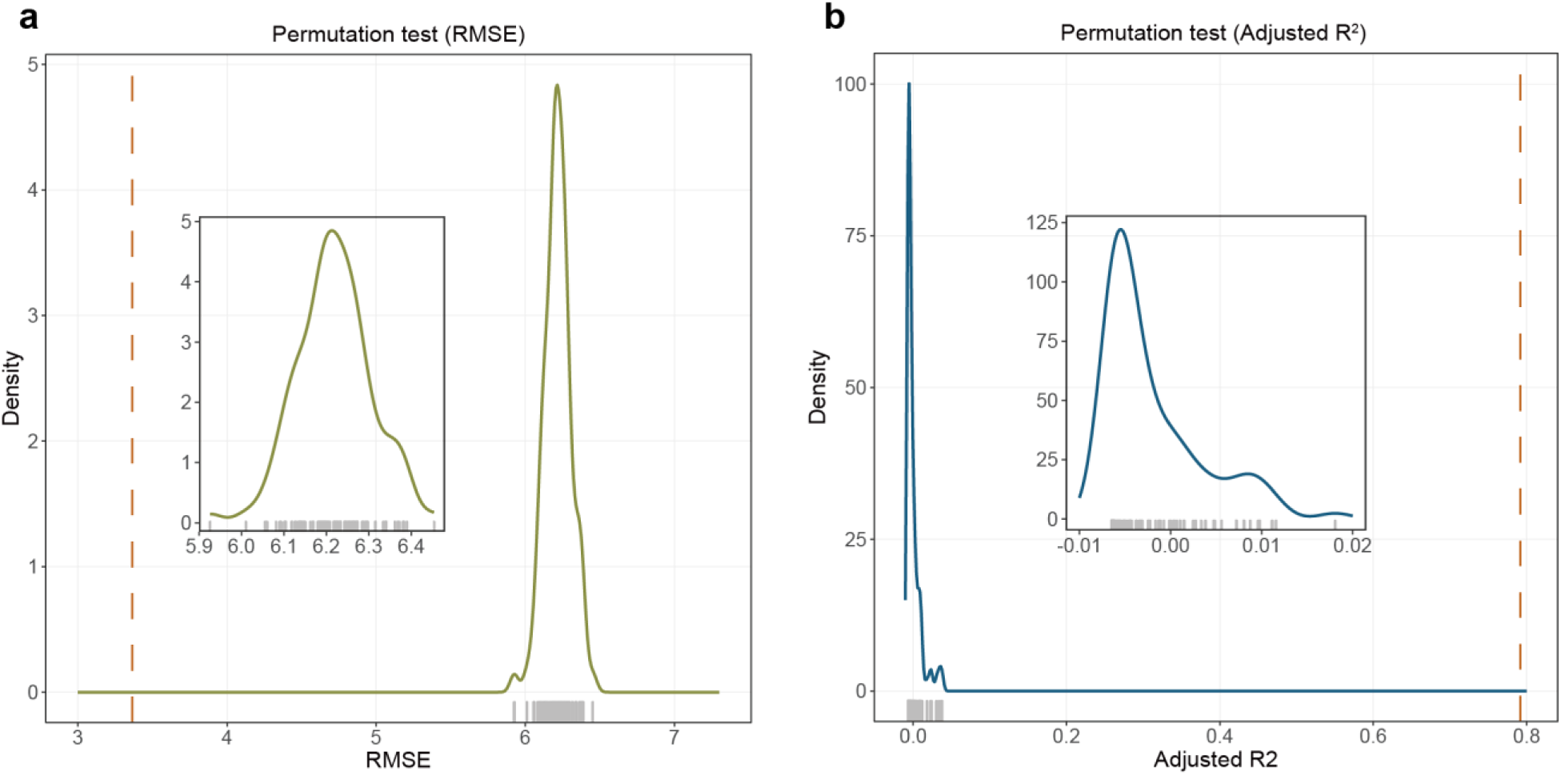
Permutation test for prediction module for gestational age. (a) Null distribution of RMSE values. (b) Null distribution of adjusted R^2^ values.

**Figure S12.**
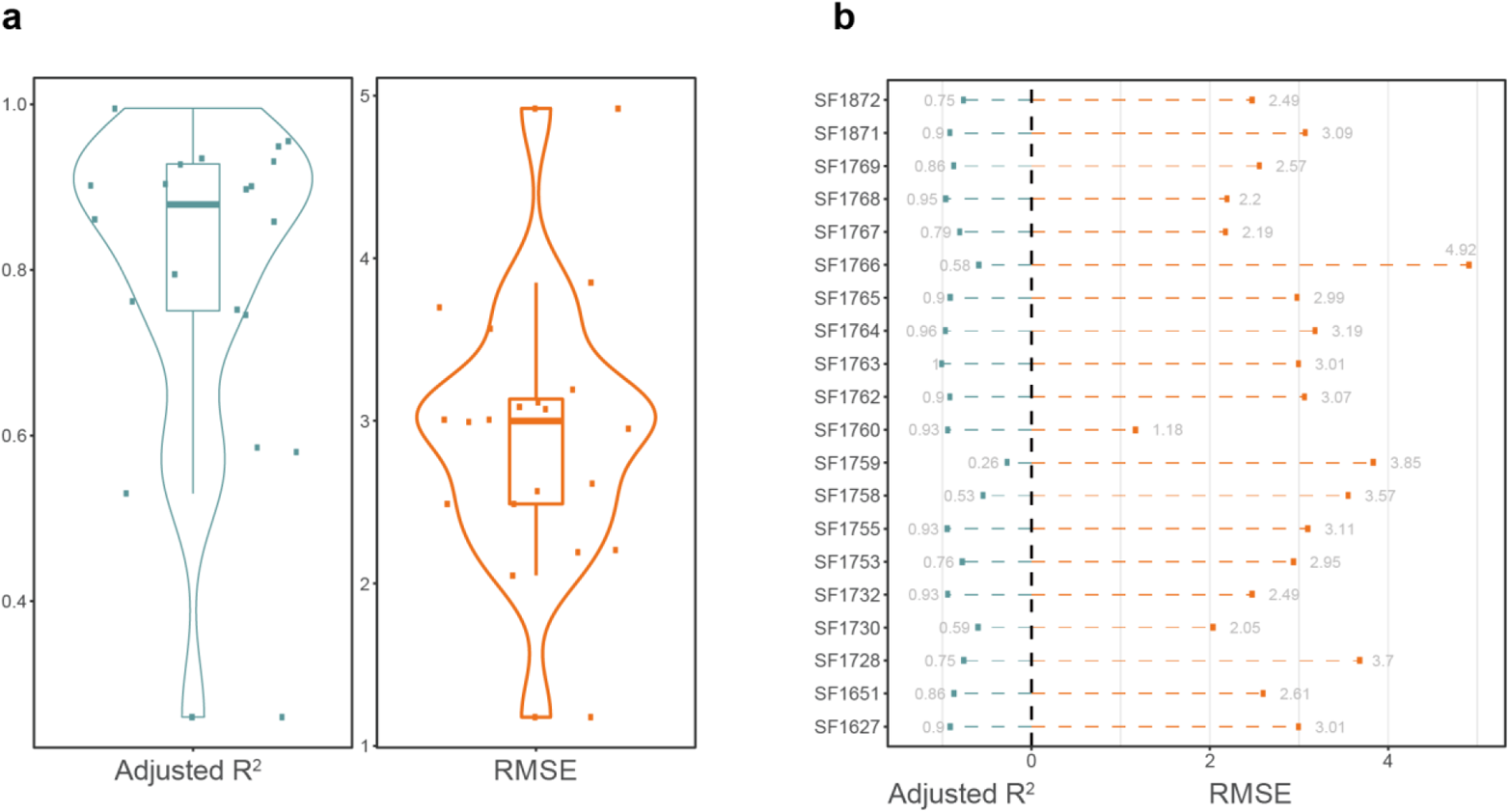
Gestational age prediction result for each participant. (**a**) Distribution of RMSE and adjusted R^2^ for each participant in the validation dataset. (**b**) RMSE and adjusted R^2^ for each participant in the validation dataset.

**Figure S13.**
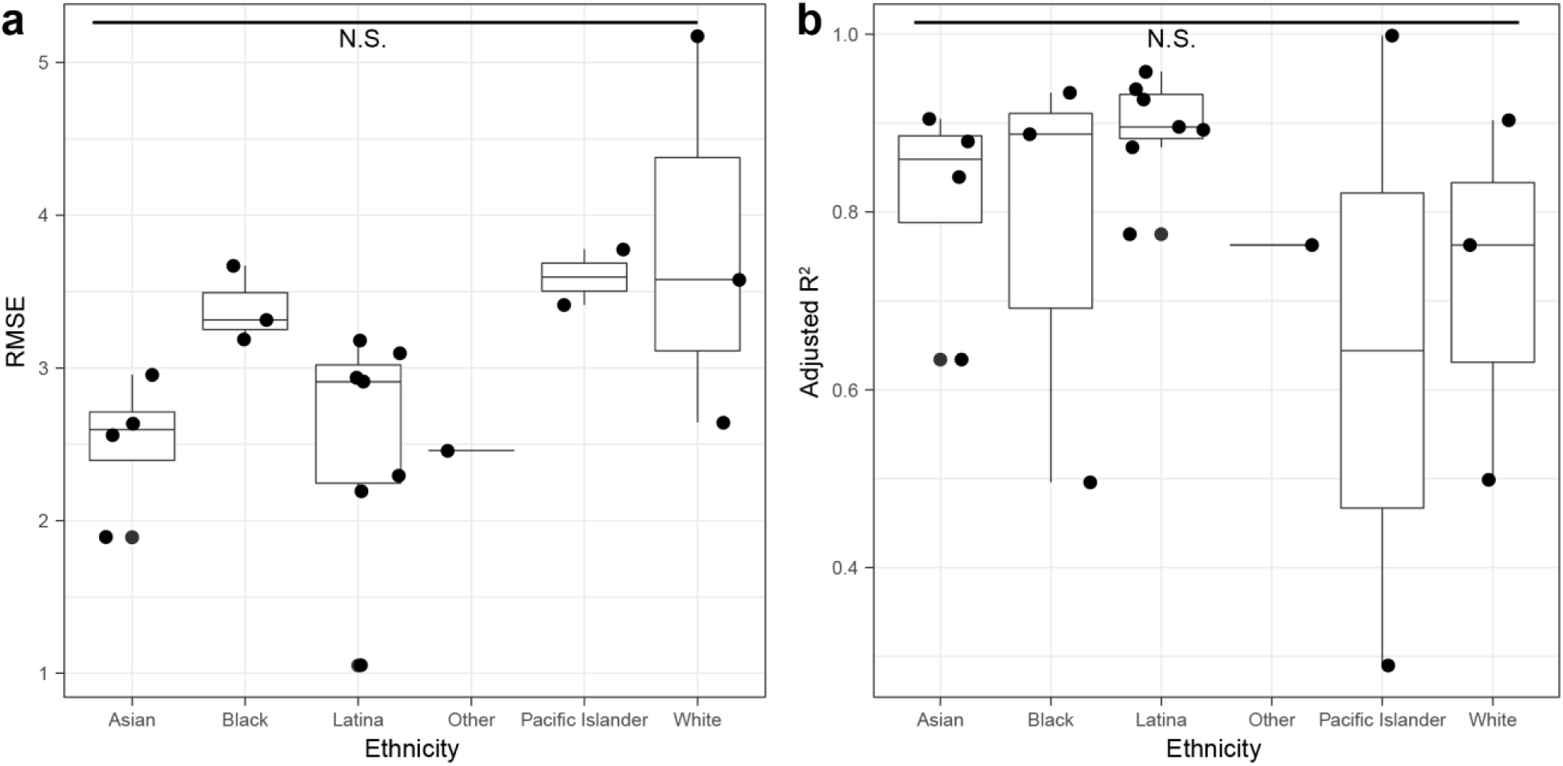
Prediction accuracy with characteristics in GA model. (**a**) RMSE in different ethnicities are not significant. (**b**) Adjusted R2 in different ethnicities are not significant.

**Figure S14.**
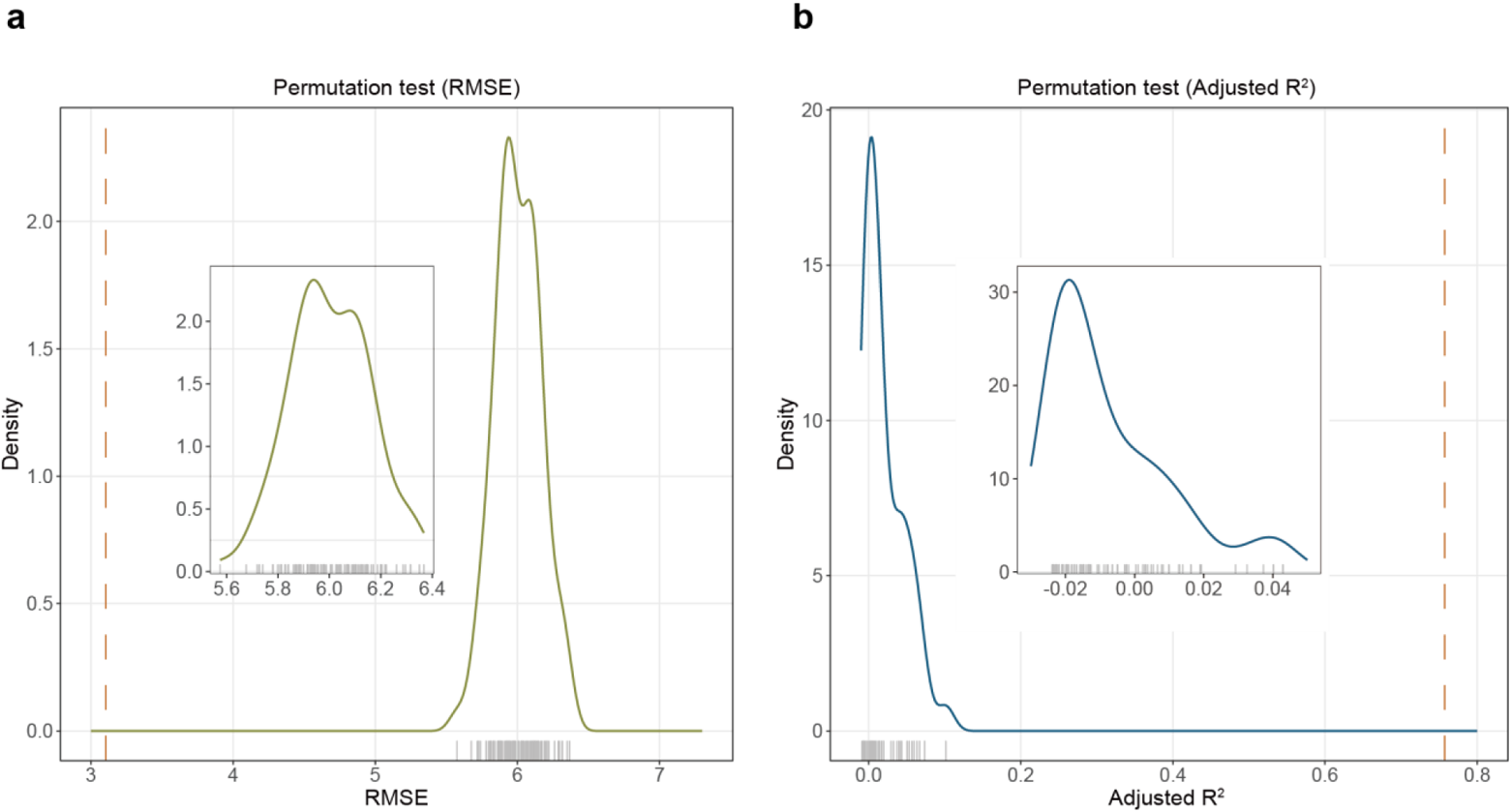
Permutation test for prediction module for sampling time to delivery. (**a**) Null distribution of RMSE values. (**b**) Null distribution of adjusted R^2^ values.

**Figure S15.**
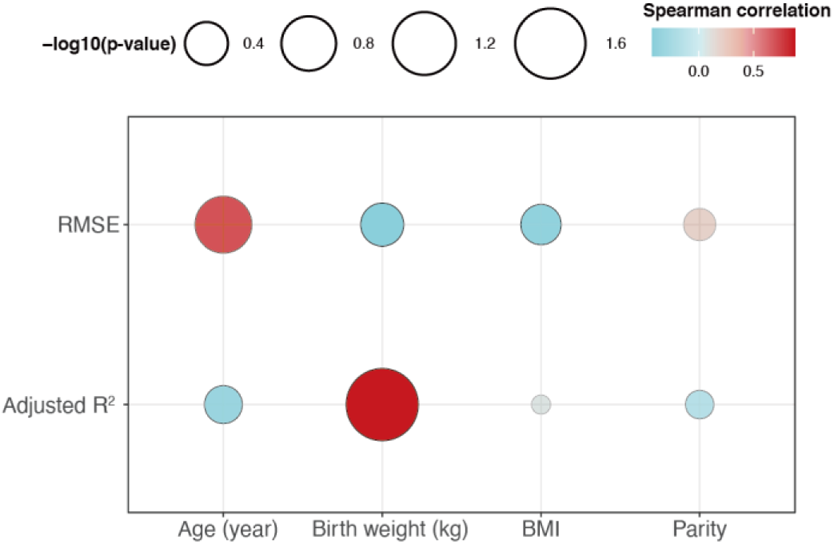
The continuous characteristics have no effect on sampling to delivery prediction accuracy.

**Figure S16.**
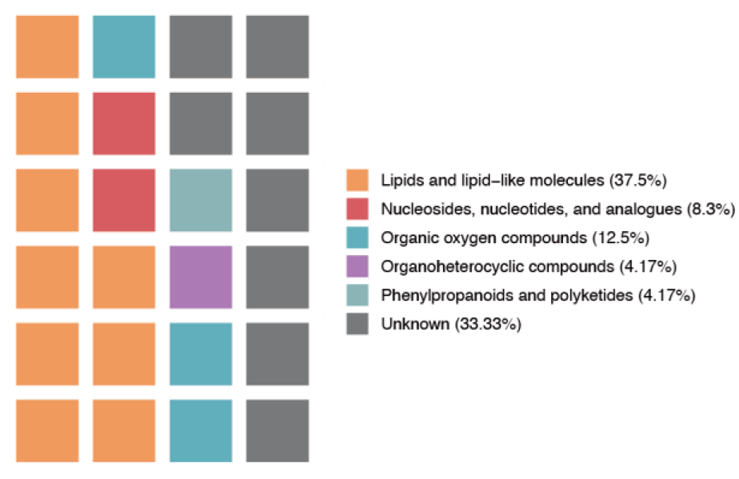
Chemical class of 24 metabolite biomarkers in GA and sampling time to delivery models.

**Figure S17.**
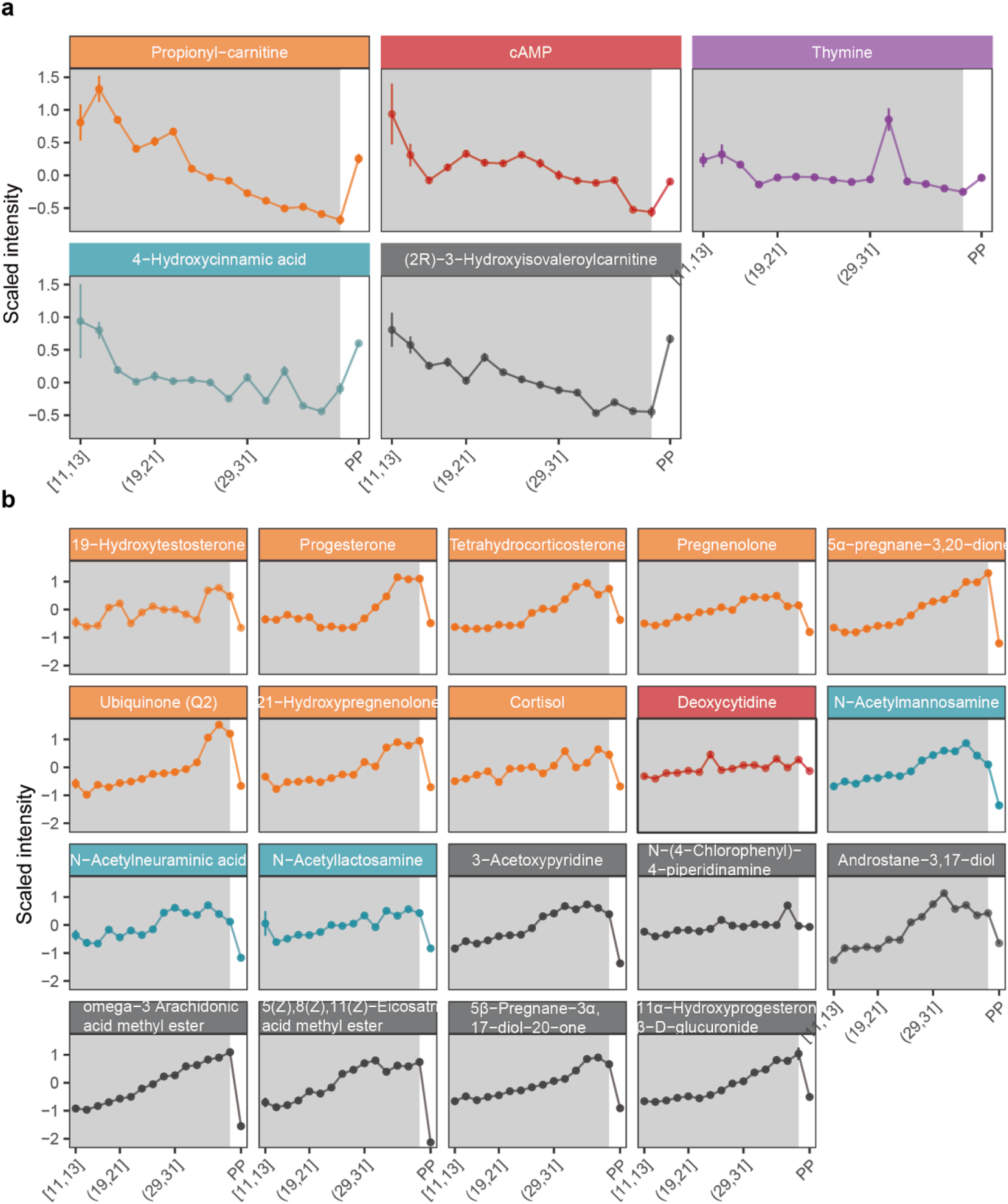
The trends of 24 metabolite markers during pregnancy. (**a**) Five metabolite markers decrease during pregnancy and increase after childbirth. (**b**) Nineteen metabolite markers increase during pregnancy and decrease after childbirth.

**Figure S18.**
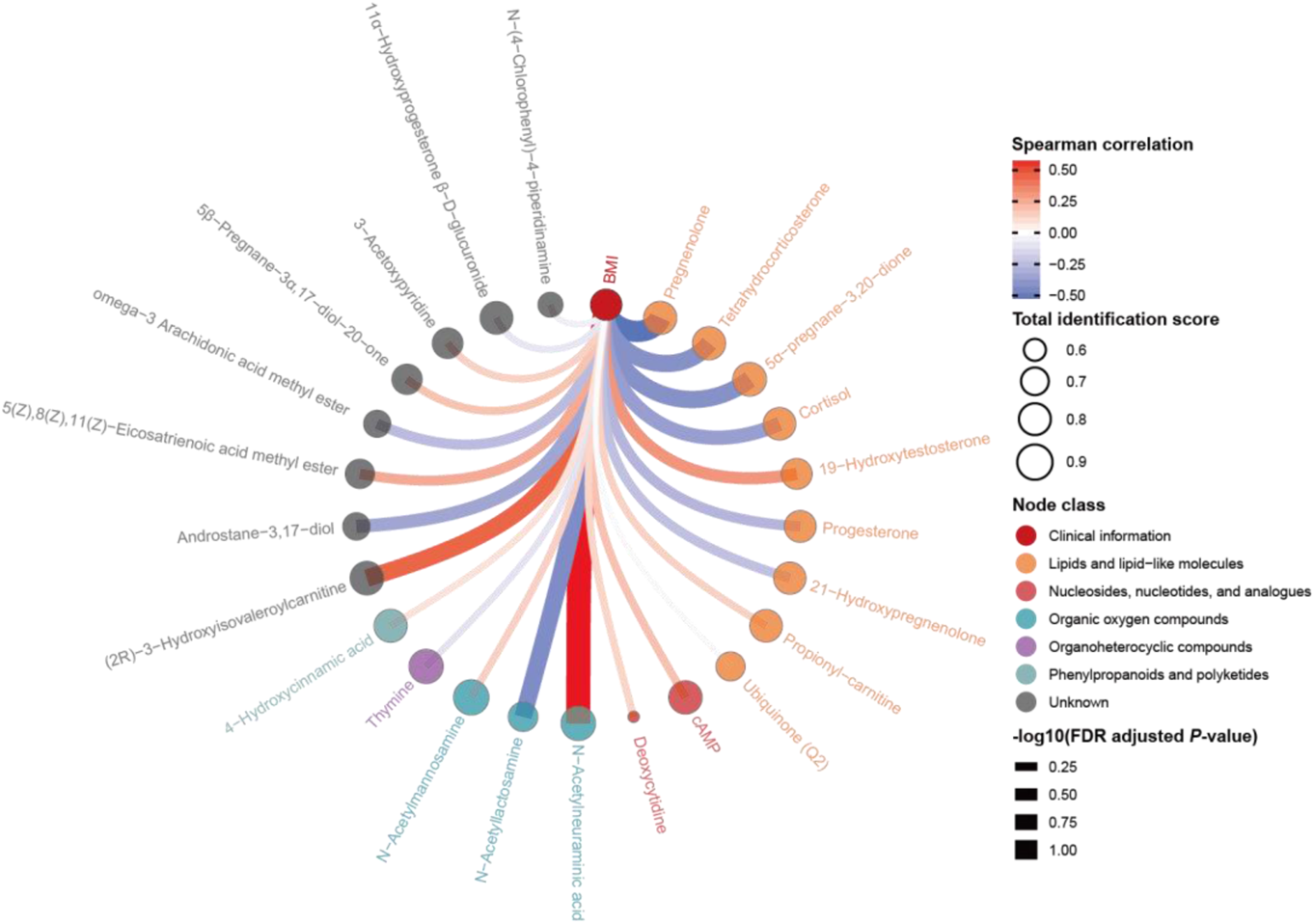
Correlation network between BMI and metabolites markers.

**Figure S19.**
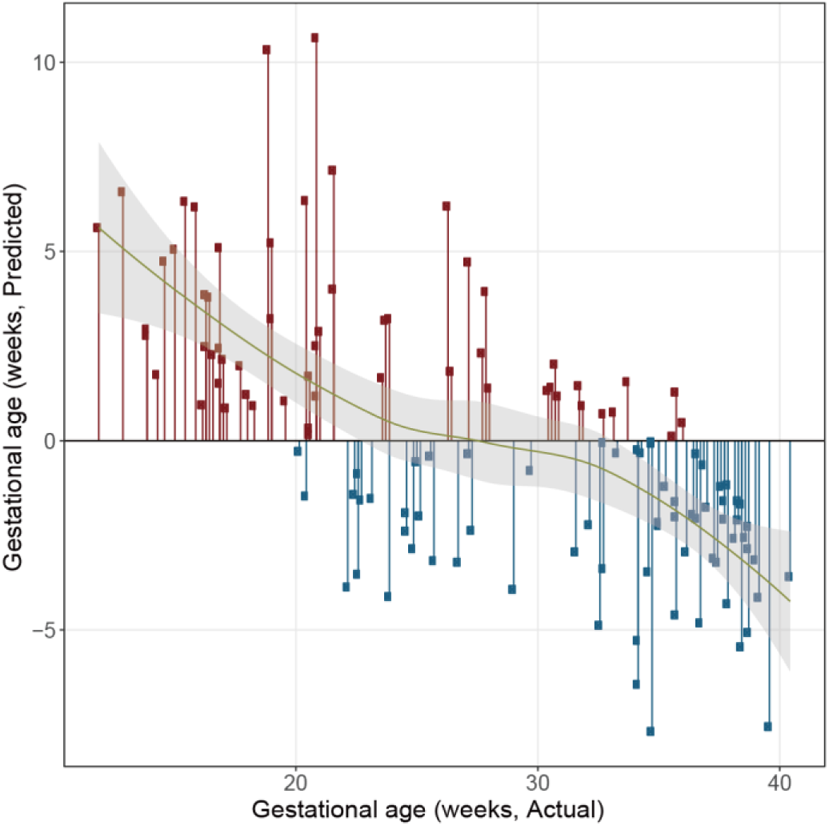
Prediction error using Random Forest to predict gestational age.

## ACKNOWLEDGEMENT

We thank the dedicated pregnant women that with their generous participation throughout their pregnancy period and at postpartum made this study possible. this work was funded by the UCSF California Preterm Birth Initiative (PTBi-CA).

## AUTHOR CONTRIBUTION

LR and LJP organized and contributed to the collection of pregnancy samples. LL, SC, XS, LJP, LR and MS conceptualized the study and analysis plan. SC, LL, MA, XS and HZ processed and analyzed the samples, XS, SC and LL analyzed the data, SC, XS, LL, LJP, LR, and MS wrote the manuscript, and all authors contributed to the reviewing and editing of the manuscript.

## Data and materials availability

All data associated with this study are present in the paper or the Supplementary Materials.

