## Supplemental for "Longitudinal Urine Metabolic Profiling and Gestational Age Prediction in Pregnancy"

**Table S1.** The detailed information for all the participants in SMART-D study.

**Table S2.** 28 potential metabolic peak markers to predict gestational age.

**Table S3.** The detailed information of 24 metabolite biomarkers to predict gestational age and sampling time to delivery.

**Table S4.** Prediction result for each participant in prediction gestational age model.

**Table S5.** The detailed information for each urine sample in SMART-D study.

**Table S6.** Parameters for MS raw data conversion using ProteoWizard.

| Parameter | Setting |
|---|---|
| Output format | mzXML |
| Binary encoding precision | 64-bit |
| Write index | Check |
| Use zlib compression | Check |
| TPP compatibility | Check |
| Package in gzip | Uncheck |
| Filter | Peak Picking |

**Table S7.** All software and R packages used in data analysis.

| **Tools** | **Source** | **Source** | **Version** |
|---|---|---|---|
| R | R Core Team | https://www.r-project.org/ | 3.6.0 |
| RStudio | RStudio, PBC | https://rstudio.com/ | 1.2.5019 |
| plyr | CRAN | https://cran.r-project.org/web/packages/plyr/ | 1.8.5 |
| stringr | CRAN | https://cran.r-project.org/web/packages/stringr/ | 1.4.0 |

| | | | |
|---|---|---|---|
| dplyr | CRAN | https://cran.r-project.org/web/packages/dplyr/ | 0.8.3 |
| purrr | CRAN | https://cran.r-project.org/web/packages/purrr/ | 0.3.3 |
| readr | CRAN | https://cran.r-project.org/web/packages/readr/ | 1.3.1 |
| readxl | CRAN | https://cran.r-project.org/web/packages/readxl/ | 1.3.1 |
| tidyr | CRAN | https://cran.r-project.org/web/packages/tidyr/ | 1.0.0 |
| tibble | CRAN | https://cran.r-project.org/web/packages/tibble/ | 2.1.3 |
| ggplot2 | CRAN | https://cran.r-project.org/web/packages/ggplot2/ | 3.2.1 |
| ggsci | CRAN | https://cran.r-project.org/web/packages/ggsci/ | 2.9 |
| patchwork | CRAN | https://cran.r-project.org/web/packages/patchwork/ | 1.0.0 |
| igraph | CRAN | https://cran.r-project.org/web/packages/igraph/ | 1.2.4.2 |
| Boruta | CRAN | https://cran.r-project.org/web/packages/Boruta/ | 7.0.0 |
| randomForest | CRAN | https://cran.r-project.org/web/packages/randomForest/ | 4.6-14 |
| e0171 | CRAN | https://cran.r-project.org/web/packages/e1071/ | 1.7-3 |
| Mfuzz | Bioconductor | https://bioconductor.org/packages/devel/bioc/html/Mfuzz.html | 3.12 |
| ggraph | CRAN | https://cran.r-project.org/web/packages/ggraph/ | 2.0.3 |
| ggplotify | CRAN | https://cran.r-project.org/web/packages/ggplotify/index.html | 0.0.5 |
| xcms | Bioconductor | https://www.bioconductor.org/packages/release/bioc/html/xcms.html | 3.10.1 |
| impute | Bioconductor | https://bioconductor.org/packages/release/bioc/html/impute.html | 1.62.0 |

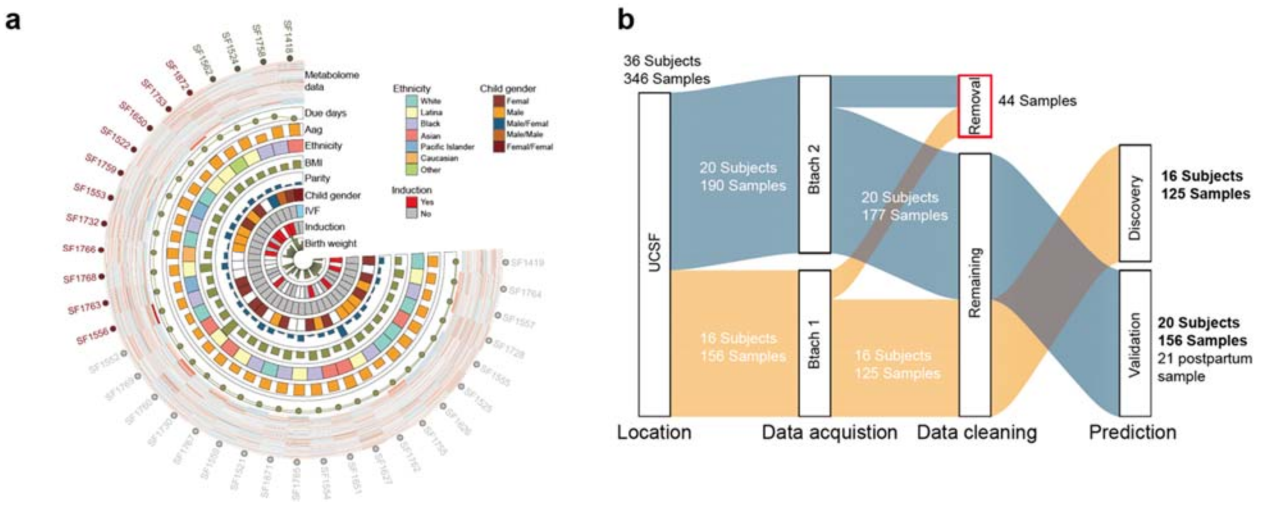

**Figure S1. Study design of SMART-D study.** (**a**) The metabolome and clinical information of 36 participants in SMART-D study. (**b**) Sample collection, data acquisition, data processing and analysis design for our study. UCSF (University of California, San Francisco), ZSFGH (Zuckerberg San Francisco General Hospital).

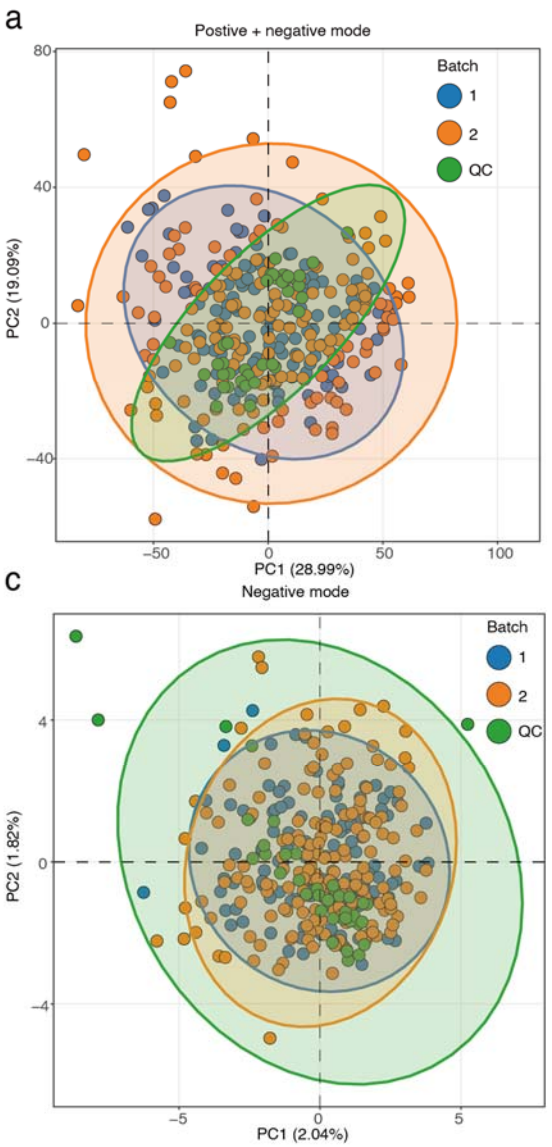


**Figure S2. Data quality of urine metabolomics data.** (**a**) Positive and negative mode data. (**b**) Only positive mode data. (**c**) Only negative data.

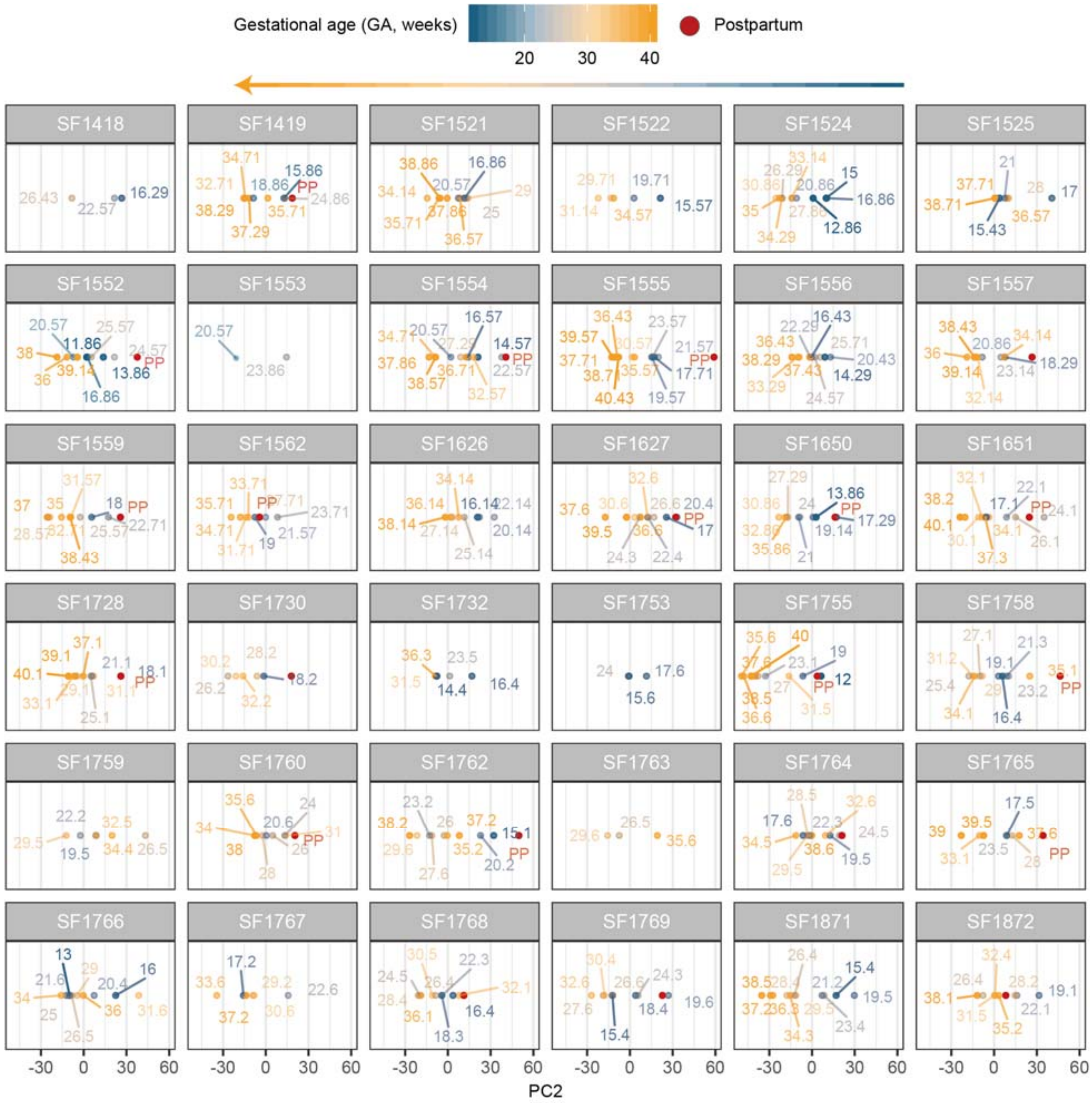


**Figure S3. PCA score plot for each participant in SMART-D study.**

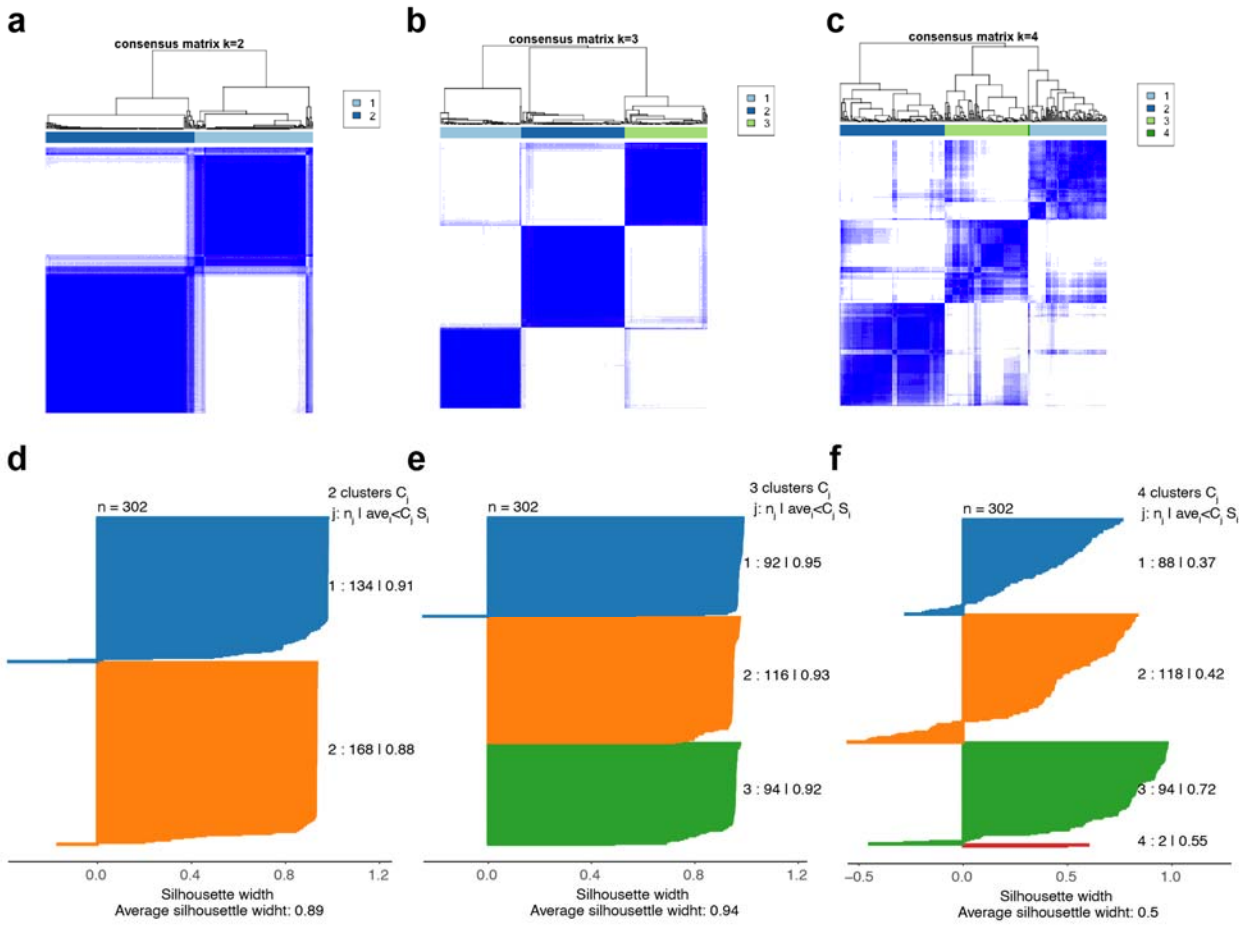


**Figure S4. K means consensus-clustering for urine metabolome in 302 samples.** The heatmaps of consensus matrix for k = 2 (**a**), k = 3 (**b**) and k = 4 (**c**) clusters based on 1,000 resampled datasets. The silhouette plots for k = 2 (**d**), k = 3 (**e**) and k = 4 (**f**) clusters.

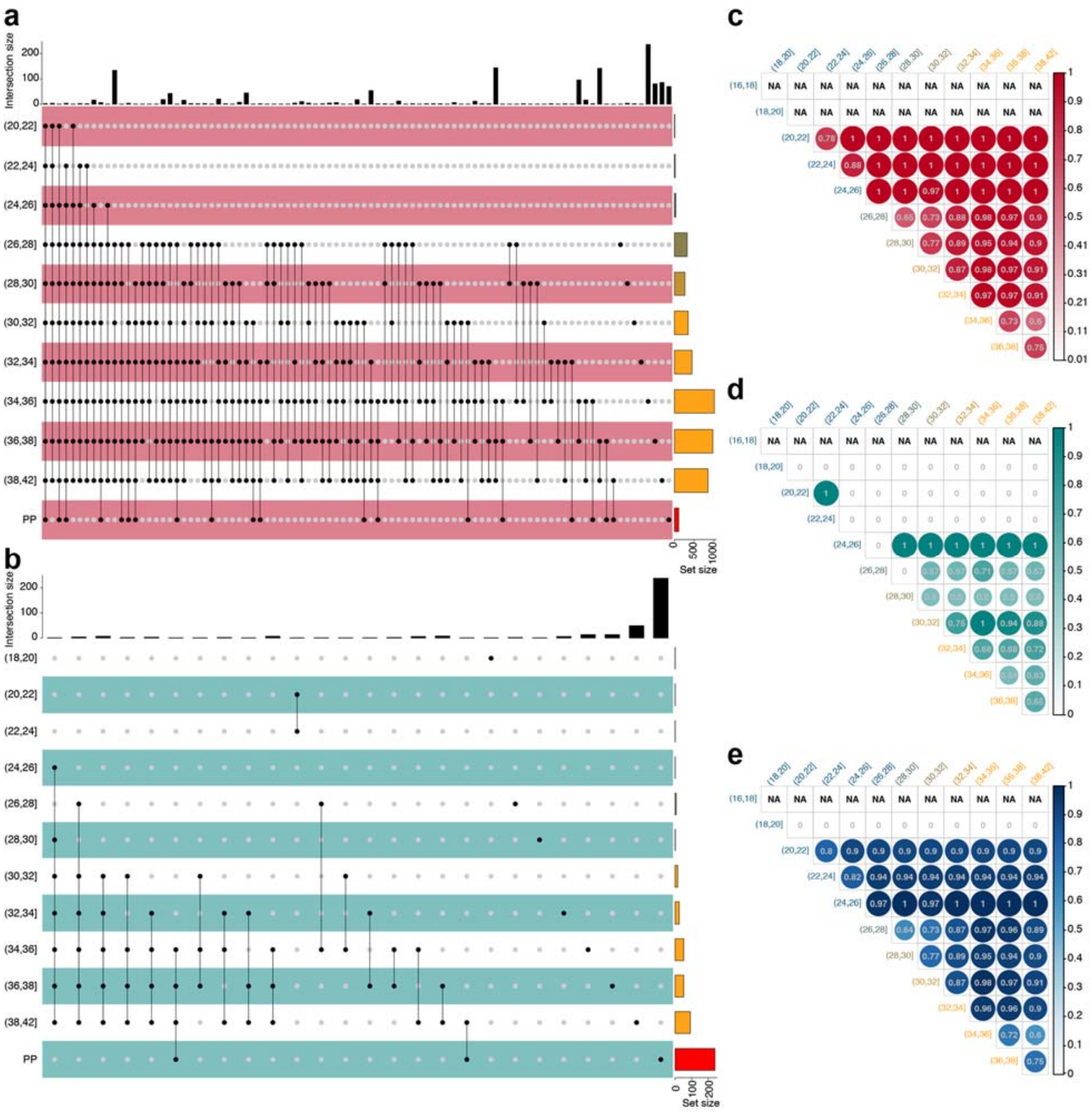


**Figure S5. Altered metabolic peaks in different GA ranges and their consistency.** (**a**) The upregulated metabolic peaks in different GA ranges and their overlap shown by upset plot. (**b**) The downregulated metabolic peaks in different GA ranges and their overlap shown by the upset plot. (**c**) The upregulated metabolic peaks in different GA ranges and their overlap with next GA ranges. (**d**) The downregulated metabolic peaks in different GA ranges and their overlap with next GA ranges. (**e**) All altered metabolic peaks in different GA ranges and their overlap with next GA ranges.

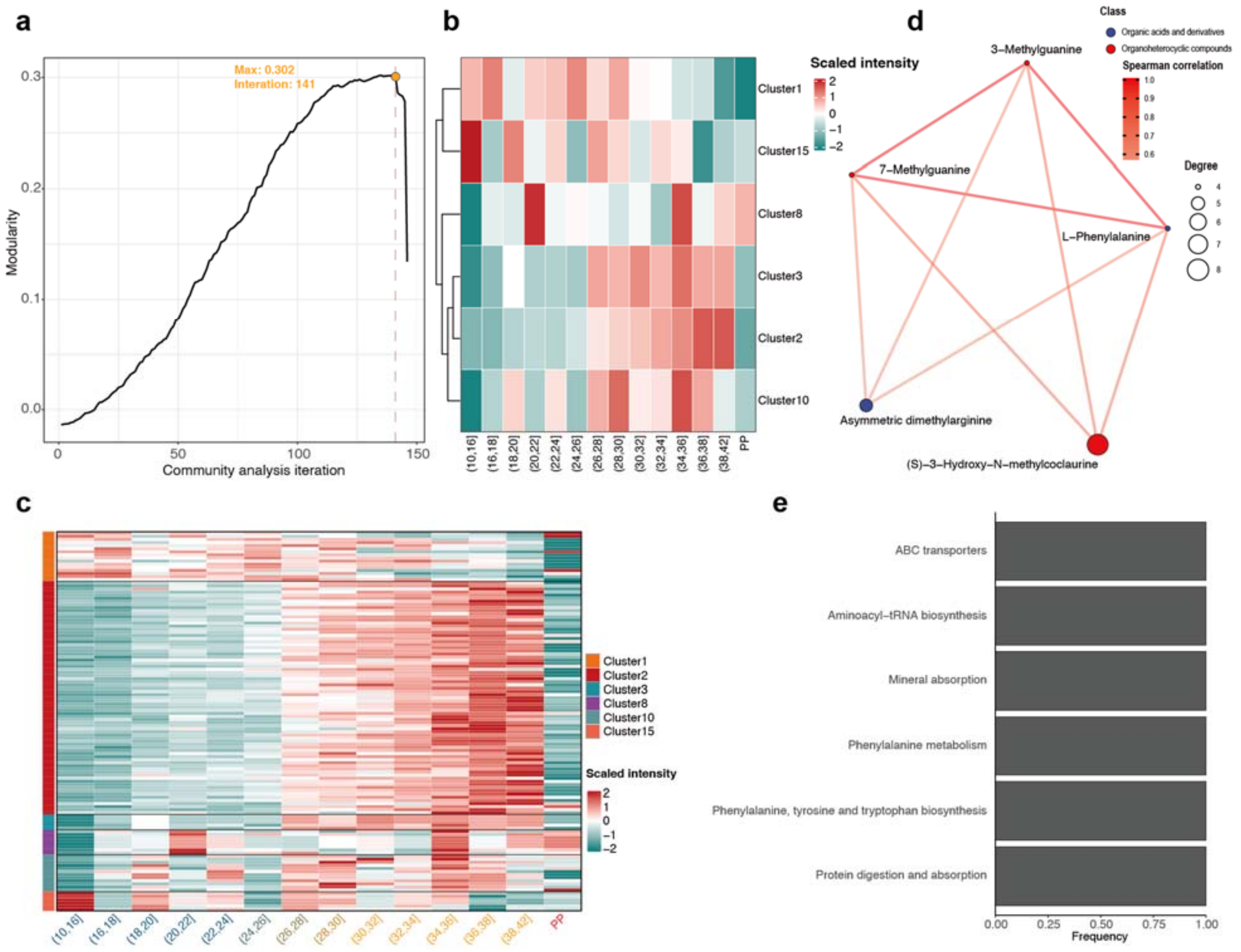

**Figure S6. Community analysis for correlation network from PIUMet.** (**a**) The maximum modularity observed in our correlation network community analysis was 0.302 at iteration 141 of community pruning. There were 150 total iterations of community analysis. (**b**) Heatmap to show the changes of 6 clusters during pregnancy at cluster level. (**c**) Heatmap to show the changes of 6 clusters during pregnancy at metabolite (metabolic peak) level. (**d**) Correlation network of cluster 3. (**e**) Bar plot to show the frequency of pathways of all metabolites in cluster 3 belong to.

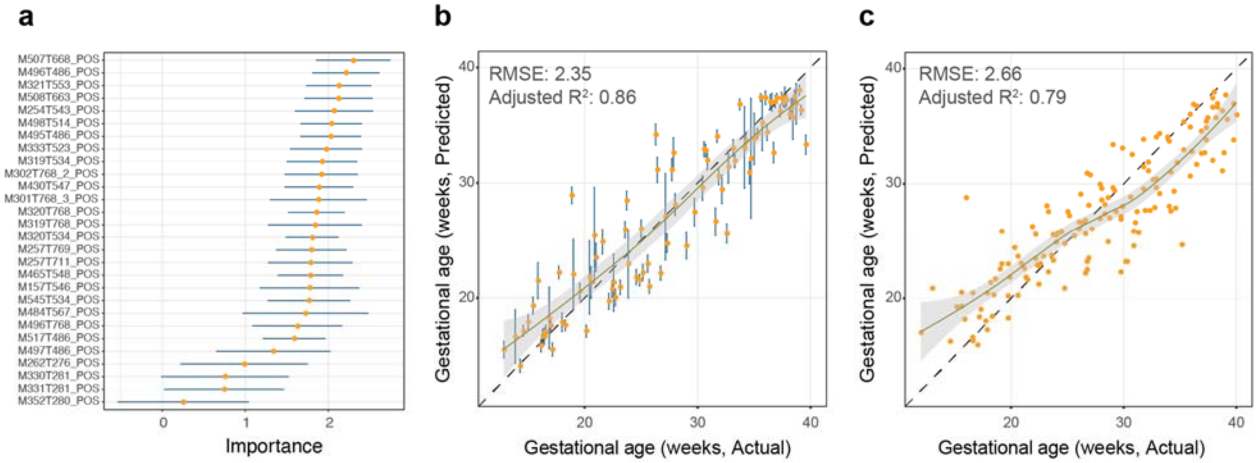


**Figure S7. Urine metabolome can be used to predict gestational age.** (**a**) 28 metabolic peaks were selected as potential biomarkers based on the Boruta algorithm for Random Forest prediction model. (**b-c**) Using 28 metabolic peak biomarkers to build prediction model, gestational age predicted by 28 metabolic peaks (Y-axis) is highly concordant to clinical values determined by the standard of care (first-trimester ultrasound, x-axis) in internal validation (**b**) and external validation dataset (**c**).

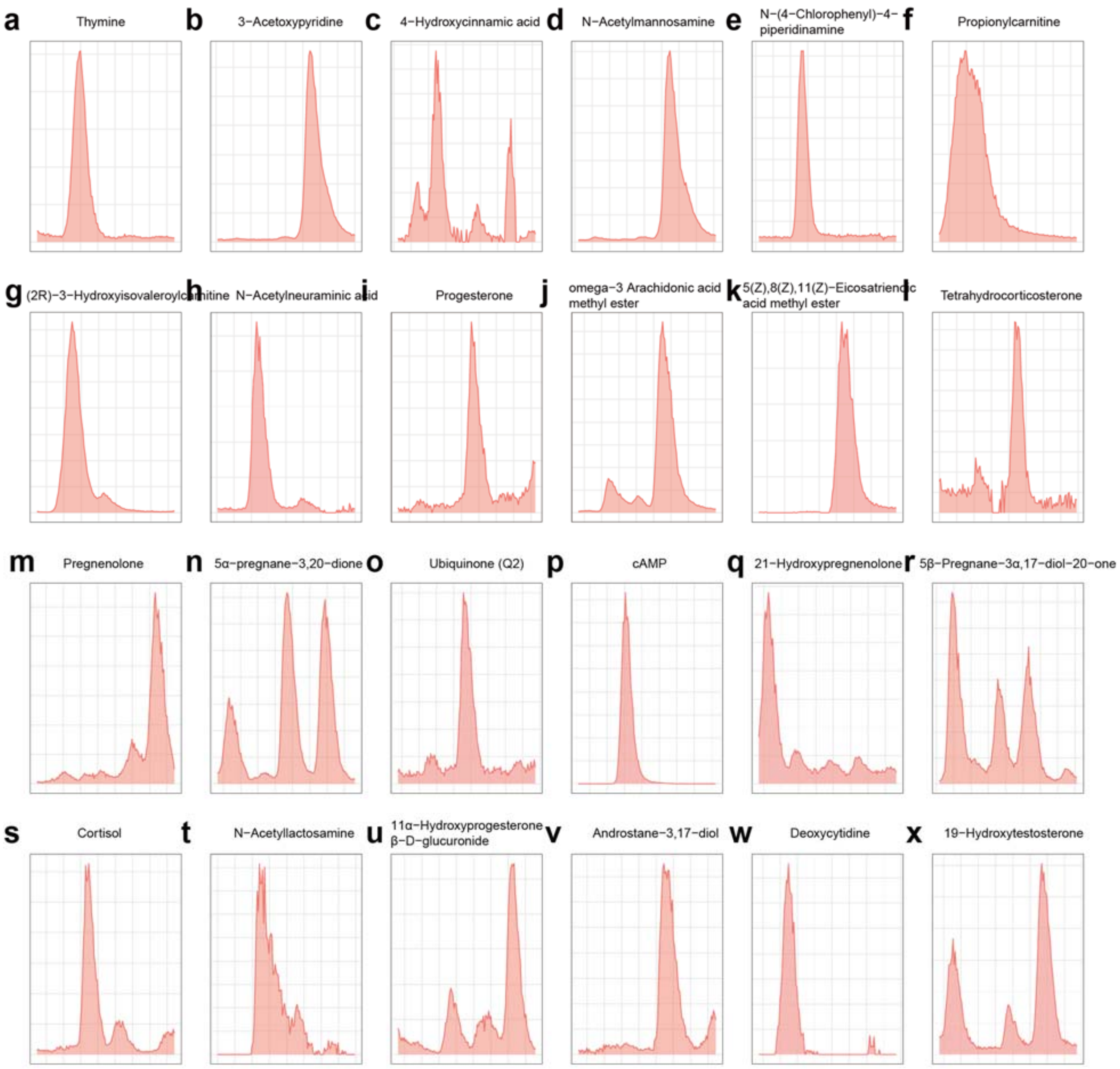


**Figure S8. The peak shapes of 24 metabolite biomarkers in GA and sampling time to delivery models.**

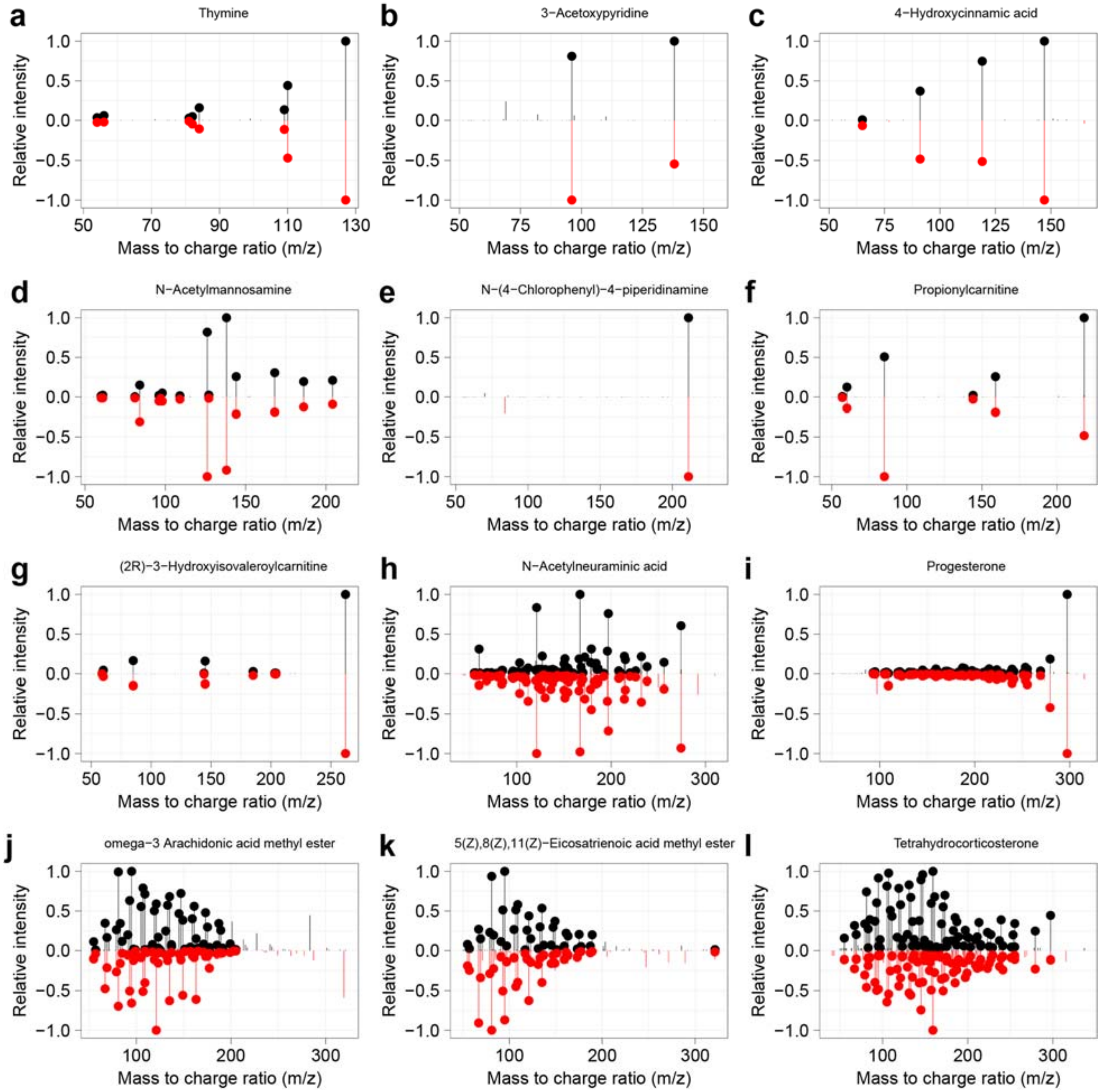

a
Thymine
b
3−Acetoxypyridine
c
4−Hydroxycinnamic acid
d
N−Acetylmannosamine
e
N−(4−Chlorophenyl)−4−piperidinamine
f
Propionylcarnitine
g
(2R)−3−Hydroxyisovaleroylcarnitine
h
N−Acetylneuraminic acid
i
Progesterone
j
omega−3 Arachidonic acid methyl ester
k
5(Z),8(Z),11(Z)−Eicosatrienoic acid methyl ester
l
Tetrahydrocorticosterone
Relative intensity
Mass to charge ratio (m/z)

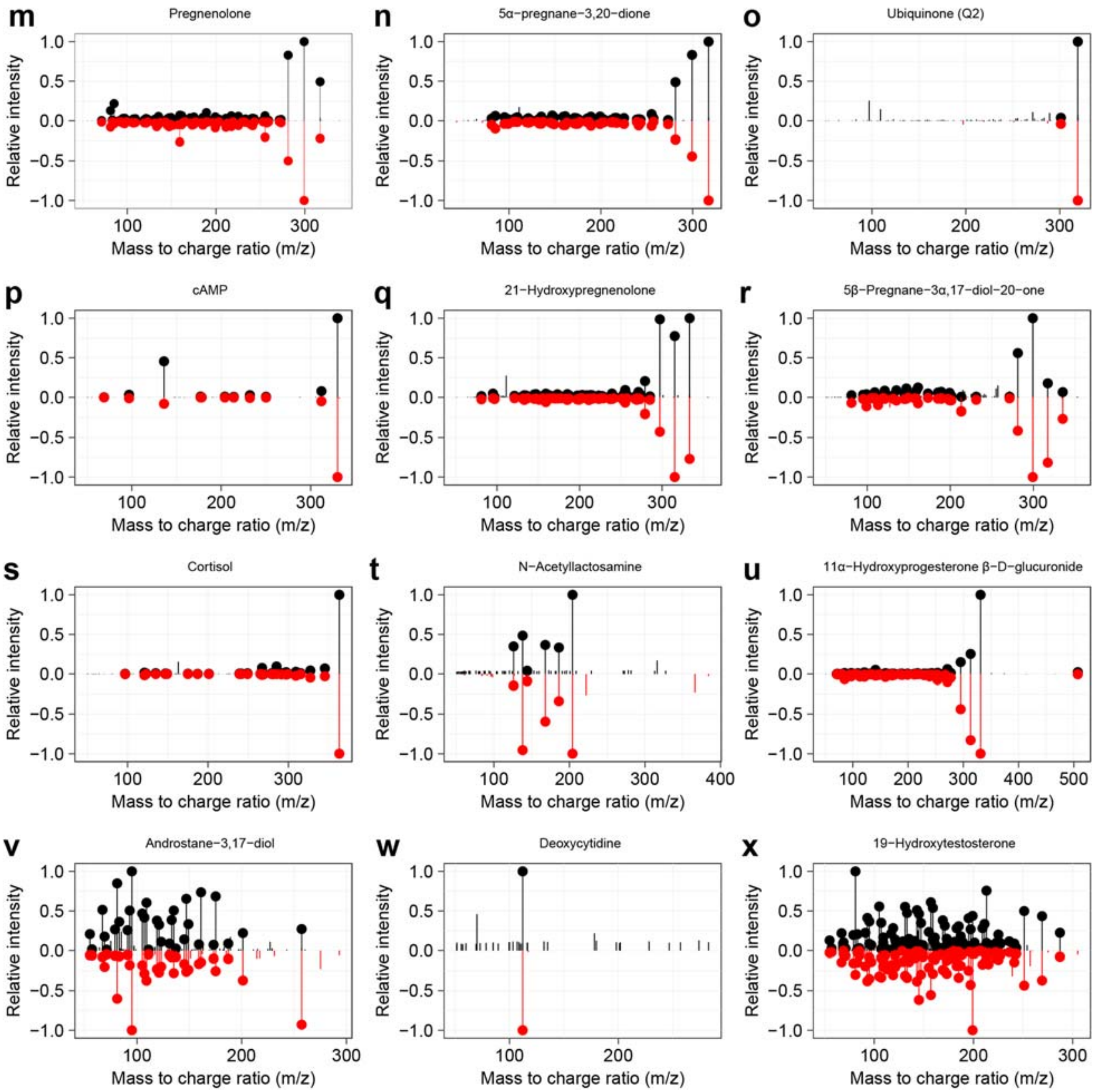


**Figure S9. The $MS^2$ spectra match of 24 metabolite biomarkers in gestational age and sampling time to delivery models.**

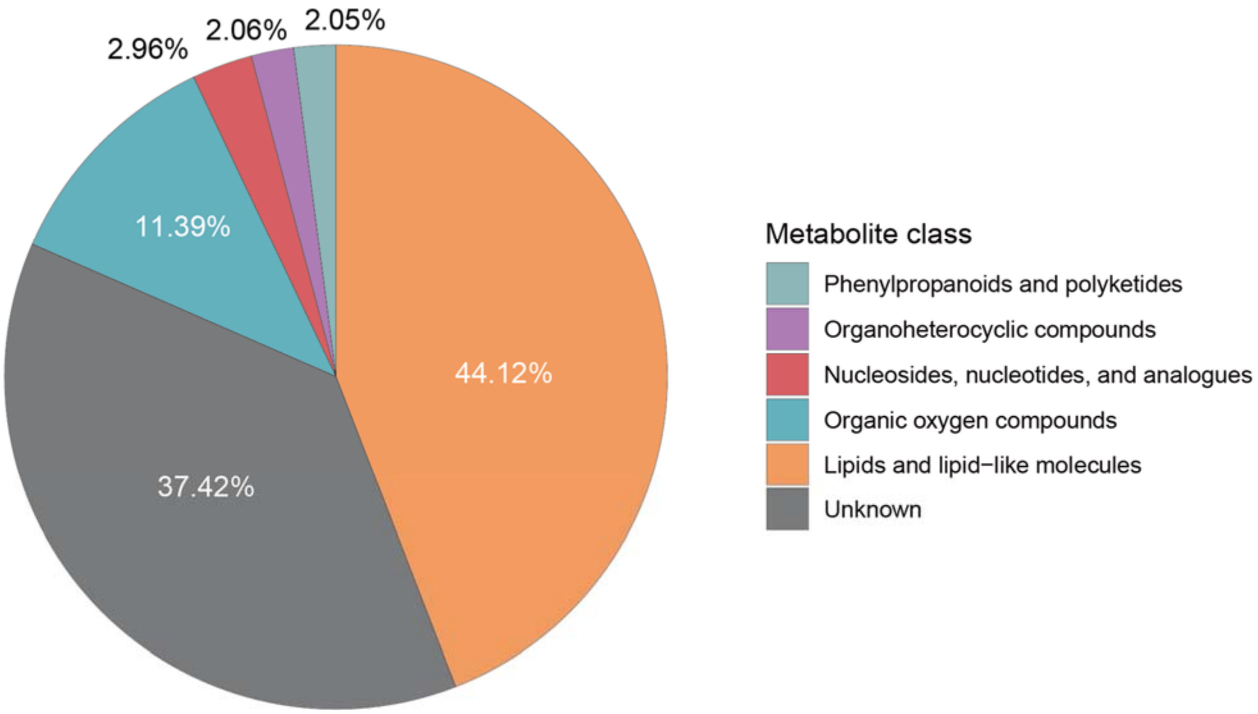


**Figure S10**. **Importance ratio of different chemical class in prediction model for gestation age.**

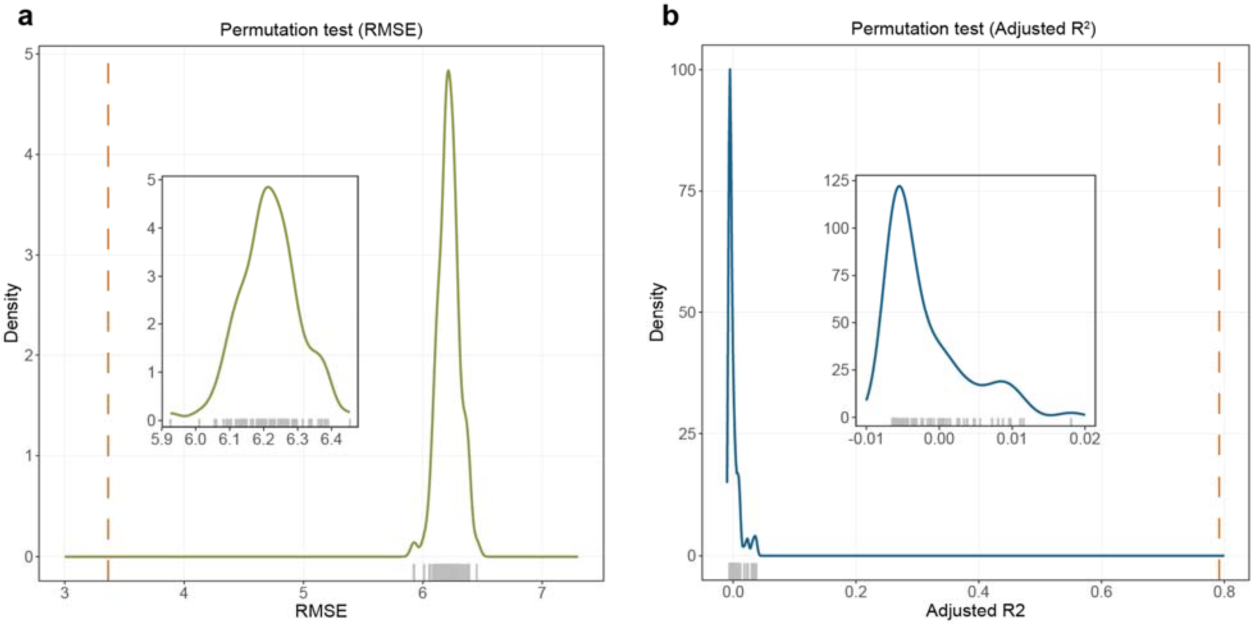


**Figure S11.** Permutation test for prediction module for gestational age. (**a**) Null distribution of RMSE values. (**b**) Null distribution of adjusted $R^2$ values.

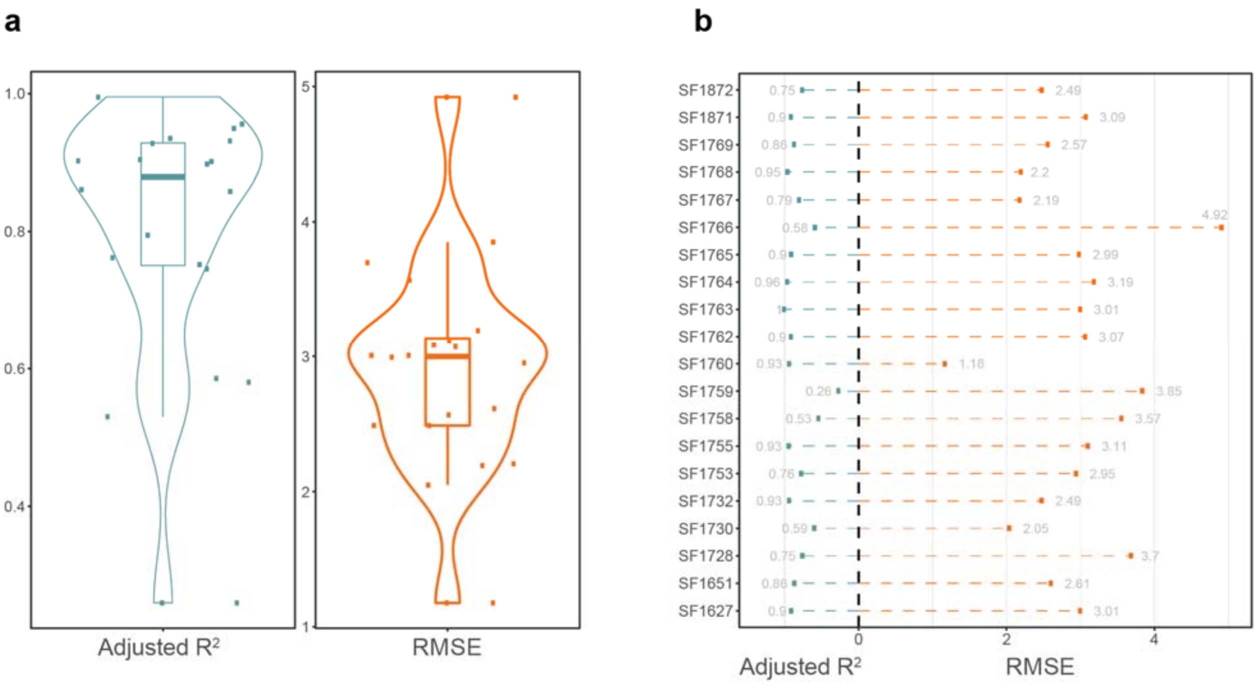


**Figure S12. Gestational age prediction result for each participant.** (**a**) Distribution of RMSE and adjusted $R^2$ for each participant in the validation dataset. (**b**) RMSE and adjusted $R^2$ for each participant in the validation dataset.

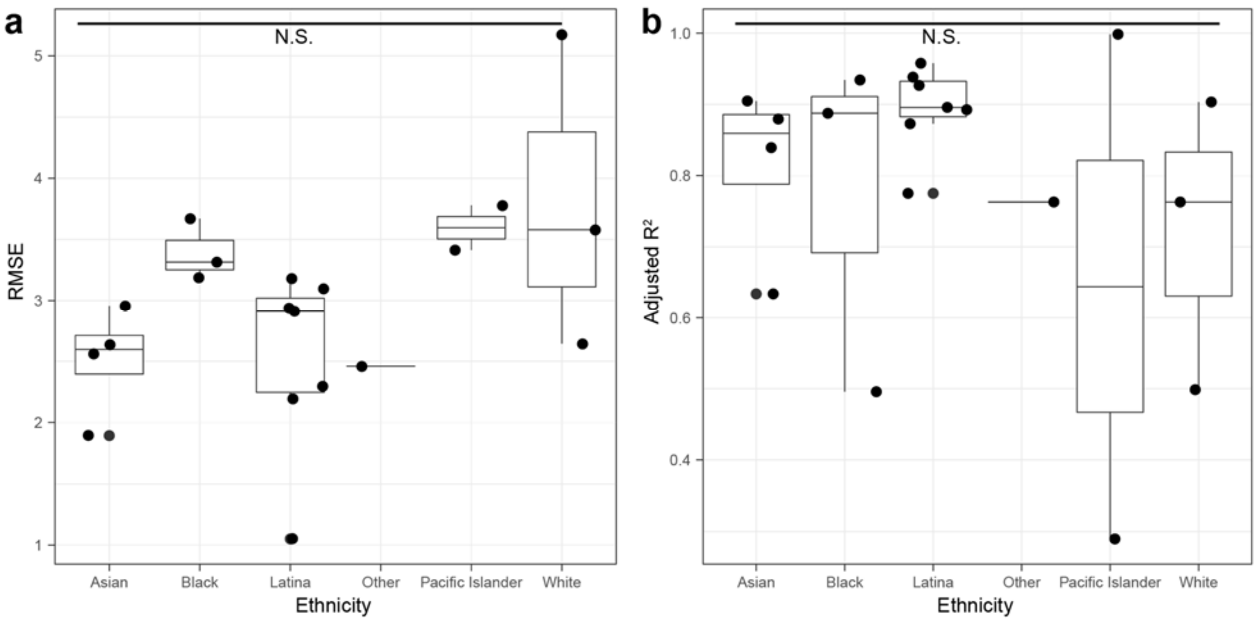


**Figure S13. Prediction accuracy with characteristics in GA model.** (**a**) RMSE in different ethnicities are not significant. (**b**) Adjusted R2 in different ethnicities are not significant.

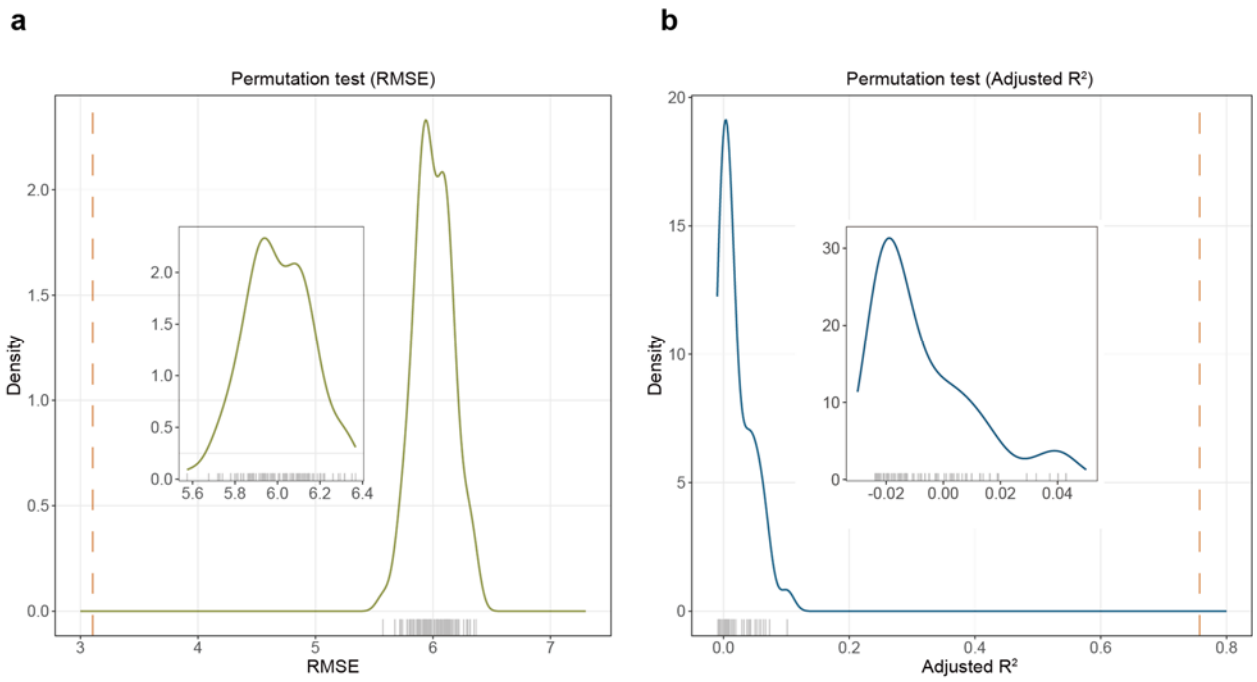


**Figure S14. Permutation test for prediction module for sampling time to delivery.** (**a**) Null distribution of RMSE values. (**b**) Null distribution of adjusted $R^2$ values.

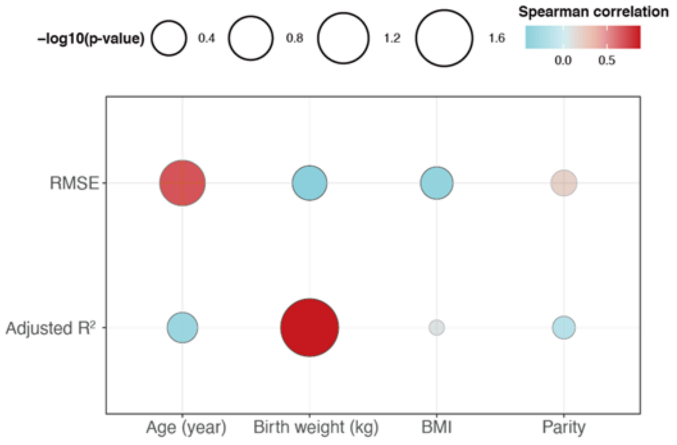


**Figure S15. The continuous characteristics have no effect on sampling to delivery prediction accuracy.**

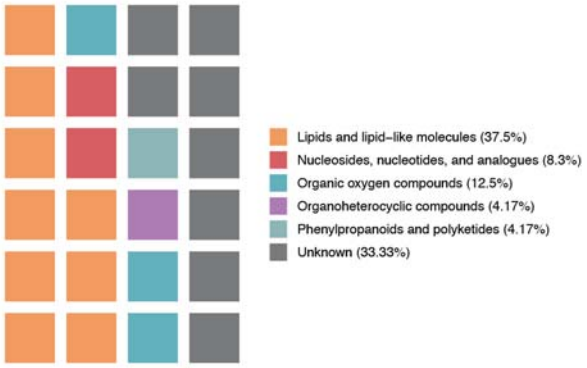


**Figure S16. Chemical class of 24 metabolite biomarkers in GA and sampling time to delivery models.**

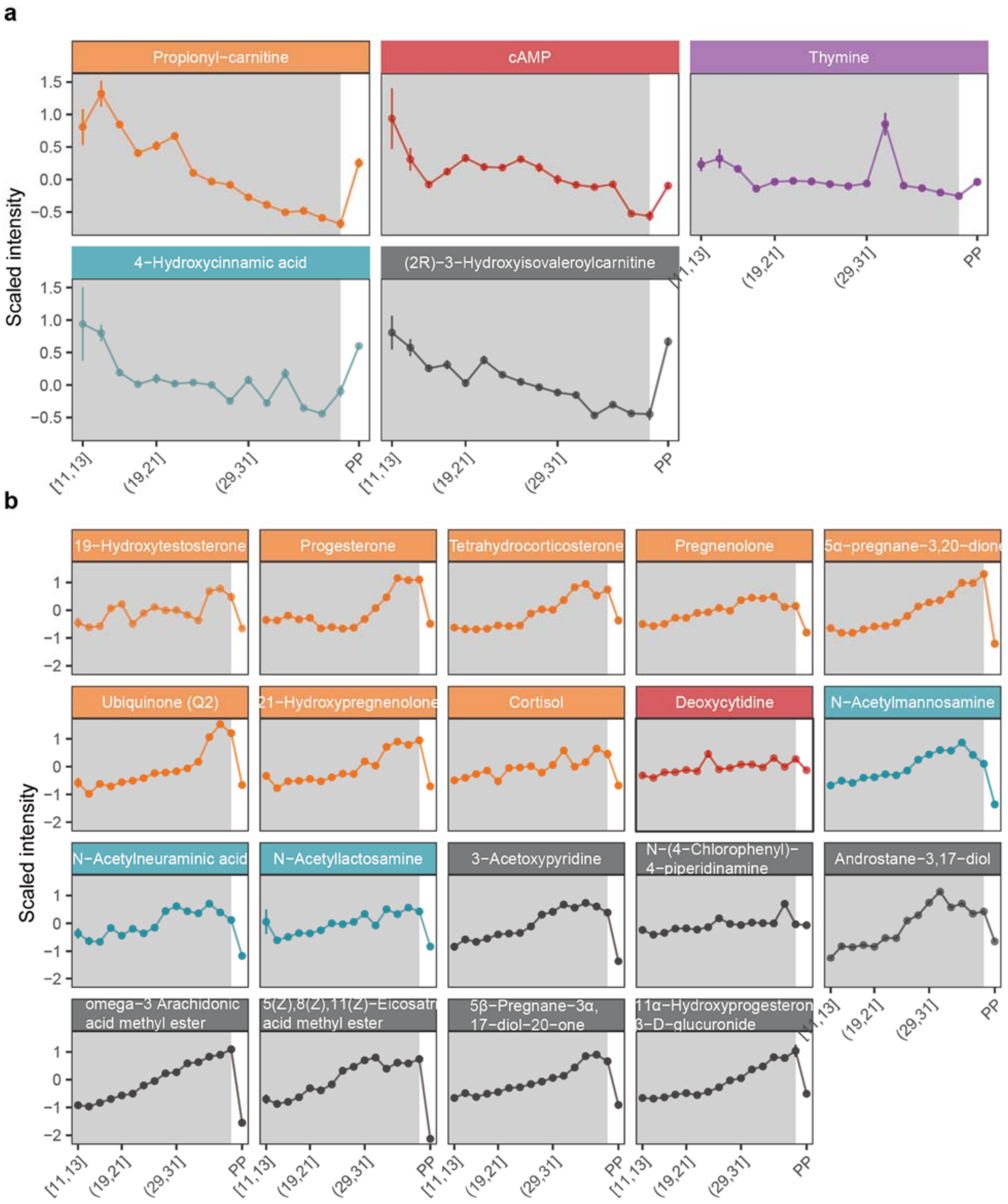


**Figure S17. The trends of 24 metabolite markers during pregnancy.** (**a**) Five metabolite markers decrease during pregnancy and increase after childbirth. (**b**) Nineteen metabolite markers increase during pregnancy and decrease after childbirth.

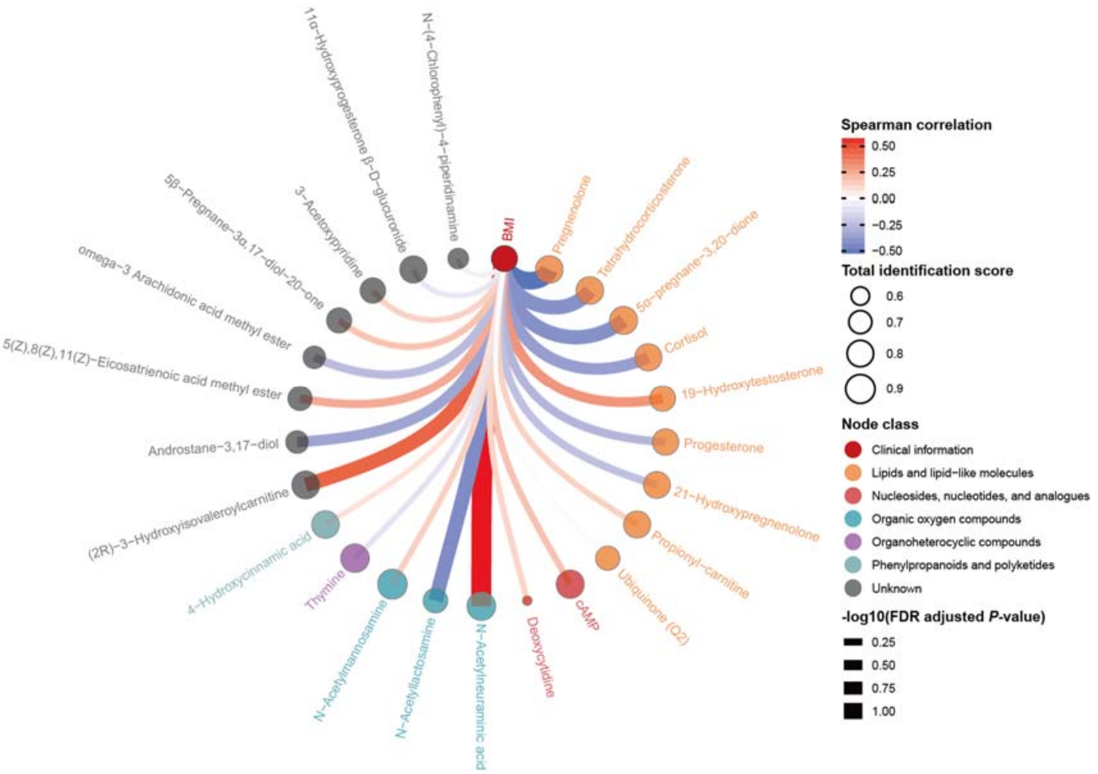


**Figure S18. Correlation network between BMI and metabolites markers.**

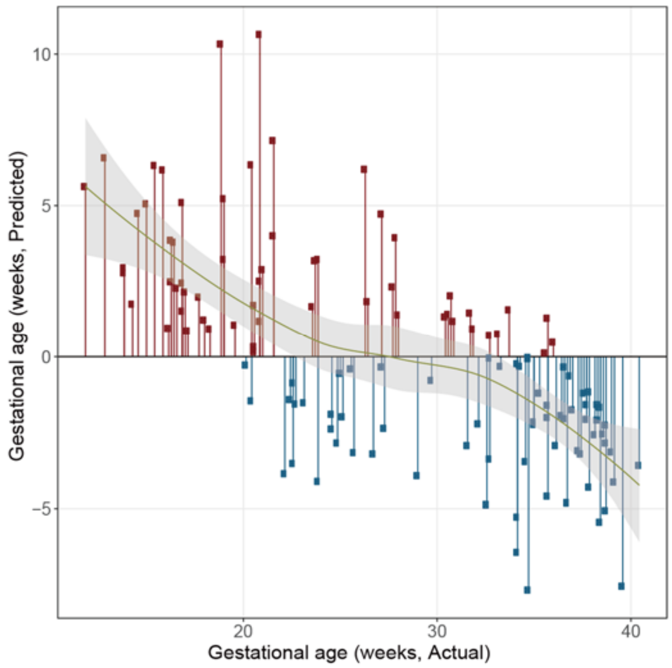


**Figure S19. Prediction error using Random Forest to predict gestational age.**
